# Cognitive network interactions through communication subspaces in large-scale models of the neocortex

**DOI:** 10.1101/2024.11.01.621513

**Authors:** Ulises Pereira-Obilinovic, Sean Froudist-Walsh, Xiao-Jing Wang

## Abstract

Neocortex-wide neural activity is organized into distinct networks of areas engaged in different cognitive processes. To elucidate the underlying mechanism of flexible network reconfiguration, we developed connectivity-constrained macaque and human whole-cortex models. In our model, within-area connectivity consists of a mixture of symmetric, asymmetric, and random motifs that give rise to stable (attractor) or transient (sequential) heterogeneous dynamics. Assuming sparse low-rank plus random inter-areal connectivity constrained by cognitive networks’ activation maps, we show that our model captures key aspects of the cognitive networks’ dynamics and interactions observed experimentally. In particular, the anti-correlation between the default mode network and the dorsal attention network. Communication between networks is shaped by the alignment of long-range communication subspaces with local connectivity motifs and is switchable in a bottom-up salience-dependent routing mechanism. Furthermore, the frontoparietal multiple-demand network displays a coexistence of stable and dynamic coding, suitable for top-down cognitive control. Our work provides a theoretical framework for understanding the dynamic routing in the cortical networks during cognition.

## 1 Introduction

During behavior, widespread sets of brain areas engage in different cognitive processes [1, 2, 3, 4, 5]. These sets of areas are sometimes referred to as cognitive networks [6, 7], as they are associated with distinct cognitive functions. For example, whether the focus of attention at any moment is on the external world or internally-generated thoughts depends on competition between the dorsal attention network (DAN) [8, 9, 10] and the default mode network (DMN) [11, 12, 13, 14, 15]. These two networks are each distributed in a non-contiguous arrangement with nodes across the parietal, temporal, and prefrontal cortex [16]. Two other commonly studied higher cognitive networks are the salience network, and the frontoparietal multiple-demand network. The salience network activates transiently to detect salient stimuli, and may bias competition between the DAN and DMN [17, 18] through an unknown mechanism. The frontoparietal network (also called the multiple demand network) [19, 20, 21, 22, 23], is engaged for cognitively demanding tasks and can flexibly couple with the DAN or DMN depending on the task at hand [22, 24, 25, 26], although the circuit mechanisms of how this flexible coupling is achieved are not yet understood. An important open question in cognitive neuroscience is: where are the switches [27] that enable the dynamic reconfiguration of cognitive networks according to task demands?

Recent multi-regional recordings in non-human primates and rodents suggest that long-range projections in the cortex act as dimensionality bottlenecks that restrict the maximum dimension of the routed signals across the cortex [28, 29, 30, 31]. These bottlenecks, referred to as communication subspaces, allow the selective routing of some dimensions of cortical activity while others remain private [28]. Depending on the alignment of the local cortical activity with the communication subspaces, signals can be flexibly routed, generating time-dependent functional subnetworks that correlate with behavior [31, 30]. This work opens the possibility that cognitive networks dynamically coordinate through communication subspaces in space and time.

Large-scale brain models that use anatomical data as structural constraints offer a theoretical approach to understanding the multiregional cortex [32]. Such connectome-based modeling has been developed for the resting state [33, 34, 35, 36, 37, 38, 39, 40], more recently for working memory [41, 42, 43, 44], decision-making [45, 46] and conscious access [47]. However, their local circuit dynamics are modeled with only a few variables, limiting their ability to capture communication of behaviorally selective activity in high-dimensional spaces defined by the spanned activity of a large number of neurons. By increasing the number of variables for modeling the neural dynamics of each area, one can enhance these models’ capacity to represent higher-dimensional dynamics, and explain how different types of information are represented and propagated along the cortex. Using these higher-dimensional representations, one could analyze how neural dynamics are routed through communication subspaces. However, increasing the number of variables can quickly lead to a loss of interpretability in these models [48].

To balance our ability to reproduce in neural network models the routing of high-dimensional neural dynamics through communication subspaces and maintain the model’s interpretability, we focus on building a class of connectivity-constrained multi-regional recurrent neural networks (MRNNs) that capture key qualitative features of population encoding of task variables. During behavior, population activity can be stable [49, 50, 51, 52, 53], albeit with sometimes strong temporal heterogeneities, or dynamic [54, 51, 52, 53, 55, 56] altering the encoding of task variables over the course of a trial. Areas where stable encoding has typically been observed, such as the parietal, secondary motor, and prefrontal cortex, can also display dynamic coding [57, 58, 59]. In our model, we hypothesize that cognitive network dynamics are shaped by the interplay between stable and dynamic encoding through attractor dynamics and sequential population activity, coordinated by communication subspaces, which facilitates the flexible interaction of cognitive networks in response to changing cognitive demands.

Based on these previous observations, we build both macaque and human whole-cortex MRNN models with a large number of neural pools (i.e., units) per area and analyze how local dynamics and inter-area communication organize cognitive networks. We model within-area connectivity as a mixture of symmetric [60, 61, 62], asymmetric [63, 64, 65], and random [66] motifs with systematic variation of their strength along the cortical hierarchy [67]. Communication subspaces are defined by modeling long-range projections as low-rank, sparse connectivity matrices [68], with their sparsity constrained by anatomical data [69, 36].

We find that specific alignment of local connectivity with the communication subspaces can explain the characteristic dynamic interactions of the DAN, DMN, FPN and salience network during cognition. Our model also demonstrates how, depending on the flexible alignment of local connectivity and inter-areal communications subspaces, stable and dynamic coding of cognitive variables can coexist in the frontoparietal network. Our work introduces a theoretical framework for understanding the dynamics and interactions of cognitive networks during cognition.

## 2 Results

### 2.1 Multi-regional recurrent neural network

We build and analyze a class of connectivity-constrained multi-regional recurrent neural network (MRNN) models to dissect how cognitive network dynamics are shaped by the interplay of attractor dynamics and sequential activity and coordinated through communication subspaces. In these models, each region is represented by a distinct local RNN with *N* units. Each unit represents a neural pool (or assembly) that is selective for specific features. Their local synaptic weight 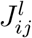 between a pre-synaptic unit *j* and a post-synaptic unit *i* in region *l* consists of a combination of symmetric (Fig. 1a), asymmetric (Fig. 1b), and random (Fig. 1c) connectivity motifs

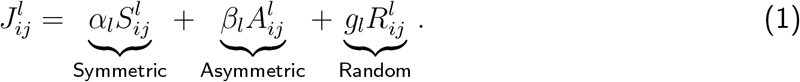

**Figure 1:**
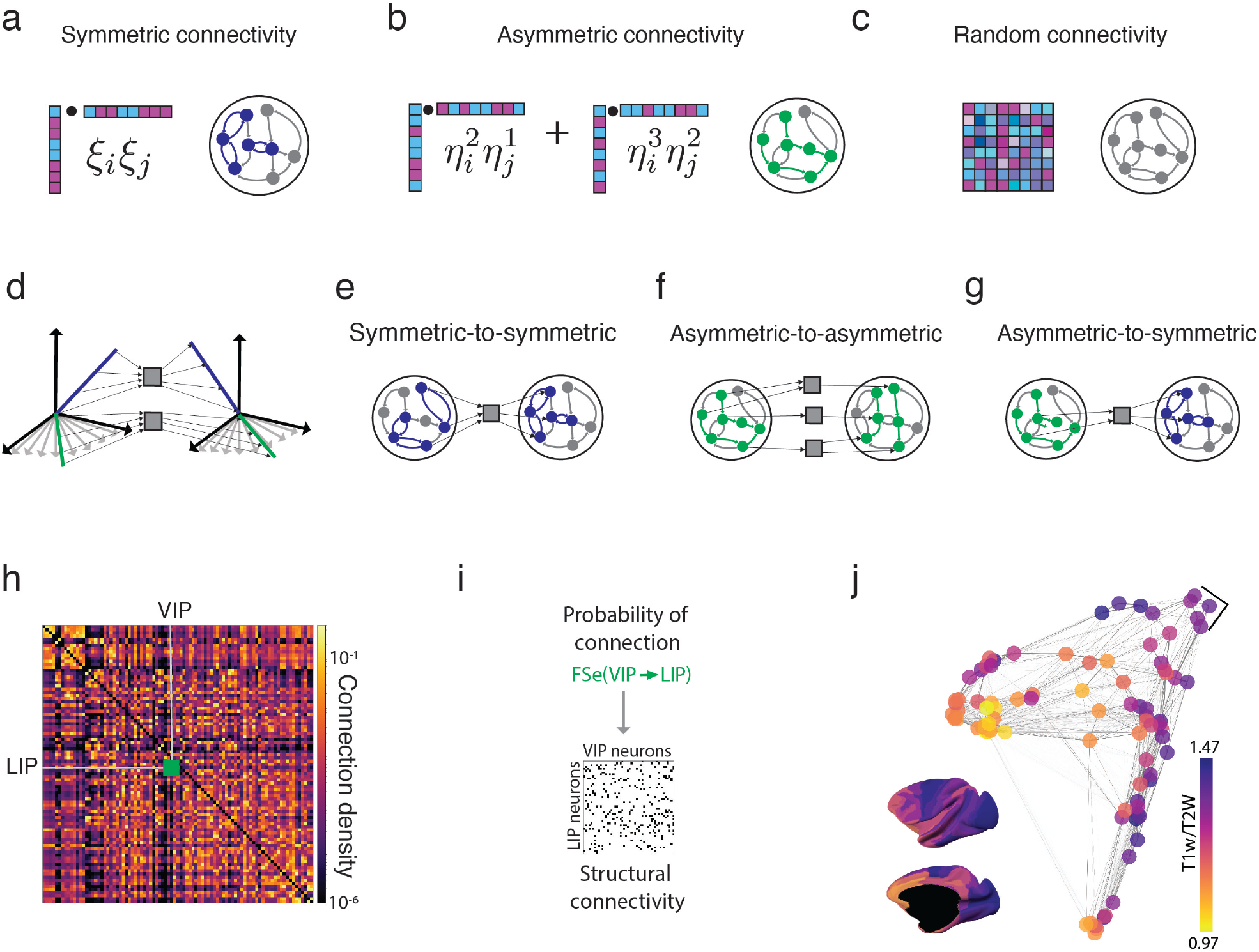
Local dynamics, communication subspaces, and anatomical constraints. (a-c) Schematics of a local RNN with symmetric (a) Eq.(7), asymmetric (b) Eq.(8), and random (c) Eq.(9) connectivity motifs. (d) Schematic of a rank-2 communication subspace that reads out activity patterns corresponding to the symmetric (blue) and asymmetric (green) connectivity motifs and projects these to activity patterns corresponding to the symmetric and asymmetric connectivity motifs, respectively. (e) Schematic of symmetric-to-symmetric rank-1 communication subspace connectivity between two areas. (f) Schematic of the same areas connecting via a asymmetric-to-asymmetric rank-3 communication subspace. (g) Schematic of the same areas connecting via an asymmetric-to-symmetric rank-1 communication subspace. (e-g) are schematics of sparse communication subspaces (Eq. (4)). (h) Connectivity matrix, deriving from dMRI tractography of the macaque cortex parcellated into 89 cortical regions [69]. (i) The sparsity of the projections across areas is proportional to the normalized entries of the tractography data (Methods 4.4, Eq.(4)). (j) The T1w/T2w measure across the macaque cortex and dMRI connectivity graph. The edges thicknesses are proportional to the weights in the connectivity matrix. The nodes are colored according to the T1w/T2w value. Nodes are projected into 2-d space using the diffusion map algorithm (Methods 4.2).

The parameters *α*_*l*_, *β*_*l*_, and *g*_*l*_ determine the strength of the symmetric, asymmetric, and random connectivity motifs, respectively. The symmetric connectivity is such that synaptic weights between any pair of neurons have the same strength in both directions, i.e., 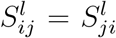 (Fig. 1a). As in network models for attractor dynamics [61, 62] the symmetric connectivity motif in our model is built as the sum of the outer product of *N*-dimensional patterns (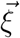in Fig. 1a, Methods 4.4.1, Eq. (7)). This type of symmetric connectivity generates strong recurrent loops (Fig. 1a), producing multi-stable attractor dynamics and stable persistent activity in recurrent networks [60, 61, 62, 70].

The asymmetric connectivity motif is such that 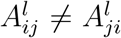 and has an effective feed-forward structure [63, 64, 71, 72, 73, 74, 65] given by the sequential outer product of *p*_A_ patterns (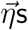in Fig. 1b, Methods 4.4.1, Eq. (8)). The asymmetric connectivity motif in Eq. (8) produces transient sequential population activity in recurrent networks [64, 63, 71, 65, 74]. Additionally, multiple sequences can be embedded in each local RNN’s connectivity by adding multiple independent sequences of patterns [65].

The random connectivity motif (Fig. 1c, see Methods 4.4.1, Eq. (9)) is known to produce high-dimensional chaos in RNNs characterized by strongly temporally heterogeneous dynamics [66].

The rationale for having such a mixture of connectivity motifs is that, depending on the local circuit’s parameters (i.e., *α*_*l*_, *β*_*l*_, and *g*_*l*_) and their inputs [75], local circuits can exhibit various qualitative types of dynamics, such as attractor dynamics, sequences, and chaos.

In our MRNNs, the synaptic weights of long-range projections that connect pre-synaptic unit *j* in area *l* with post-synaptic unit *i* in area *k* is given by two components

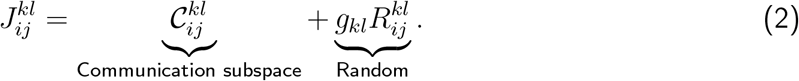

These projections are comprised of a structured component 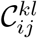, which we refer to as the communication subspace, and a random component 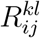, similar to the random connectivity in the local RNNs in Eq. (9). The effect of 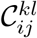 is reading out from a selected number of directions in a *N*-dimensional neural space defined by the spanned activity in the projecting area *l* and projecting to another small set of selected directions in the connected area *k*. Therefore, long-range projections can selectively route a small number of dimensions of neural dynamics while maintaining others private. The random component, 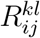,introduces variability that acts as quenched random interference in the communication subspace.

In our model, communication subspaces are constructed by reading out activity patterns and projecting to the directions of patterns 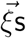 and 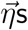 that comprise the symmetrically or asymmetrically connected ensembles of units in the local RNNs (Fig. 1d). For instance, symmetric-to-symmetric communication subspaces read out from one direction (i.e. a pattern comprising a particular symmetrically connected ensemble of units in the source area) and project in the direction of another pattern (corresponding to another symmetrically connected ensemble in the target area; Fig. 1e). Asymmetric-to-asymmetric (Fig. 1f) and asymmetric-to-symmetric (Fig. 1g) communication subspaces work similarly.

### 2.2 Anatomical constraints for whole-cortex models

We use diffusion-weighted Magnetic Resonance Imaging (dMRI) tractography data for the macaque [69] (Fig. 1h) and human [36] (Fig. S1a) cortices for constraining the long-range projections of our macaque and human whole-cortex MRNNs. In a departure from previous large-scale models, this anatomical data is used to construct the structural connectivity of the MRNNs, determining which units are connected across areas but not the values of the synaptic weights, which are determined by Eq. (2). Areas with small values in the weights of the dMRI tractography matrix indicate a small number of fibers connecting those brain regions, while larger values indicate denser projections. In our models, long-range connections between cortical areas are sparse. The sparsity of the projections is modeled with a sparse random (i.e., Erdős-Rényi) structural connectivity matrix, 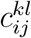, with average sparsity given by the normalized dMRI tractography matrix F_*kl*_ (i.e., 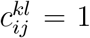 with probability 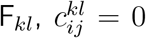 with probability 1 − F_*kl*_, Methods 4.4, Eq. (4)). Therefore, for each inter-areal connection, we first construct the (fully connected) inter-areal synaptic weights according to Eq. (2), and then apply a sparse random mask constructed based on the anatomical connectivity. The resulting long-range projections are a sparse version of the long-range connectivity matrix in Eq. (2) (Fig. 1h and i; Methods 4.4, Eqs. (3,4)).

Additionally, our model assumes linear differences in the strength of local and long-range connectivity across areas (Methods 4.4, Eqs. (3,5)). This assumption is based on the strong correlation observed between the spine count per neuron, a proxy for the strength of synaptic excitation, and the hierarchical position of cortical areas [67]. We use MRI-derived T1-weighted/T2-weighted (T1w/T2w) neuroimaging value to approximate the hierarchical position of each cortical region (Figs. 1j, S1b; Methods 4.1.2 and 4.1.4). The T1w/T2w values negatively correlate with the spine count [35, 36]. Lastly, the MRNN dynamics are given by the standard rate equations [76, 61] (Eq. (3)).

### 2.3 Stable population coding and heterogeneous neural dynamics in the frontoparietal network

The frontoparietal network is a large-scale brain network primarily involved in higher-order cognitive functions, including executive control, decision-making, attention, and working memory. This network spans the frontal cortex and the parietal cortex.

To define the frontoparietal network in the monkey cortex, we used the results of cross-species functional alignment [77] of the canonical maps by Yeo and colleagues, as previously [16, 78]. We defined areas as belonging to the frontoparietal network based on overlap (Methods 4.3.2) and added lateral intraparietal cortex (area LIP) since it is a key node in the working memory and decision-making system [79] (Fig. 2a; Table 1). Most of these areas present elevated persistent activity during working memory tasks [2].

**Table 1:**
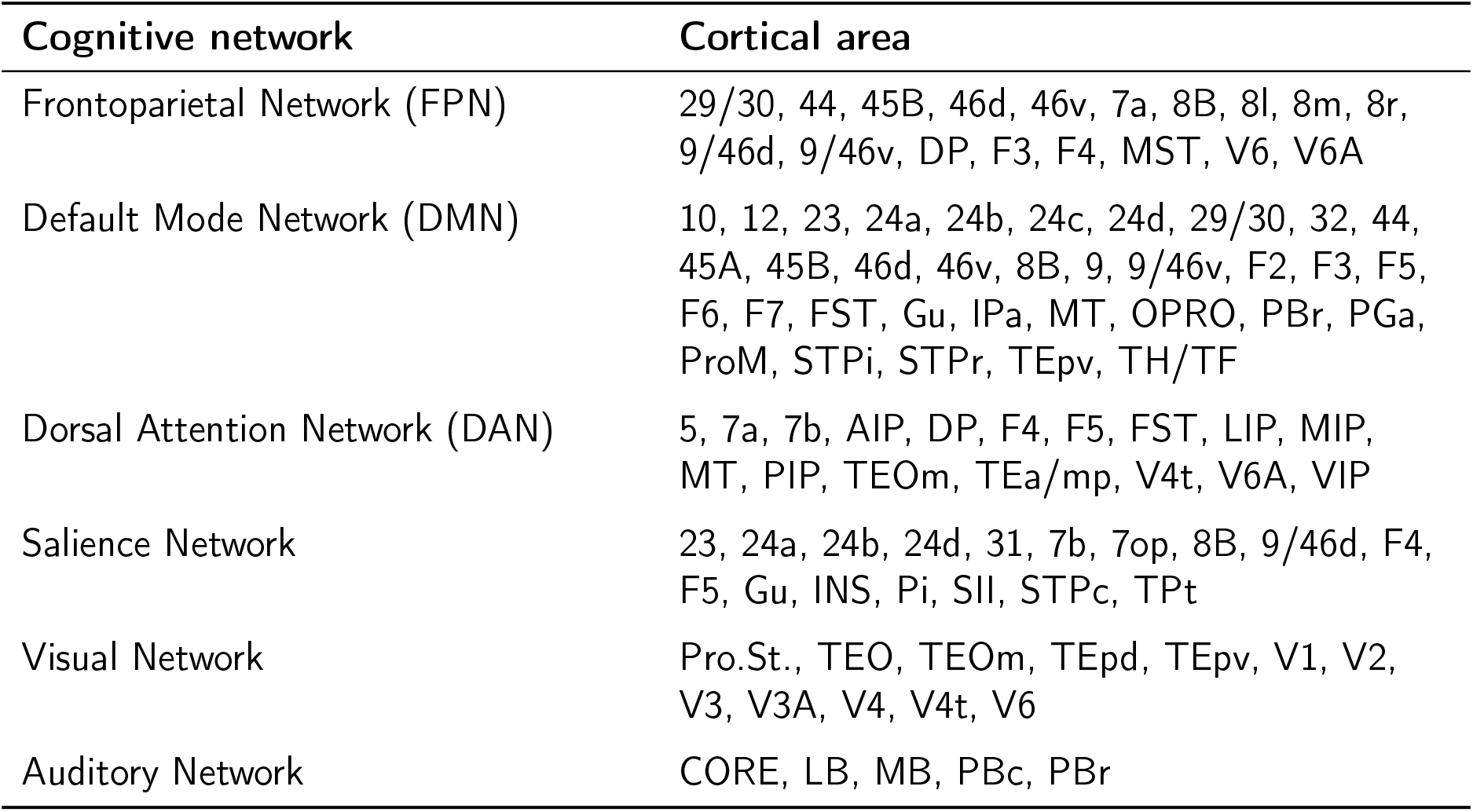
Functionally aligned cognitive networks and corresponding cortical areas for the macaque cortex.

| Cognitive network | Cortical area |
| --- | --- |
| Frontoparietal Network (FPN) | 29/30, 44, 45B, 46d, 46v, 7a, 8B, 8l, 8m, 8r, 9/46d, 9/46v, DP, F3, F4, MST, V6, V6A |
| Default Mode Network (DMN) | 10, 12, 23, 24a, 24b, 24c, 24d, 29/30, 32, 44, 45A, 45B, 46d, 46v, 8B, 9, 9/46v, F2, F3, F5, F6, F7, FST, Gu, IPa, MT, OPRO, PBr, PGa, ProM, STPi, STPr, TEpv, TH/TF |
| Dorsal Attention Network (DAN) | 5, 7a, 7b, AIP, DP, F4, F5, FST, LIP, MIP, MT, PIP, TEOm, TEa/mp, V4t, V6A, VIP |
| Salience Network | 23, 24a, 24b, 24d, 31, 7b, 7op, 8B, 9/46d, F4, F5, Gu, INS, Pi, SII, STPc, TPt |
| Visual Network | Pro.St., TEO, TEOm, TEpd, TEpv, V1, V2, V3, V3A, V4, V4t, V6 |
| Auditory Network | CORE, LB, MB, PBc, PBr |

**Figure 2:**
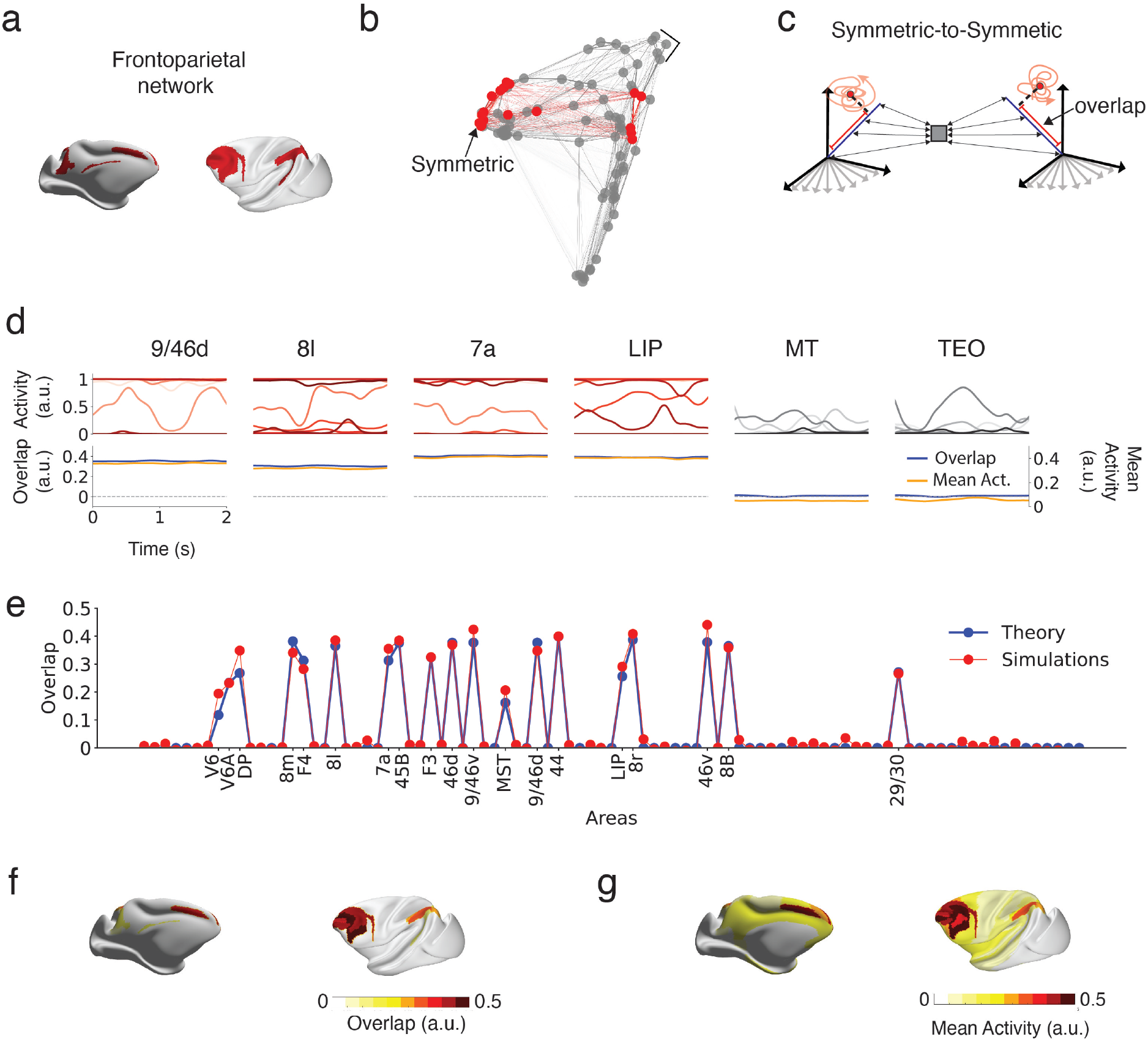
Distributed stable population coding through communication subspaces coexists with heterogeneous neural dynamics in the frontoparietal network. (a) Frontoparietal network areas. (b) dMRI connectivity graph. The edges are proportional to the connectivity strength from the dMRI tractography matrix. The nodes are placed using the diffusion map algorithm (Methods 4.2). Frontoparietal areas are connected by symmetric-to-symmetric long-range and local projections (red). (c) Schematics of a symmetric-to-symmetric communication subspace (rank-1) in frontoparietal areas. The activity within each frontoparietal area corresponds to a chaotic attractor with stable mean population activity aligned to the communication subspace that recurrently connects only frontoparietal areas. The overlaps with the patterns comprising the local symmetric connectivity motifs are indicated in red. (d) Neural dynamics for 15 representative units (top), overlaps (bottom, left axis, blue), and mean activity (bottom, right axis, yellow) for six representative areas. The horizontal dashed line at the bottom indicates zero overlap. The areas with red traces for the activity, 9/46d (prefrontal cortex), 8l (prefrontal cortex), 7a (parietal cortex), and LIP (parietal cortex), belong to the frontoparietal network, while MT (higher visual cortex) and TEO (temporal cortex) do not. (e) Average overlap over 4.5s vs. prediction from the mean field theory (Methods 4.5). (f) Overlaps at 2s. (g) Mean activity at 2s.

For modeling the selective persistent activity observed in areas in the frontoparietal network, we first include symmetric local connectivity only in those areas (Methods 4.4.3). The rationale is that, as in attractor neural networks [61, 80], symmetrically connected ensembles with strong enough connectivity will lead to discrete attractor dynamics that selectively elevate persistent activity in a subset of neurons in each area within the frontoparietal network. However, in our model, the local symmetric connectivity is weak, and none of the areas in the frontoparietal network can sustain attractor dynamics on their own. Areas within the frontoparietal network are connected by long-range symmetric-to-symmetric connectivity subspaces (Fig. 2e), in which units locally connected by symmetric connectivity are projected to symmetrically connected units in the projected area (Methods 4.4.3). These long-range symmetric-to-symmetric connectivity subspaces are recurrent, connecting all areas within the frontoparietal network (Fig. 2b).

For each area, we compute the projection of the patterns comprising the symmetric and asymmetric connectivity motifs onto the population vector for that area (see Methods 4.5). We refer to these projections as the overlap, as they measure how aligned the population activity of each area in the MRNNs is with the patterns comprising the local symmetric or asymmetric connectivity motifs (Methods 4.5; Eq. (39)). The overlaps serve as a proxy for how linearly decodable the specific pattern, from which the overlap was computed, is and provide a measure of the extent to which the symmetric and asymmetric connectivity motifs are driving the local RNN dynamics.

When areas in the frontoparietal network are stimulated, the network converges to a state in which units in most areas present strong fluctuations in their activity (Fig.2d top). Multiple areas within the frontoparietal network present stable non-zero overlaps with the pattern corresponding to the local symmetric connectivity motif (Fig. 2d, see bottom). Although the population activity within the frontoparietal network is aligned with the symmetric-to-symmetric communication subspace and is stable in time, the activity of single units strongly fluctuates. These fluctuations are a signature of chaotic dynamics with stable encoding properties, which have been previously characterized in local circuit models as attractor neural networks [81, 82] and low-rank networks [83]. Therefore, our multi-regional network converges to a chaotic attractor in which the population activity within the frontoparietal network is sustained through communication subspaces through which symmetrically connected cortical ensembles interact across areas. We developed a mean field theory [60, 81] that predicts the overlap values (Fig. 2e; Methods 4.5), which are high only in areas within the frontoparietal network (Fig. 2e and f). Units in areas that do not belong to the frontoparietal network also display chaotic fluctuations, but their activity is not aligned with any communication subspace (Figs. S2 and S3), and the mean activity of those areas is generally lower (Figs. 2g, S2, and S3). There is a gradient of mean activity across areas with predominantly higher mean activity in the frontoparietal network (Figs. 2g, S2, and S3).

### 2.4 Dynamic and stable population coding coexist in the frontoparietal network

During certain working memory and decision-making task conditions, population coding of memoranda is dynamic rather than stable [54, 51, 52, 53, 55]. Recent findings indicate that dynamic coding occurs in the parietal cortex [57] and motor cortex [58] during decision-making tasks, as well as in the prefrontal cortex during navigation working memory tasks [59]. Can a distributed dynamic coding emerge in our model and coexist with stable encoding?

For modeling the dynamic coding in the frontoparietal network, we incorporated asymmetric local connectivity in addition to the previously described local symmetric connectivity (Methods 4.4.4). Also, in addition to the symmetric-to-symmetric connectivity subspaces described in the previous section, areas within the frontoparietal network are connected by long-range asymmetric-to-asymmetric connectivity subspaces (Fig. 3b). In these connectivity subspaces, ensembles of units locally connected by asymmetric connectivity project to asymmetrically connected ensembles in the target area (Methods 4.4.4, Eq. (17, 18)). The dimension (i.e., rank) of these asymmetric connectivity subspaces is equal to the number of patterns that comprise the local asymmetric connectivity in the local circuits minus one. Importantly, they read out the population activity of each area in the frontoparietal network when it aligns with a pattern in the sequence, and then project the resulting current in the direction of the next pattern in the sequence in the connected area (Fig. 3b). Similar to symmetric-to-symmetric connectivity subspaces, these long-range asymmetric-to-asymmetric connectivity subspaces are recurrently arranged across areas (Methods 4.4.4, Eq. (17)), connecting all areas within the frontoparietal network (Fig. 3b).

**Figure 3:**
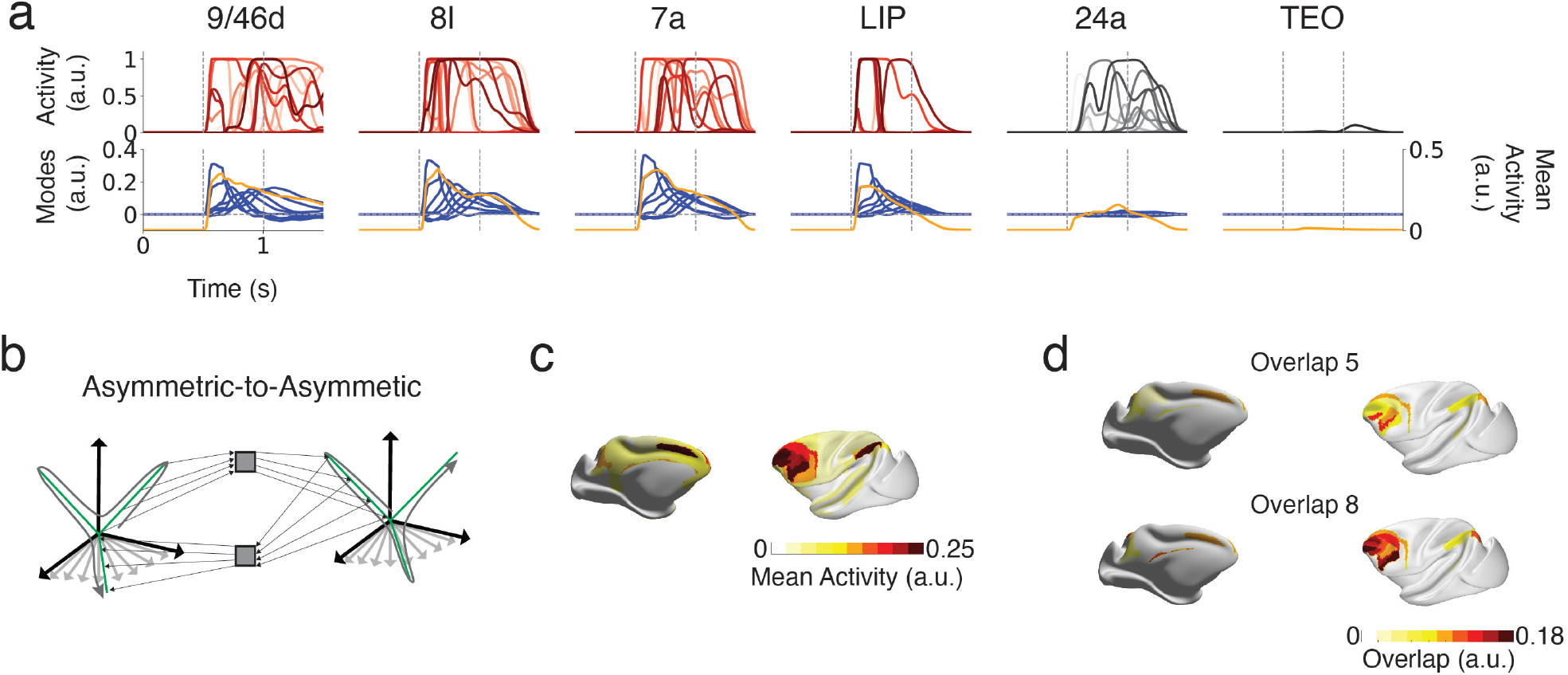
Distributed dynamic population coding through communication sub-spaces in the frontoparietal network. (a) Neural dynamics for 15 representative units (top), overlaps (bottom left axis, blue), and mean acitivity (bottom right axis, yellow) for six representative areas. The areas with red traces for the activity, 9/46d (prefrontal cortex), 8l (prefrontal cortex), 7a (parietal cortex), and LIP (parietal cortex), belong to the frontoparietal network, while 24a (cingulate cortex) and TEO (temporal cortex) do not (gray traces). The leftmost gray dashed line represents the moment when all frontoparietal areas are transiently stimulated with the first pattern in the sequence at the 0.5s. The rightmost vertical dashed line indicates 1s. The gray horizontal dashed line at the bottom indicates zero overlap. (b) Schematics of an asymmetric-to-asymmetric communication subspace in frontoparietal areas. When the population activity of each area in the frontoparietal network is transiently aligned with a given pattern that comprises the asymmetrical local connectivity indicated by the dashed line, it projects in the direction of the next pattern in the sequence in the connected area. (c) Mean activity at 1s. (d) Value of the 5th and 8th overlap in the sequence at 1s. The quantities in (c) and (d) are taken at 1s in the network simulation.

We stimulate all areas in the frontoparietal network with input current of 150ms of duration aligned with the first pattern in the sequence. A burst of heterogeneous activity spreads across cortical areas (Fig. 3a,c; Fig. S4). Only the population activity in areas within the frontoparietal network shows sequential overlaps (Fig. 3a bottom, d; Fig. S5), thus transiently aligning with the asymmetric-to-asymmetric communication subspace that recurrently connects these areas. These regions exhibit sequential population activity (Fig. 3a, bottom, d; Fig. S5). The asymmetric-to-asymmetric communication subspaces drive distributed sequential dynamics by connecting locally asymmetrically connected assemblies in brain regions across the frontoparietal network. The mean activity peaks within the frontoparietal network but elevated activity is observed across the cortex, including in regions that do not represent the sequence information (Fig. 3a bottom, orange trace; Fig. 3c).

When we stimulate the frontoparietal network with stimulus aligned with the symmetric-to-symmetric communication subspace, the network converges to a state where units in most areas exhibit strong fluctuations in their activity, but stable coding persists within the frontoparietal network, as previously shown in Fig. 3 (Figs. S6 and S7). Therefore, both distributed dynamic and stable coding can be observed in the frontoparietal network in our model, depending on the stimuli.

### 2.5 Frontoparietal attractor dynamics coordinates DMN-DAN interplay

During behavior, humans and non-human primates switch from unengaged to goal-directed behaviors. The default mode network (DMN), first characterized in humans [11] and later described in non-human primates [12, 13, 14, 15], is a large-scale cognitive network that is active when subjects are not focused on any task or the outside world [84]. In contrast, the dorsal attention network (DAN) is a set of areas activated when subjects engage in tasks requiring sustained externally directed attention and goal-directed behaviors [8, 9, 10]. A key observation is that both networks show significant anti-correlation across a wide array of cognitive tasks and unconstrained rest [85, 9]. Neuroimaging experiments suggest that the frontoparietal network may flexibly control the engagement of the DMN and DAN during behavior by coupling with both cognitive networks [22]. What is the mechanism for the competition between the DMN and DAN and the control by the frontoparietal network? Here, we propose a possible mechanism based on attractor dynamics mediated by communication subspaces.

We aligned the frontoparietal, DMN and DAN activation map in humans to the macaque cortex using cross-species functional alignment [77](Fig.4a-d). We define areas in both networks as regions in the macaque cortex that exhibit an overlap with the network map exceeding a specific threshold (Methods 4.3.2; Table 1).

To model the competition between the DMN and DAN and the control by the frontoparietal network, we first connect areas in the frontoparietal network through symmetric local connectivity. This connectivity consists of two symmetric connectivity components that define two distinct but partially overlapping ensembles of units (Methods 4.4.5). One of these ensembles in the frontoparietal network connects through long-range symmetric-to-symmetric communication sub-spaces with areas in the DMN. We will refer to these as the FP-DMN ensemble and the corresponding overlap as the FP-DMN overlap. The other ensemble in the frontoparietal network connects through a different long-range symmetric-to-symmetric communication sub-space with the DAN. We will refer to this as the FP-DAN ensemble with the FP-DAN corresponding overlap. Areas in both the DMN and the DAN have local connectivity consisting of only one component of symmetric connectivity and only one long-range symmetric-to-symmetric communication sub-space that recurrently connects these areas (Fig. 1e and Fig. 4d,e). These FP-DMN and FP-DAN ensembles of units in the frontoparietal network are connected by distinct long-range symmetric-to-symmetric communication sub-spaces, generating two distinct distributed attractor states (Fig.1e and Fig. 4e). One attractor encompasses the frontoparietal network and DMN (FP-DMN attractor), while the other encompasses the frontoparietal network and DAN (FP-DAN attractor).

**Figure 4:**
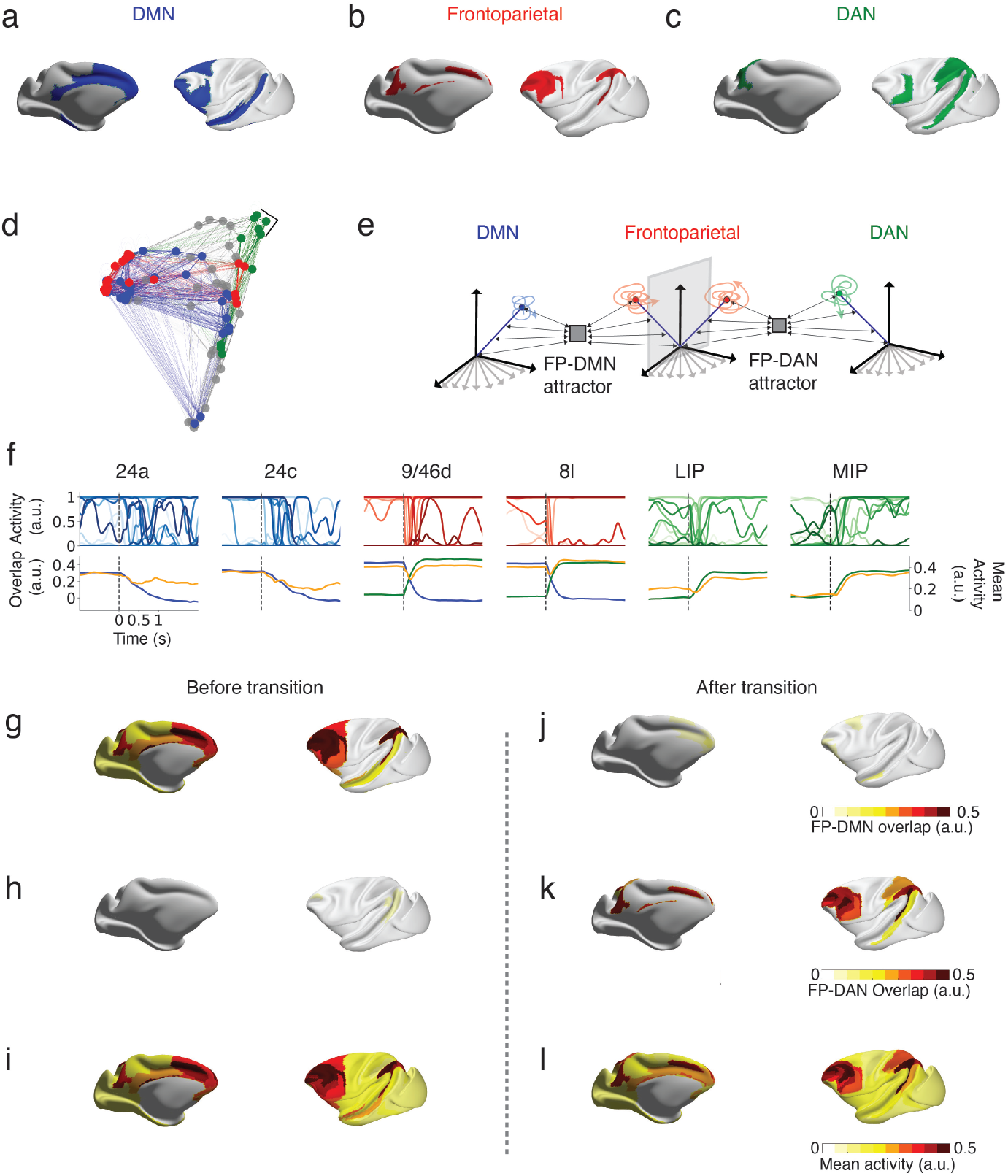
Frontoparietal control of default mode and dorsal attention networks through communication sub-spaces. (a) Default mode network (DMN). (b) Frontoparietal network (FP). (c) Dorsal attention network (DAN). (d) dMRI connectivity graph in diffusion map space (Methods 4.2). The edges and nodes that belong to the frontoparietal, default mode, and dorsal attention networks are colored blue, red, and green, respectively. (d) Schematics of two distinct attractors in the frontoparietal network. These attractors within the frontoparietal network interact separately with the default mode and dorsal attention networks through two distinct symmetric-to-symmetric communication subspaces. One attractor forms a distributed state, engaging both the frontoparietal and default mode networks (FP-DMN attractor), while the other engages the frontoparietal and dorsal attention networks (FP-DAN attractor). (f) Neural dynamics for 15 representative units (top), overlap (bottom left axis; blue: FP-DMN attractor; green: FP-DAN attractor), and mean activity (bottom right axis, yellow) for six representative areas. The first and second columns correspond to default mode areas 24a and 24c (activity traces in blue). The third and fourth columns correspond to frontoparietal areas 9/46d and 8l (activity traces in red). The fifth and sixth columns correspond to the dorsal attention network areas LIP and MIP (activity traces in green). The gray vertical dashed line represents the moment when an external constant stimulus switches its alignment from the FP-DMN to the FP-DAN attractor. (g and j) Overlap corresponding to the FP-DMN attractor. (h and k) Overlap corresponding to the FP-DAN attractor. (i and l) Mean activity. (g, h, i) Before the stimulus switch. (j, k, l) After the stimulus switch.

To investigate the transition between the FP-DMN and FP-DAN attractor states, we first introduced a constant input to the frontoparietal network areas. Initially, the input was aligned with the FP-DMN ensemble in the frontoparietal network. Despite the network exhibiting strong temporal variability due to chaotic dynamics (Fig. 4f and Fig. S8), the population activity overlap with the FP-DMN remained stable and elevated in the frontoparietal and DMN areas, while the FP-DAN overlaps were consistently low across all cortical areas (Fig. 4g-h; Figs. S8,S9). Mean activity peaked in the frontoparietal and DMN networks, but several areas across the cortex showed elevated mean activity, without encoding the FP-DMN overlap pattern (Fig. 4i and Fig. S8). When the input to the frontoparietal network shifted its alignment from the FP-DMN to the FP-DAN ensemble (dashed line in Fig. 4f), the FP-DMN overlap decreased in the frontoparietal and DMN areas, while the FP-DAN overlap increased (dashed line in Fig. 4f, j, k; Fig. S9). Consequently, the mean activity decreased in the DMN areas and increased in the DAN areas post-transition.

Therefore, our model recapitulates the anti-correlation in the mean activity between the DMN and the DAN, mediated by the frontoparietal network.

### 2.6 Dynamic routing of sensory stimuli via salience-dependent communication subspaces

Although a plethora of sensory stimuli are constantly present available to the senses, the brain prioritizes processing of only most salient. What is the mechanism for filtering out non-salient stimuli and selecting salient ones? We propose that communication subspaces can route salient sensory stimuli from sensory cortices to cognitive networks involved in flexible behavior.

To demonstrate this in our model, we study the interactions between visual and auditory cortices (Fig.5a-c) with cortical regions involved in filtering stimuli and recruiting relevant cognitive networks for executive control. These areas include the cingulate and insular cortices and are broadly defined as the salience network in humans [17, 18]. As previously done with the frontoparietal network, we aligned the activation map of the salience network in humans to the macaque cortex using cross-species functional alignment (Fig.5a and d). We define areas in the salience network as regions in the macaque cortex that exhibit an overlap exceeding a specific threshold (Methods 4.3; Table 1).

In our model, local connectivity in the visual, auditory, and salience networks is asymmetric and composed of three patterns to model transient network activation due to sensory stimuli lasting about 300-400ms. Areas within the visual, auditory, and salience networks are connected through recurrent asymmetric-to-asymmetric communication sub-spaces. To model the interactions between sensory cortices and the salience network, we also recurrently connected the visual and auditory networks with the salience network separately through asymmetric-to-asymmetric communication sub-spaces to two different sets of asymmetrically connected patterns (Fig. 5e and Methods 4.4.6). Stimuli in the sensory areas can be aligned with the first pattern in the local and asymmetric-to-asymmetric communication sub-space, and thus referred to as salient, or not aligned, and thus referred to as non-salient (Fig. 5e). We use random patterns for non-salient stimuli that are approximately orthogonal to the communication sub-space in large networks (Methods 4.4.6).

**Figure 5:**
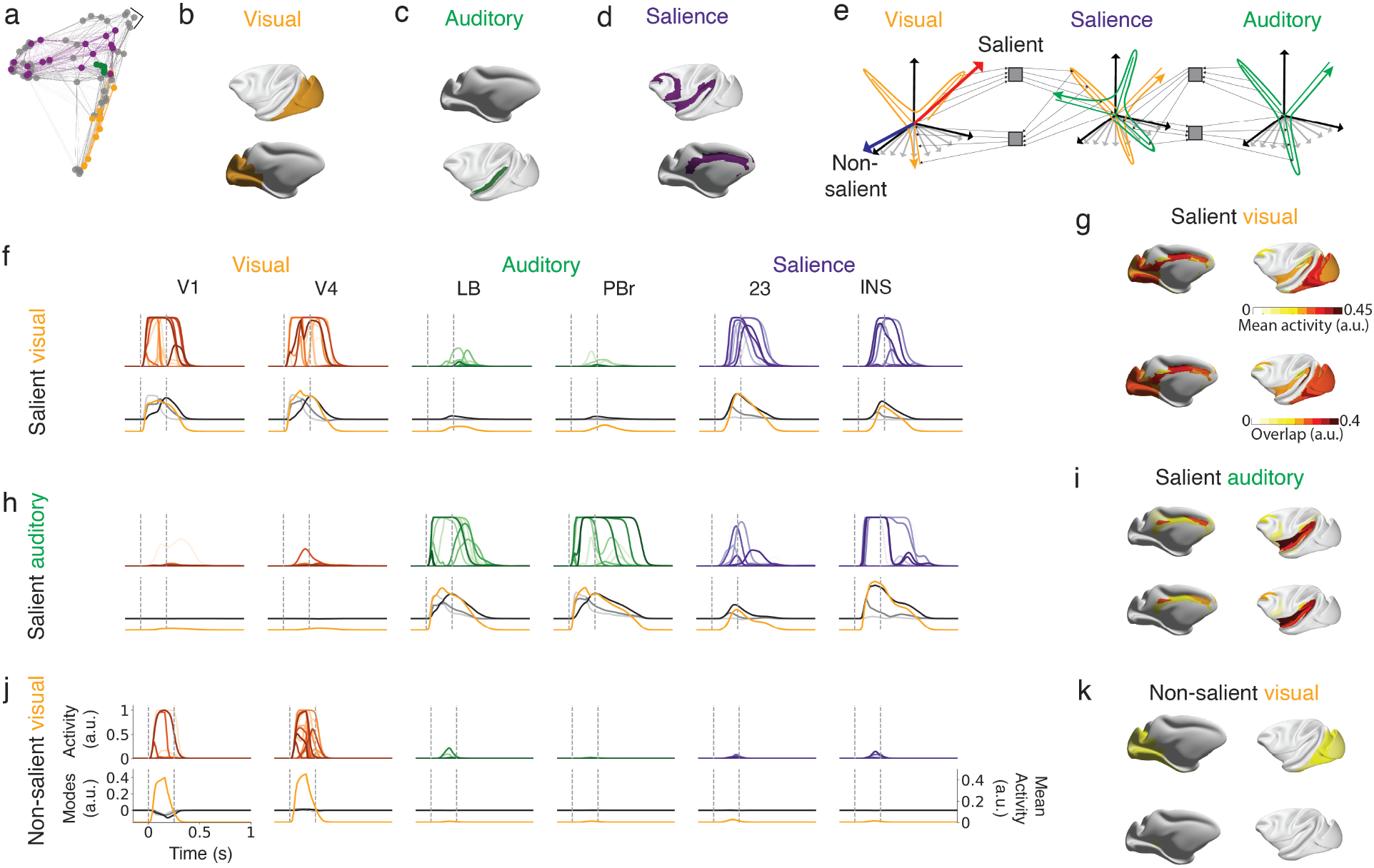
Dynamic routing of sensory stimuli via saliency-dependent communication sub-spaces. (a) dMRI connectivity graph in diffusion map space (Methods 4.2). The edges and nodes that belong to the visual, auditory, and salience networks are colored yellow, green, and purple, respectively. (b) Visual areas. (c) Auditory areas. (d) Salience network areas. (e) Schematics of an asymmetric-to-asymmetric communication subspace that connects visual and auditory networks with the salience network through asymmetric-to-asymmetric communication subspace. Stimuli into the sensory areas (visual or auditory) aligned with the asymmetric-to-asymmetric communication subspace are referred to as salient. An example of a salient visual stimulus is indicated by a red arrow. Stimuli that are not aligned with the communication subspace are referred to as non-salient. An example of a non-salient visual stimulus is indicated by a blue arrow. (f) Network response for a salient visual stimulus. (h) Network response for a salient auditory stimulus. (j) Network response to a nonsalient visual stimulus. (f, h, j) Neural dynamics for 15 representative units (top), overlaps (bottom left axis, grays), and mean activity (bottom right axis, yellow) for six representative areas. The first and second columns correspond to visual areas V1 and V4 (activity traces in red). The third and fourth columns correspond to auditory areas LB and PBr (activity traces in green). The fifth and sixth columns correspond to the salience network areas 23 and INS (activity traces in purple). The leftmost gray dashed line represents the moment when all visual areas (f and j) or auditory areas (h) are transiently stimulated at time 0s for 150 ms. (g, i, k) Mean activity and overlap with the third pattern in the sequence for a salient visual (g), salient auditory (i), and nonsalient visual (k) stimuli at 250ms after the stimulus onset indicated in the rightmost dashed gray line.

When we stimulate the visual network with a salient input aligned with the first pattern of the sequence that comprises the local asymmetric connectivity (Fig.5e), elevated mean activity is widespread across the visual cortex and cortical areas belonging to the salience network (Fig.5f and g; Figs. S10, S11). The overlaps in the visual and salience networks exhibit transient sequential dynamics, with the overlaps being transiently activated in sequential order (Fig.5f; Figs. S10, S11). In the salience network, an increase in the overlap is observed from the first to the third overlap, with the highest value corresponding to the third pattern (Fig.5f; Figs. S10, S11). When the auditory network is stimulated with a salient stimulus, similar network dynamics are observed. In this case, both the auditory and salience networks, but not the visual network, exhibit elevated mean activity and a transient sequential increase in the overlaps (Fig.5h and i; Figs. S12, S13). Lastly, when the visual network is stimulated with non-salient stimuli, the salience network shows very small transient activity (Fig. 5j and k; Figs. S14, S15). A similar effect is observed when the auditory network is stimulated with non-salient stimuli (Figs. S16, S17).

Therefore, in our model, non-salient stimuli in the sensory areas, that are not aligned with the asymmetric-to-asymmetric communication subspace will not elicit transient sequential activity in the salience network areas. Salient stimuli, which are aligned with the communication subspace, lead to transient and widespread elevated activity in the salience network.

### 2.7 Salient-frontoparietal interaction gates DMN-DAN transition

It has been suggested that the salience network, which transiently activates in response to salient stimuli, may mediate the switch between the DMN and DAN [17, 18]. In this process, salient stimuli might gate the transition from internally oriented to externally oriented attention. Here, we propose a mechanism for this salience network dependent gating based on communication subspaces.

To investigate this mechanism, we built macaque and human MRNNs. For the macaque MRNN, as in the previous sections, we aligned the activation maps of the salience, frontoparietal, DMN, and DAN networks in humans to the macaque cortex using cross-species functional alignment (Methods 4.3.2; Table 1). For the human MRNN, we used the canonical networks from [16] (Methods 4.3.1; Table 2).

**Table 2:** Cognitive networks and corresponding cortical areas for the human cortex.

| Cognitive network | Cortical area |
| --- | --- |
| Frontoparietal Network (FPN) | 11l, 23d, 31a, 33pr, 44, 46, 55b, 7Pm, 8Av, 8BM, 8C, 9a, 9-46d, a9-46v, a10p, a32pr, a47r, AV, FJa, FSa, FSp, i6-8, P1, P2, PFm, PHT, POS2, p9-46v, p10p, p47r, SC, s6-8, TE1p |
| Default Mode Network (DMN) | 7m, 8Ad, 8BL, 9a, 9m, 9p, 10d, 10r, 23d, 31a, 31pd, 31pv, 45, 47l, 47m, 47s, a24, a47r, d23ab, d32, p24, p32, PCV, PGi, PGs, POS1, SFL, s32, s6-8, SC, STGa, STSda, STSdp, STSva, STSvp, TE1a, TE1m, TE1p, v23ab |
| Dorsal Attention Network (DAN) | 6a, 6r, 7AL, 7Am, 7PC, 7PL, AIP, FEF, FJp, FST, MST, MIP, PCV, Pd, PE1, PGp, PH, PFt, PIP, POS1, P0, PS1, Pv, TE2p, TPOJ2, TPOJ3, VIP. |
| Salience Network | 23c, 33pr, 43, 44, 45, 55b, 5mv, 6ma, 6r, 9-46d, AAIC, a24pr, FOP1, FOP3, FOP4, FOP5, M, P, PEF, PF, PFcm, PFop, Pol1, Pol2, p24pr, p32pr, PSL, SCEF, SFL, STV, TPOJ1, TPOJ2 |

We begin by considering MRNNs of the macaque and human cortex with connectivity between the frontoparietal network, DMN, and DAN, as shown in Fig.4 (see also Fig.6a and Methods 4.4.7). As previously demonstrated, this connectivity results in competition between the DMN and DAN, with the frontoparietal network exerting control (Fig.4). Inputs to the frontoparietal network govern the transitions between FP-DMN and FP-DAN attractor states (Fig.4). We incorporate the salience network into our MRNNs, connecting regions within it through recurrent asymmetric-to-asymmetric communication subspaces, as the MRNN in Fig.5. To model interactions between the salience and frontoparietal networks, we introduced an asymmetric-to-symmetric communication subspace that projects activity from the first pattern in the asymmetric connectivity motif in the salience network to the frontoparietal network pattern aligned with the FP-DAN attractor (Fig.6a and Methods 4.4.7).

We initialize the MRNNs in a state aligned with the FP-DMN attractor and stimulate salience network areas with a 150ms input aligned to the first pattern of the local asymmetric connectivity motif. Before the stimulation, the population activity overlap with the FP-DMN remained stable and elevated in the frontoparietal and DMN areas, while the FP-DAN overlaps were consistently low across all cortical areas (Figs. 6d, e, S18, S19 for macaque cortex; Figs. 6j, k, S20, S21 for human cortex) similar to Fig. 4. Most frontoparietal and DMN areas showed elevated mean activity (Fig. 6f, S18 for macaque cortex; and Fig. 6i, S21 for human cortex), although gradients of mean activity were observed across the cortex. In response to that input, areas in the salience network exhibit transient sequential dynamics, with the two overlaps being transiently activated in sequential order (Figs. 6d, e, S18, S19 for macaque cortex; Figs. 6j, k, S20, S21 for human cortex). The frontoparietal network shifted from the FP-DMN to the FP-DAN attractor (dashed line in Figs. 6b, S19 for macaque cortex; Figs. 6c, S21 human cortex), the FP-DMN overlap decreased in the frontoparietal and DMN areas, while the FP-DAN overlap increased (Figs. 6d, g vs. e, h for macaque cortex; Figs. 6j, k vs. m, n for human cortex). Consequently, the mean activity decreased in the DMN areas and increased in the DAN areas post-transition (Figs. 6f vs. i for macaque cortex; Figs. 6l vs. o for the human cortex).

**Figure 6:**
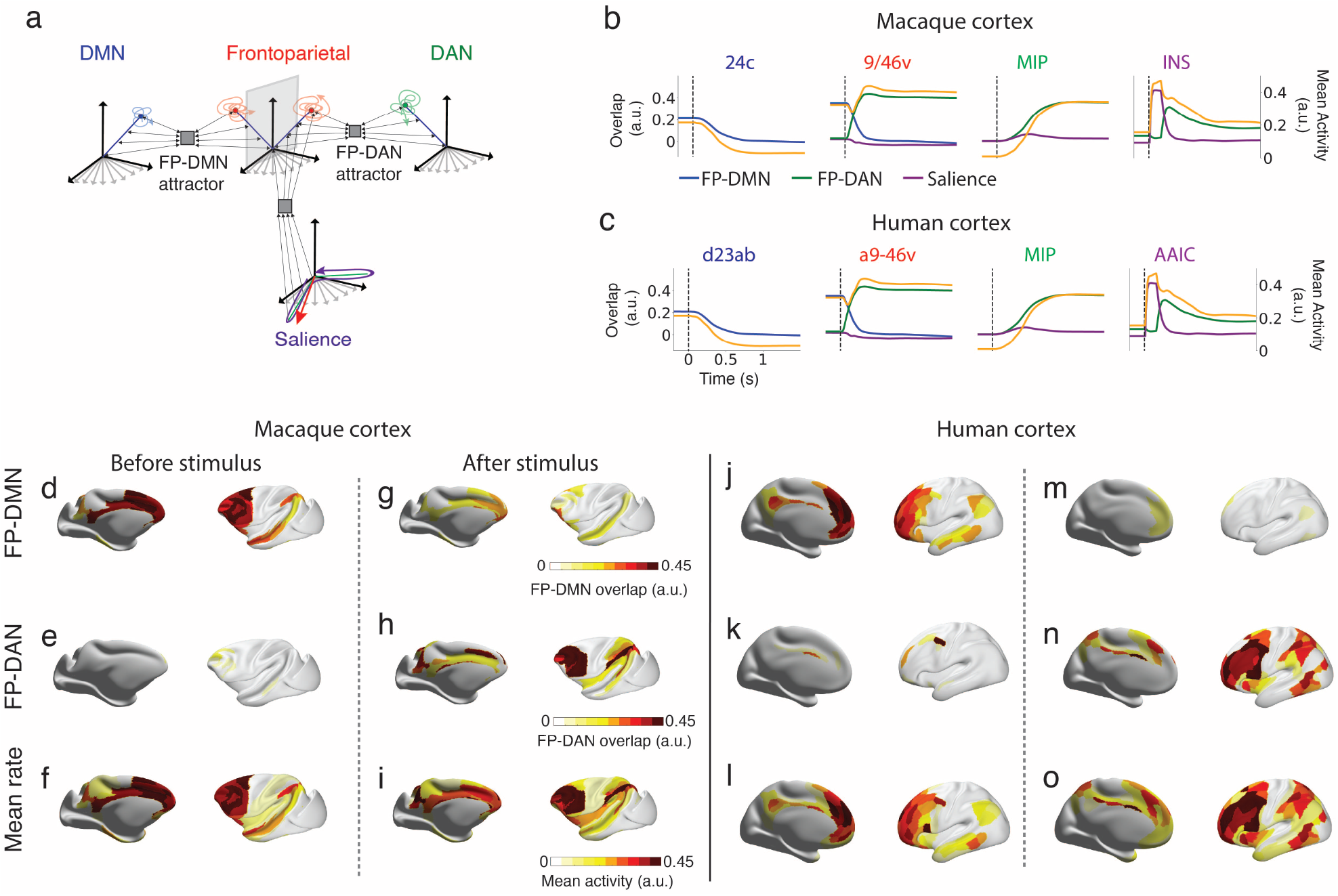
Salient and frontoparietal networks interaction gates the transition between DMN and DAN. (a) Schematic illustrating the interactions between the salience network, frontoparietal network, DMN, and DAN. A transient stimulus aligned with the local asymmetric connectivity pattern in the salience network gates transient activity. This activity aligns with the salience-frontoparietal asymmetric-to-symmetric communication subspace, triggering a transition from the frontoparietal-DMN attractor state to the frontoparietal-DAN attractor state. (b, c) Network response to stimulation to the salient network for the macaque (b) and human (c) cortex. Overlaps and mean activity in four areas: DMN (24c in macaque cortex, d23ab in human cortex), frontoparietal (9/46v in macaque cortex, a9-46v in human cortex), DMN (MIP in both), and salience (INS in macaque cortex, AAIC in human cortex). Blue: overlap FP-DMN attractor; Green: overlap FP-DAN attractor; Purple: overlap with the pattern stimulated in the salience network. The gray dashed line marks the transient stimulation of salience areas at 0s for 150 ms. Overlap for FP-DMN attractor in the macaque (d, g) and human (j, m) cortex, and for FP-DAN attractor in the macaque (e, h) and human (k, n) cortex. Mean activity for the macaque (f, i) and human (l, o) cortex. Panels (d–f, j–l) correspond to 500ms before salience network stimulation, and panels (g–i, m–o) show data 400ms after stimulation for the macaque and human cortex, respectively.

Thus, our model introduces a mechanism based on communication subspaces for saliency-dependent gating between the DMN and DAN, mediated by the frontoparietal network.

## 3 Discussion

In this work, we develop a class of connectivity-constrained whole-cortex models for macaque and human brains to understand how neocortex-wide activity can dynamically organize into distinct large-scale cognitive networks through communication subspaces. Our modeling work explains key aspects of cognitive network dynamics and interactions observed experimentally. First, it provides a network mechanism for the anti-correlation [85, 9] between the default mode network (DMN) and the dorsal attention network (DAN). Second, it presents a mechanism based on communication subspaces for the observed frontoparietal coupling with the DMN and DAN [22]. Third, it proposes a bottom-up routing mechanism for the observed salience-dependent DMN to DAN switching [17, 18]. Fourth, it demonstrates that the frontoparietal network can display a coexistence of stable and dynamic coding with strong temporal variability and substantial heterogeneity, qualitatively matching the stable [49, 50, 51, 52, 53] and dynamic [54, 51, 52, 53, 55, 57, 58, 59] coding observed in electrophysiological recordings within the frontoparietal network.

We model within-area connectivity as a mixture of symmetric [60, 61, 62], asymmetric [63, 64, 65], and random [66] motifs, producing either stable (attractor) or transient (sequential) heterogeneous dynamics, integrating classic local circuit models into multi-regional models for cortical dynamics. Unlike previous connectivity-constrained multi-regional models [33, 34, 35, 36, 86, 41, 42, 43, 44, 45, 46, 47], we use connectivity data as a proxy for inter-area connection sparsity rather than total synaptic strength. This allows us to model long-range projections as sparse low-rank communication subspaces. Inspired by recent electrophysiological recordings that investigate two-area interactions [28], our model routes local dynamics through these low-rank communication subspaces, depending on the partial alignment of sparse long-range projections with local connectivity.

Overall, our work provides a theoretical framework that bridges neuroanatomy, electrophysiology, and neuromiaging for understanding the dynamic routing of large-scale cognitive networks during cognition.

### 3.1 Frontoparietal control of the DMN-DAN transition

Neuroimaging experiments suggest that the frontoparietal network may flexibly control transitions between the DMN and DAN during behavior by coupling with both cognitive networks [22]. These networks exhibit significant anti-correlation across a range of states [85, 9] and are thought to realise the competition between externally- and internally-directed attention. However, despite the central importance of this phenomenon to human cognitive neuroscience, the mechanism for how this competition may play out is not known, and computational modeling proposals for the underlying mechanism are rare [87, 88]. Here, we propose a mechanism based on dynamic control of two distributed attractors mediated by communication subspaces. One attractor involves the frontoparietal network and the DMN (FP-DMN), while the other involves the frontoparietal network and the DAN (FP-DAN). This result is consistent with multiple recent reports of two subnetworks within the frontoparietal network [24, 25, 26, 89], and demonstrates an attractor mechanism by which competition between these subnetworks can drive the transition between the DMN and DAN. Our proposed mechanism - communication subspaces connecting the frontoparietal network with distinct higher cognitive networks - is also a mechanism by which the frontoparietal network can be recruited during challenging tasks engaging various other cognitive networks, and therefore helps explain how this network can function as a “multiple demand network” [19].

It has been suggested that the salience network, which transiently activates in response to salient stimuli, may mediate the switch between the DMN and DAN [17, 18]. When incorporating the salience network and modeling interactions between the salience and frontoparietal networks, we introduced an asymmetric-to-symmetric communication subspace that reads out activity from one dimension comprising the local asymmetric connectivity in the salience network and projects to a direction in the frontoparietal network activity space aligned with the FP-DAN attractor. We show that transient inputs to the salience network, which gate sequential population activity within it, can control the DMN-DAN switch through these asymmetric-to-symmetric communication subspaces, aligning transient salience network activity with FP-DAN attractor dynamics. This introduces a mechanism for saliency-dependent gating between the DMN and DAN, mediated by the frontoparietal network via communication subspaces.

### 3.2 Distributed stable and dynamic coding

Working memory and decision-making involve a distributed network of areas in the frontal and parietal regions [19, 90, 91, 57, 92]. Recent models of have accounted for stable persistent activity [42, 41, 43, 47] and activity-silent [42] maintenance of short-term/working memories throughout the frontoparietal network. The distributed frontoparietal network in our model departs from previous models by displaying both stable [49, 50, 51, 52, 53] and dynamic [54, 51, 52, 53, 55] population coding. The dynamic coding is generated through sequential population activity, driven by asymmetric local connectivity [64, 63, 71, 74, 73] and distributed asymmetric-to-asymmetric communication subspaces within the frontoparietal network. This dynamic coding coexists with stable encoding, which occurs via attractor dynamics [61, 62, 80], driven by distributed ensembles of units connected through symmetric-to-symmetric communication subspaces. Our model shows that these two types of coding can co-exist in the frontoparietal large-scale network, and provides a mechanistic explanation for recent observations of dynamic coding in the parietal cortex during decision-making tasks [57], the motor cortex during action planning [58], and the prefrontal cortex during navigation working memory tasks [59], while also accounting for the classic observations of stable delay activity during working memory tasks [93, 94, 2, 50].

In the frontoparietal network, during delay periods and stable coding, the encoding of memoranda remains stable at the population level, while the activity of individual neurons can exhibit significant heterogeneity [49, 95, 50]. The mechanism underlying this combination of distributed stable encoding and heterogeneous dynamics in our network is the distributed chaotic dynamics and stable encoding previously observed in local circuit models [81, 83, 82]. We derived a mean field theory that successfully describes the overlaps for each area, similar to the approach described in [96]. Our theory is designed to describe the dynamics when the network reaches stable fixed points, approximately capturing the overlap for the chaotic phase. However, our theory cannot predict the transition to chaos [66, 81, 83, 82].

### 3.3 Dynamic routing of salient sensory stimuli via communication subspaces

In our model, salient sensory stimuli are defined as those that align with the asymmetric-to-asymmetric communication subspaces connecting the sensory cortices to the salience network. Salient stimuli are routed from the sensory cortices to the salience network, eliciting sequential population activity. Non-salient sensory stimuli, defined as those not aligned with these communication subspaces, do not elicit activity in the salience network. We model the representations of auditory and visual stimuli in the salience network as two distinct population sequences, preserving the identity of the stimuli (e.g., auditory vs. visual) within the network.

The mechanism we propose for salience-dependent gating is bottom-up, relying on the alignment of sensory signals in the sensory cortices (bottom) with predefined communication subspaces in order to be routed to higher-order cortical regions in the salience network (up). In our model, these communication subspaces are *hard-coded*, meaning they do not change over time. This is consistent with the concept of saliency maps [97] in vision science.

In a two-area network model for context-dependent decision making [98], communication between auditory cortex and the prefrontal cortex can be selectively routed in a context-dependent fashion through communication subspaces. Such context-dependent mechanisms may also be implemented in our network to selectively route auditory or visual information depending on the context controlled by the frontoparietal network.

Disinhibitory connectivity motifs involving different cell types and cell-type-specific local connections have been proposed as a mechanism for routing sensory information [99]. Our model does not include cell types, and exploring the implementation of communication subspaces through cell-type-specific connections is a potential direction for future research.

### 3.4 Communication subspaces as routing mechanism of large-scale cognitive networks

In our MRNNs, distributed attractor dynamics and sequential population activity shape the temporal evolution of the cognitive networks during task performance. Coordinating the activity of these networks is governed by the selective routing of neural activity through communication subspaces. These subspaces act as dimensionality bottlenecks [28, 29, 31, 30], restricting the maximum dimension of routed signals and allowing for the dynamic reconfiguration of cognitive networks according to task demands depending on the alignment of the local dynamics with the communication subspaces.

Multiple different mechanisms for large-scale brain communication have been proposed, see, for example, the Perspective in [100]. Multiple groups argue that multi-regional communication is achieved through the synchronization of oscillatory activity [101, 102, 103, 104, 105, 106, 107, 108, 109], which serves as the mechanism enabling coordination across brain regions during cognition. Although this remains a topic of scientific debate [110]. The mechanism proposed here for routing large-scale cognitive networks is not mutually exclusive with the synchronization hypothesis; both mechanisms may contribute to the coordination and routing of large-scale cognitive networks.

In our current model, the local dynamics and the communication subspaces are low-dimensional. This is due to the fact that the local connectivity is low-rank [83], and the dimensionality of the mean activity is at most equal to the total number of patterns defined by the symmetric and asymmetric connectivity motifs. Communication subspaces act as bottlenecks for routing local dynamics, as the number of dimensions available for inter-area communication is smaller than the dimensionality of local dynamics and significantly smaller than the number of neurons, i.e., Rank(𝒞) < *p*_A_ + *p*_S_ ≪*N*. While neural activity observed during neuroscience tasks is generally low-dimensional, much smaller than the number of recorded neurons, recent large-scale neural recordings show that neural activity in the brain can be high-dimensional [111, 112], with dimensionality scaling linearly with the number of neurons. We expect our theory to remain applicable in this high-dimensional regime, one can envision an MRNN model with high-dimensional local dynamics that scale linearly with network size, alongside communication subspaces whose dimensionality also scales linearly with network size but remains smaller than the dimensionality of the areas they connect.

### 3.5 Multi-regional RNN models

Computational models of multi-regional dynamics that incorporate anatomical constraints have typically been constructed using highly simplified mean field models, which represent the overall activity of each area with only a few dynamic variables [33, 34, 35, 36, 86, 41, 42, 43, 44, 45, 46, 47]. These models have provided valuable insights into the network mechanisms driving multiregional dynamics during the resting state [33, 34, 35, 36], working memory [41, 42, 43, 44], decision-making [45, 46] and conscious access [47]. However, by modeling local circuit dynamics with only a few variables, these models fail to capture the high-dimensional activity of large neural populations, which is likely crucial for understanding how information is represented and dynamically routed in multi-regional brain systems.

Theoretical studies have explored the dynamics of modular networks composed of multiple units within each module [113, 96, 114, 115, 116]. Notably, recent work has studied the dynamics of large networks with a rank-one structure within and across modules [114]. However, these models primarily focus on describing transitions to chaotic dynamics [113, 96, 114, 115] or the storage capacity of attractor states [116] and do not model specific experimental data or incorporate biological constraints.

Recently, MRNNs with multiple units in each area have been employed to fit multi-regional recordings using machine learning methods [117]. While these models have multiple units in each brain region, the observed dynamics can be challenging to interpret [48]. Moreover, these models are solely constrained by neural activity, which, due to the non-convex nature of the training process [118], may lead to the discovery of multi-regional interactions during the fitting procedure that are inconsistent with anatomical data. Here, we build and analyze a class of connectivity-constrained MRNN models to dissect how cognitive network dynamics are shaped by the interplay between attractor dynamics and sequential population activity, coordinated by communication subspaces. We constrain our model using anatomical and neuroimaging data. Unlike previous large-scale models, anatomical data were used to constrain the sparsity of inter-area connections but not the synaptic weights between units. Thus, anatomy serves as a sparsity scaffold for synaptic weights, providing a soft constraint by penalizing weaker anatomical connections with sparser projections.

Due to the mixture of symmetric, asymmetric, and random connectivity, the network dynamics can be interpreted as the overlap of population activity in each area with ensembles of units comprising the symmetric and asymmetric connectivity motifs, as demonstrated recently for local circuit models [65, 75]. This mixture of connectivity within the local circuits of each area leads to a straightforward interpretation of the communication subspaces as interactions between symmetric and asymmetric connected ensembles. We show that the spatial positioning of symmetric-to-symmetric, asymmetric-to-asymmetric, and asymmetric-to-symmetric communication subspaces allows different types of distributed attractors or transient sequential activity to be routed within the network. We therefore provide a model of the representation and dynamic routing of information within and across cognitive networks via local connectivity-aligned communication subspaces.

## 4 Methods

### 4.1 Anatomical datasets

#### 4.1.1 Macaque cortex diffusion MRI tractography

For macaque structural connectivity data, we used a fully reconstructed 91×91 area connectivity matrix from [69], via the BALSA neuroimaging data sharing site https://balsa.wustl.edu/study/W336 [119]. Briefly, those authors acquired the data from postmortem diffusion MRI scans of an immersion-fixed brain from an adult male macaque monkey *(M. fascicularis)*, infused with gadolinium contrast. The acquisition was on a 4.7T Bruker machine, 120 diffusion-weighting directions b = 8000 s/mm^2^. Probabilistic tractography was performed using FSL’s probtrackx [120], seeded from the cortical grey/white matter boundary. The connection weights were normalised such that the normalised weight of a pathway originating in some area A and terminating in an area B is defined as the ratio of the number of streamlines originating at A and terminating at B to the total number of streamlines that either originate at A or terminate at B (excluding within-area connections). The connectivity matrix was then symmetrised, by averaging the connectivity weights for each pathway across the two directions. For further details, see [69]. In this work, we exclude the Subiculum and the Piriform cortex from this dataset.

#### 4.1.2 Macaque cortex T1w/T2w

We obtained the macaque cortex T1w/T2w data from the Yerkes19 group average dataset (https://balsa.wustl.edu/reference/976nz). Briefly, this was based on structural T1-weighted and T2-weighted (0.5 mm isotropic resolution) imaging of 19 adult macaques scanned at the Yerkes National Primate Center, Emory, USA. The scans were processed through a macaque variant of the Human Connectome Project pipeline [121] to extract surfaces and align the scans to a template. We then averaged the T1w/T2w values within each of the 91 areas in the Lyon (M132) cortical parcellation [122].

#### 4.1.3 Human cortex dMRI tractography

For human structural connectivity data, we used a fully reconstructed 180×180 area connectivity matrix of the left cortical hemisphere produced in [36]. The data in that study was based on 334 subjects from the Human Connectome Project [121]. The 180 areas represent the areas in the cortical parcellation in [121]. Similarly to the macaque data, tractography was performed using FSL’s probtrackx [120].

#### 4.1.4 Human cortex T1w/T2w

The human T1w/T2w data was based on the S1200 subject release of the Human Connectome Project, and downloaded from BALSA (https://balsa.wustl.edu/reference/pkXDZ).

We then averaged the T1w/T2w values within each of the 180 areas of the Glasser parcellation in the left cortical hemisphere [123].

### 4.2 Diffusion map embedding on dMRI connectivity

We use the diffusion map embedding method [124] for analyzing the dMRI connectivity data. This method has been recently successfully applied to human and macaque connectomes for uncovering the organization of the transmodal default-mode network [125]. The method is based on an hypothetical *diffusion processes* that diffuses from the nodes along the edges of the dMRI connectome. This diffusion process give rise a to a *diffusion space* in which a distance between cortical areas is defined. Areas that are closer in diffusion space have stronger connections and share a larger number of paths connecting them, while areas that are farther apart they have weaker connections and share fewer paths. Using this method, the dMRI connectivity is embedded in a low number of ‘principal gradients’, which are the principal component of the normalized graph Laplacian of the diffusion process, leading to a low dimensional embedding of the dMRI connectivity. Since dMRI connectivity matrix is symmetric, a requirement for applying the method, we applied directly the diffusion map method on the dMRI connectivity matrix *W*.

### 4.3 Cognitive networks

#### 4.3.1 Human cognitive networks

The human cognitive networks are the canonical networks described by Yeo, Krienen and colleagues [16], aligned to the Human Connectome Project FS-LR 32k space and downloaded from BALSA (https://balsa.wustl.edu/reference/6V6gD).

#### 4.3.2 Definition of cognitive networks in the macaque cortex

A set of canonical functional networks that co-activate during rest and tasks have been widely replicated in the human and monkey brain. The location and extent of each network has been better delineated in the human brain. Therefore, we mapped the human networks to the macaque cortex using cross-species functional alignment, and then assigned each cortical area to the overlapping networks, in proportion with the degree of overlap. Cross-species functional alignment relies on constructing a cross-species joint-similarity matrix (with cross-species similarity of each pair of vertices based on functional connectivity with 27 homologous landmarks), calculating cross-species gradients (by applying the diffusion map method to the joint-similarity matrix), and creating a vertex-to-vertex mapping based on multimodal surface mapping [126] using these gradients. For details please see [77].

### 4.4 Multi-regional recurrent neural network

#### 4.4.1 Dynamics and connectivity structure

We model the neocortex as a large-scale recurrent neural network composed of *n* cortical areas. In the case of the macaque cortex *n* = 89 and in the human cortex *n* = 180. We assume that each cortical area contains an equal number of units *N*. The value is *N* = 400, unless specified otherwise. Each unit represents a neural pool (or assembly) that is selective for specific features rather than an individual neuron. The dynamics are given by the standard rate equations [76, 61] written in the current formulation

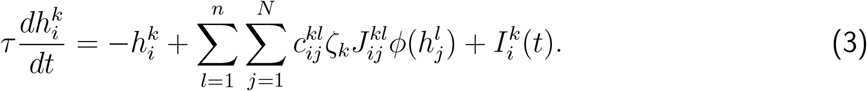

The variable 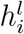 is the current to unit *i* in area *l*, while 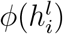 corresponds to the activity. The intrinsic time-scale of the units was taken to be *τ* = 60 ms, modeling the excitatory dynamics dominated by NMDA receptor dynamics. The function *ϕ* is the input-output transfer function of the model that, in our case, is sigmoidal: 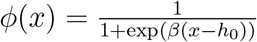, with *β* = 2.5 and *h*_0_ = 1.13.

The function 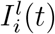 corresponds to the external input current to neuron *i* in area *l*.

The variables 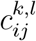 model the existence or absence of a synaptic interaction between pre-synaptic unit *j* in area *l* and post-synaptic neuron *i* in area *k*. Therefore, the variables 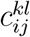 can take values 0 or 1. In our model, the long-range connectivity between areas is sparse and random. We model it as independent and identically distributed (i.i.d.) Bernoulli random variables with projection-dependent probabilities given by connectivity data

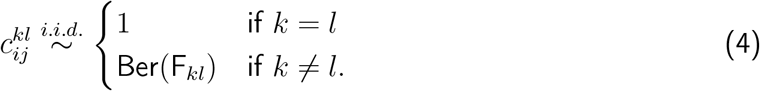

The inter-area connectivity matrix F_*kl*_ corresponds to the normalized dMRI tractography (more in Methods 4.1) for macaque monkey (Fig. 1h) and human (Fig. S1). Therefore, although the inter-area connectivity is random, the sparsity of the connections is constrained by dMRI tractography data (see Fig. 1h and i).

The parameter *ζ*_*k*_ represents an area-dependent cortical gradient that affects the strength of local and long-range incoming connections to a local RNN *k*. Our model assumes linear variation in the strength of these connections across cortical areas. This assumption is based on the strong correlation observed between spine count per neuron, which serves as a proxy for the total number of excitatory synaptic inputs, and the hierarchical position of cortical areas [127, 67]. We approximate the hierarchical position of each cortical area using MRI-derived T1w/T2w neuroimaging values (Figs. 1j and S1b). T1w/T2w values are inversely correlated with spine count [128]. Thus, the cortical gradient is defined by

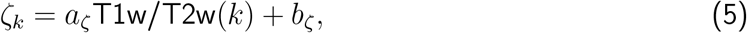

where *k* is the local RNN index. The parameter values are *a*_ζ_ = 0.6, *b*_ζ_ = 0.4 for the macaque MRNN, and *a*_ζ_ = 1, *b*_ζ_ = 0.1 for the human MRNN.

Their local synaptic weight 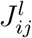 between a pre-synaptic unit *j* and a post-synaptic unit *i* in region *l* consists of a combination of symmetric (Fig. 1a), asymmetric (Fig. 1b), and random (Fig. 1c) connectivity motifs

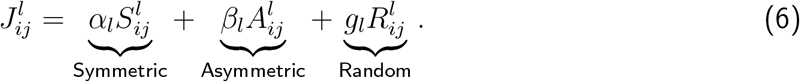

The symmetric connectivity is such that feedback and feedforward synaptic weights have the same strength, i.e., 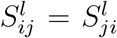 (Fig. 1a). As in network models for attractor dynamics [61, 62] the symmetric connectivity motif in our model is built as the sum of the outer product of *N*-dimensional patterns

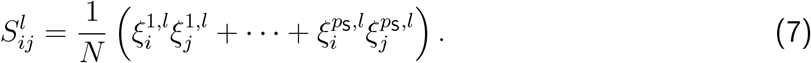

The entries of these patterns are independently and identically distributed (i.i.d.), taking values of −1 and 1 with a probability of 0.5. The number of symmetric connectivity motifs per area is *p*_*S*_. This type of symmetric connectivity in Eq. (7) generates strong recurrent loops (Fig. 1a), producing multi-stable attractor dynamics and stable persistent activity in recurrent networks [60, 61, 62, 70].

The asymmetric connectivity motif is such that 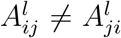 and has an effective feed-forward structure [63, 64, 71, 72, 73, 74, 65] given by the sequential outer product of *p*_A_ patterns:

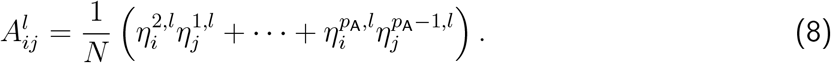

These patterns have the same statistics as the patterns in the symmetric connectivity motif. The asymmetric connectivity motif in Eq. (8) produces transient sequential population activity in recurrent networks [64, 63, 71, 65, 74]. Throughout the paper, these patterns can be the same 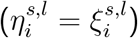 [75, 129] or different from the symmetric patterns. Additionally, multiple sequences can be embedded in each local RNN’s connectivity by adding multiple independent sequences of patterns [65].

The random connectivity is given by

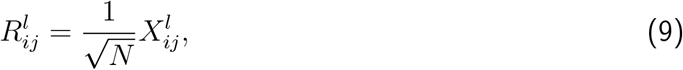

where each entry of the connectivity is independent and identically distributed as a Gaussian variable, 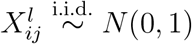 (Fig. 1c). This connectivity motif is known to produce high-dimensional chaos in RNNs characterized by strongly temporally heterogeneous dynamics [66].

#### 4.4.2 Arrangements of local connectivity motifs and communication subspaces

The long-range connectivity of our MRNNs (Eq. (2)) exhibits significant flexibility in selecting communication subspaces, allowing for multiple possible configurations within a single model. For instance, a MRNN with the same local RNNs can select various types of communication subspaces, such as symmetric-to-symmetric (Fig. 1e), asymmetric-to-asymmetric (Fig. 1f), asymmetric-to-symmetric (Fig. 1g), or a mixture of these types. Importantly, the spatial arrangement of these communication subspaces across the cortex can also vary widely. This selection qualitatively alters the multiregional network dynamics and computations, as demonstrated in the main text. Depending on the selected communication subspace and its spatial arrangement, different types and spatially distributed patterns of multiregional dynamics can be routed. We used the cognitive networks defined using the MRI activation maps (Methods 4.3, Tables 1 and 2) to constrain the spatial arrangement of the connectivity parameters for both the local RNN symmetric and asymmetric connectivity (Eqs. (6–8)) as well as the communication subspaces (Eq. (2)). As demonstrated in the main text, these spatial arrangements restrict different and co-existing attractor and sequential population dynamics to specific subsets of areas that overlap with the cognitive networks’ spatial patterns of activation.

In the next subsection, we will detail the procedure used to constrain the MRNNs’ connectivity and the spatial arrangement of communication subspaces across the cortex for Figs. 2-6. For all our simulations, all local RNNs have the same random connectivity strength, denoted by *g*_*l*_ = *g*_Loc_ = 2. Similarly, the long-range random connectivity strength will be the same for all projections and is denoted by *g*_*kl*_ = *g*_LR_. Its value ranges from 2 to 3 (i.e., *g*_LR_ ∈ [2, 3]) and will be specified accordingly.

#### 4.4.3 Stable population coding and heterogeneous neural dynamics in the frontoparietal network

In Fig. 2, only the local RNNs within the frontoparietal network of the macaque cortex (Table 1) have symmetric local connectivity (see Eq. (6)). The number of patterns generating the symmetric connectivity is *p*_*S*_ = 1 (see Eq. (7)). Consequently, the local RNNs’ connectivity is given by

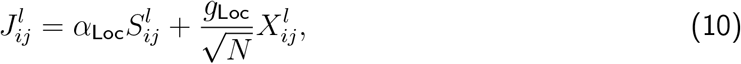

with

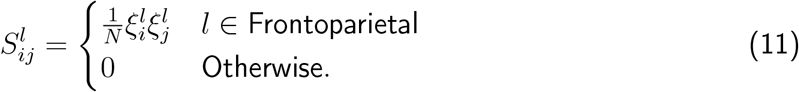

As described in the main text, the entries of the patterns 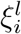 are independently and identically distributed (i.i.d.), taking values of −1 and 1 with a probability of 0.5.

The entries of the random connectivity motif are i.i.d. Gaussian variables, 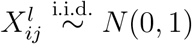.

The relative strength of the local symmetric connectivity is the same across all areas within the frontoparietal network, with *α*_Loc_ = 4.51.

For the long-range projections, the symmetric-to-symmetric communication subspace (Eq. (2)) is given by

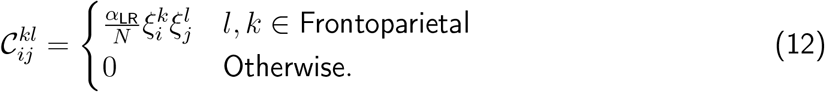

Only the projections connecting areas within the frontoparietal network have a non-zero symmetric-to-symmetric communication subspace. Therefore, the long-range projections are given by

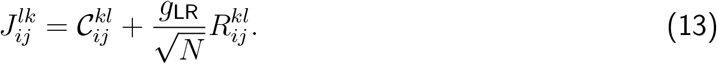

The strength of the symmetric-to-symmetric communication subspace is the same across all frontoparietal projections and is given by *α*_LR_ = 7.48.

Similar to the random connectivity for the local connectivity, the entries of the long-range random connectivity are also i.i.d. Gaussian variables, 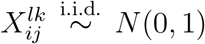. The strength of the long-range random connectivity is *g*_LR_ = 2.75.

The external input current is equal to zero 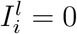.

#### 4.4.4 Dynamic and stable population coding coexist in the frontoparietal network

In Fig. 3, similar to the connectivity for Fig. 2, only the local RNNs within the frontoparietal network of the macaque cortex (Table 1) exhibit both symmetric and asymmetric local connectivity (see Eq. (6)). The number of patterns generating symmetric connectivity is *p*_*S*_ = 1 (see Eq. (7)), while for asymmetric connectivity, it is *p*_*A*_ = 8 (see Eq. (8)). Consequently, the local RNNs’ connectivity is given by

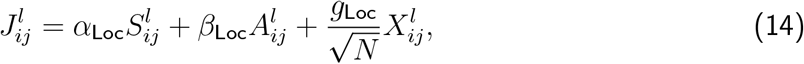

with

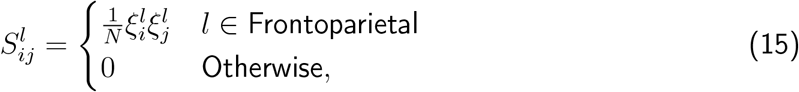

and

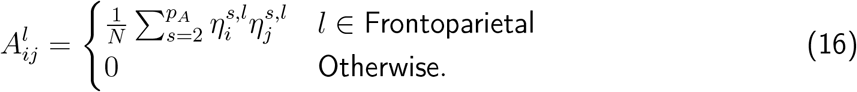

The entries of the patterns 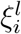 and 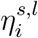 are independently and identically distributed (i.i.d.), taking values of −1 and 1 with a probability of 0.5. Therefore, the patterns comprising the symmetric, 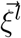, and the asymmetric connectivity motifs, 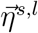, are uncorrelated. The strength of the local symmetric and asymmetric connectivity is the same across all areas within the frontoparietal network, with *α*_Loc_ = 4.41 and *β*_Loc_ = 4.41.

For the long-range projections, the communication subspace (Eq. (2)) is given by

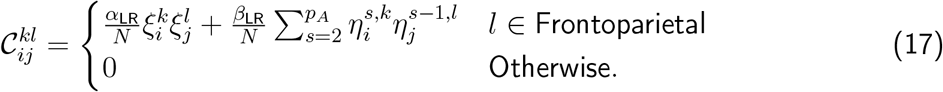

Only the projections connecting areas within the frontoparietal network have a non-zero communication subspace, otherwise 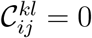. Therefore, the long-range projections are given by

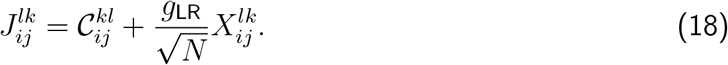

The strength of the symmetric-to-symmetric and asymmetric-to-asymmetric communication subspaces are the same across all frontoparietal projections and are given by *α*_LR_ = 7.14 and *β*_LR_ = 7.14. The strength of the long-range random connectivity motifs is *g*_LR_ = 2.75. The number of units per area is *N* = 500.

For Fig. 3, the external input current is aligned with the first pattern in the sequence and is given by

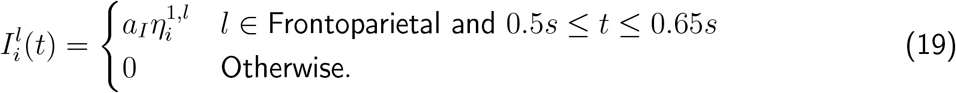

For Figs. S5 and S5, the external input current is aligned with the symmetric-to-symmetric communication subspace and is given b

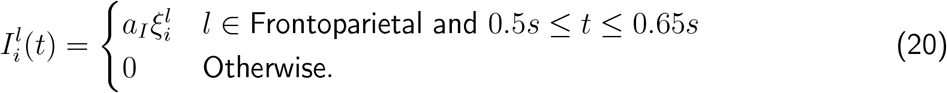

The magnitude of the external input current is *a*_*I*_ = 2 for Figs. 3, S5 and S5.

##### 4.4.5 Frontoparietal attractor dynamics coordinates DMN-DAN interplay

In Fig. 4, only the local RNNs within the frontoparietal, DMN, and DAN networks of the macaque cortex (Table 1) have symmetric local connectivity (see Eq. (6)). The local RNNs’ connectivity is given by

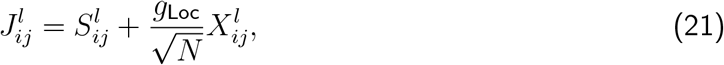

with

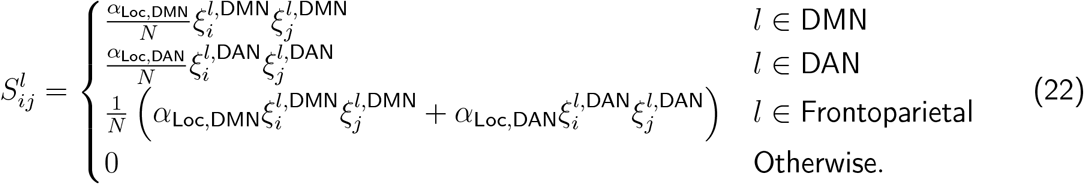

The entries of the patterns 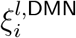 and 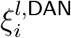 are independently and identically distributed (i.i.d.), taking values of −1 and 1 with a probability of 0.5. Therefore, the patterns 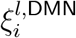 and 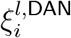, which describe the local connectivity in areas within the union of the DMN and frontoparietal networks, and the union of the DAN and frontoparietal networks, respectively, are uncorrelated. The strengths of the local symmetric connectivity are *α*_Loc,DMN_ = 2 and *α*_Loc,DAN_ = 2.3.

For the long-range projections, the communication subspace (Eq. (2)) is given by

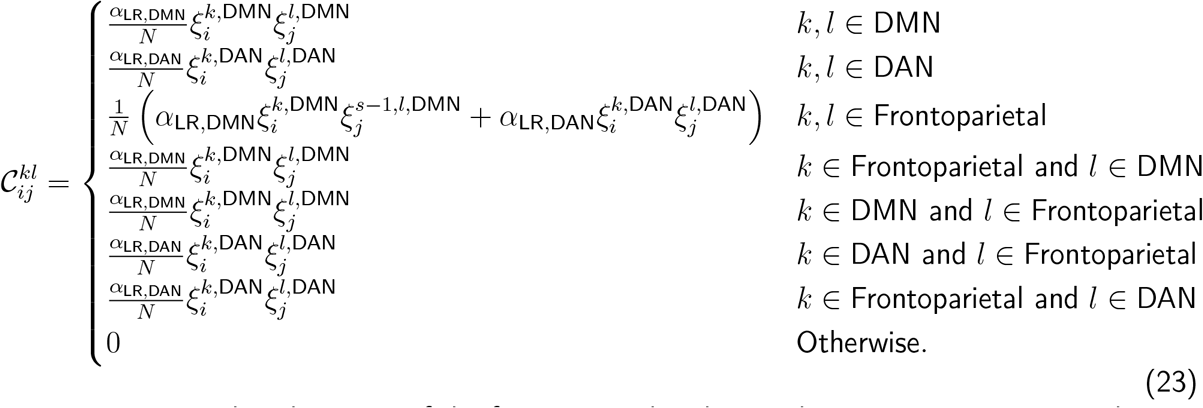

Projections within the union of the frontoparietal and DMN have a communication subspace defined by the vectors 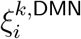, while projections within the union of the frontoparietal and DAN have a communication subspace defined by the vectors 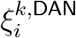. The recurrent connections within the frontoparietal network present both communication subspaces defined by 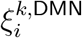 and 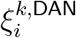. The long-range projections are given by

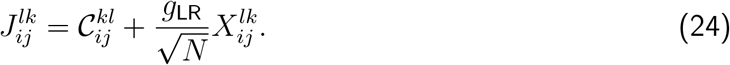

The strength of the symmetric-to-symmetric communication subspaces are given by *α*_LR,DMN_ = 7 and *α*_LR,DAN_ = 8.5. The strength of the long-range random connectivity is *g*_LR_ = 3.

To investigate the transition between the FP-DMN and FP-DAN attractor states, we first introduced a constant input to the frontoparietal network areas. Initially, the input was aligned with the FP-DMN ensemble in the frontoparietal network. At time zero input to the frontoparietal network shifted its alignment from the FP-DMN to the FP-DAN ensemble. The external input is given by

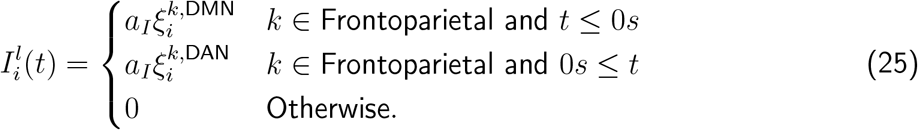

The magnitude of the external input current is *a*_*I*_ = 1.15 for Fig. 4 and the corresponding supplementary figures (Figs. S8 and S9).

#### 4.4.6 Dynamic routing of sensory stimuli via saliency-dependent communication subspaces

In Fig. 5, only the local RNNs within the Auditory, Visual, and Salience networks of the macaque cortex (Table 1) have asymmetric local connectivity (see Eq. (6)). The number of patterns generating asymmetric connectivity is *p*_*A*_ = 3 (see Eq. (8)). The local RNNs’ connectivity is given by

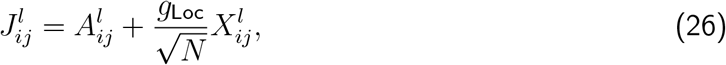

with

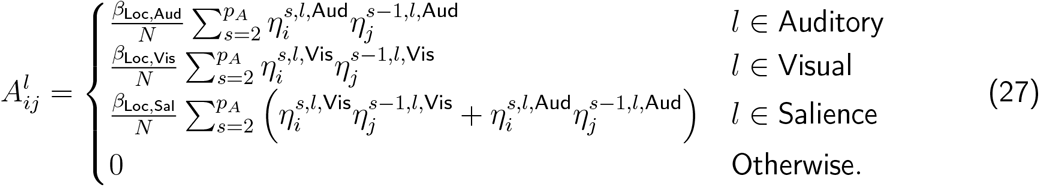

The entries of the patterns 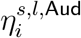 and 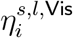 are independently and identically distributed (i.i.d.), taking values of −1 and 1 with a probability of 0.5. Therefore, the patterns 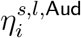 and 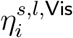,present in areas in the union of the Auditory and Salience cognitive networks and the union of the Visual and Salience cognitive networks, respectively, are uncorrelated. The strengths of the local asymmetric connectivity are *β*_Loc,Aud_ = 6.3, *β*_Loc,Vis_ = 7.35, and *β*_Loc,Sal_ = 4.73.

For the long-range projections, the communication subspace (Eq. (2)) is given by

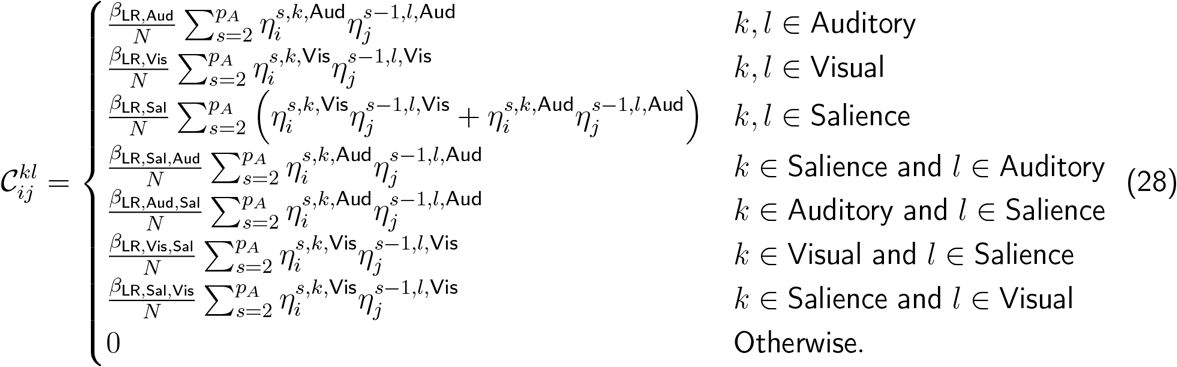

Projections within the union of the salience and visual cognitive networks have a communication subspace defined by the vectors 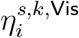, while projections within the union of the salience and auditory cognitive networks have a communication subspace defined by the vectors 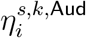. The recurrent connections within the salience network present both communication subspaces defined by 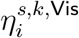 and 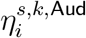. The long-range projections are given by

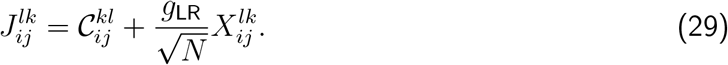

The strength of the asymmetric-to-asymmetric communication subspaces are given by *β*_LR,Vis_ = 10.2, *β*_LR,Aud_ = 11.9, *β*_LR,Sal_ = 7.65, *β*_LR,Sal,Aud_ = *β*_LR,Aud,Sal_ = 34, *β*_LR,Sal,Aud_ = *β*_LR,Aud,Sal_ = 34 and *β*_LR,Sal,Vis_ = *β*_LR,Vis,Sal_ = 27.2. The strength of the long-range random connectivity motif is *g*_LR_ = 2.75.

For the salient visual stimulus in Figs. 5f, 5g, S10, and S11, the external input current is aligned with the first pattern corresponding to the visual asymmetric-to-asymmetric communication subspace and is given by

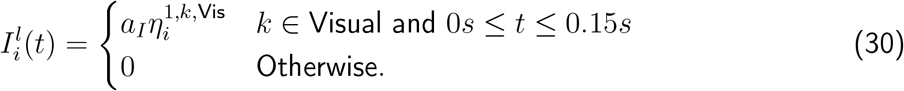

Similarly, for the salient auditory stimuli in Figs. 5h, 5i, S14, and S15, the external input current is aligned with the first pattern corresponding to the auditory asymmetric-to-asymmetric communication subspace and is given by

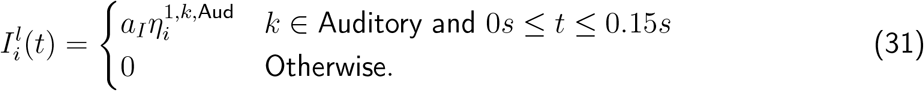

Non-salient visual (Figs. 5j, 5k, S14, S15) and auditory (Figs. S16, S17) stimuli are defined by the functions in Eqs. (30, 31), respectively, replacing 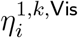 and 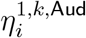 with random, uncorrelated patterns that take values of −1 or 1 with equal probability.

The magnitude of the external input current is *a*_*I*_ = 2 for Fig. 5 and all the corresponding supplementary figures (Figs. S10-S17).

#### 4.4.7 Salience Network-Frontoparietal interaction gates DMN-DAN transition

In Fig. 6, the local RNNs within the frontoparietal network, DMN, and DAN of the macaque and human cortex (Table 1 and Table 2, respectively) are comprised of symmetric local connectivity motifs (see Eq. (6)), while the salience network is generated by asymmetric local connectivity motifs. The local RNNs’ connectivity is given by

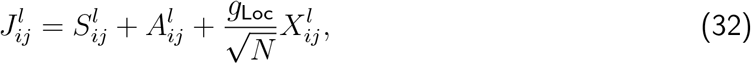

with

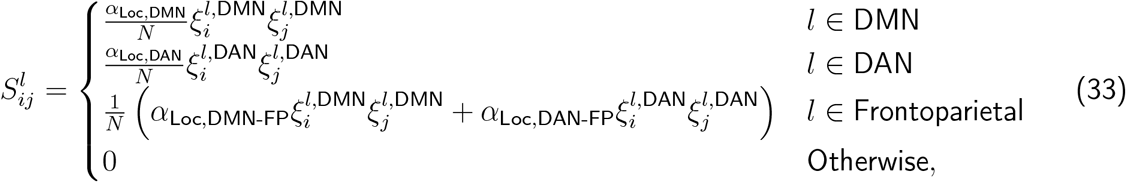

and

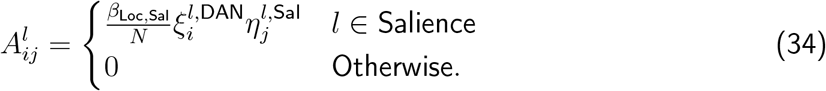

The entries of the patterns 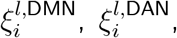 and 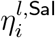 are i.i.d., taking values of −1 and 1 with a probability of 0.5. The strengths of the local symmetric connectivity for the macaque MRNN (Figs. 6b, d-i) are *α*_Loc,DMN_ = *α*_Loc,DMN-FP_ = 2.2, *α*_Loc,DAN_ = *α*_Loc,DAN-FP_ = 2.1, and *β*_Loc,Sal_ = 2.2. The strengths of the local symmetric connectivity for the human MRNN (Figs. 6c, j-o) are *α*_Loc,DMN_ = 2.1, *α*_Loc,DAN_ = 2.6, *α*_Loc,DMN-FP_ = 2.3, *α*_Loc,DAN-FP_ = 1.6, and *β*_Loc,Sal_ = 2.1.

For the long-range projections, the communication subspace (Eq. (2)) is given by

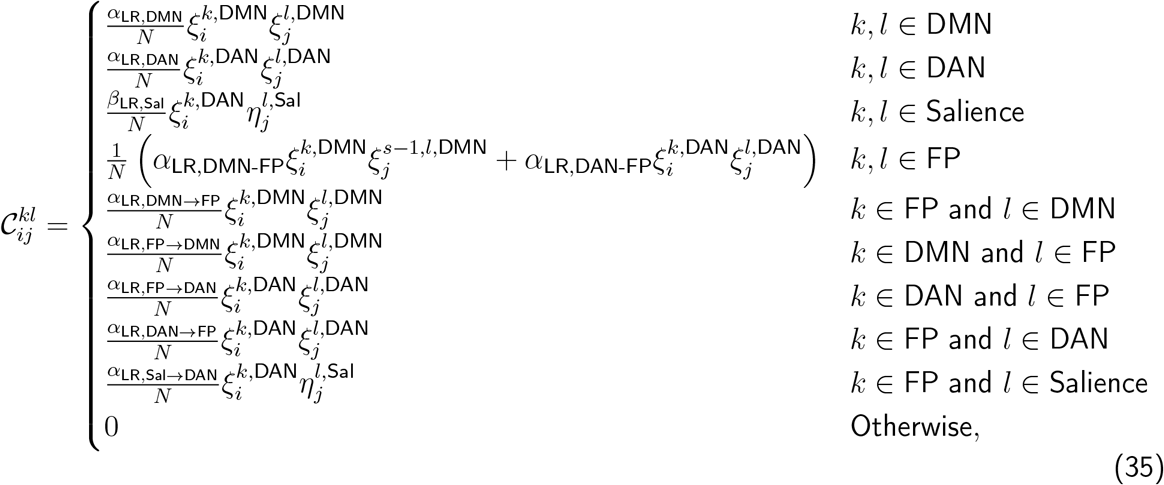

where FP refers to the frontoparietal network. As in Fig. 4 (see Methods 4.4.5), projections within the union of the frontoparietal and DMN have a communication subspace defined by the vectors 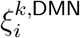, while projections within the union of the frontoparietal and DAN have a communication subspace defined by the vectors 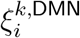. The recurrent connections within the frontoparietal network present both communication subspaces defined by 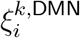 and 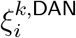. The salience network has an asymmetric-to-asymmetric communication subspace defined by 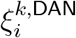 and 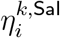. Finally, the salience network projects to the frontoparietal network through an asymmetric-to-symmetric communication subspace, proportional to 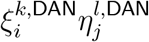. The long-range projections are given by

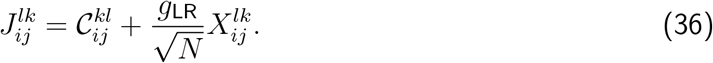

The strengths of the communication subspaces among the cognitive networks in Eq. (35) are listed in Table 3. The strength of the long-range random connectivity is *g*_LR_ = 2.

**Table 3:** Strengths of the communication subspaces across the salience network, frontoparietal network, DMN, and DAN as described in Eq. (36) for the Macaque and Human cortex MRNNs.

| Parameter | Macaque<br>cortex<br>MRNN | Human<br>cortex<br>MRNN |
| --- | --- | --- |
| $\alpha_{\text{LR,DMN}}$ | 7.7 | 7.35 |
| $\alpha_{\text{LR,DAN}}$ | 7.35 | 9.1 |
| $\beta_{\text{LR,SaI}}$ | 7.7 | 7.35 |
| $\alpha_{\text{LR,DMN-FP}}$ | 7.7 | 8.05 |
| $\alpha_{\text{LR,FP} \rightarrow \text{DMN}}$ | 7.7 | 8.05 |
| $\alpha_{\text{LR,DMN} \rightarrow \text{FP}}$ | 7.7 | 8.05 |
| $\alpha_{\text{LR,DAN-FP}}$ | 7.35 | 5.6 |
| $\alpha_{\text{LR,FP} \rightarrow \text{DAN}}$ | 7.35 | 7 |
| $\alpha_{\text{LR,DAN} \rightarrow \text{FP}}$ | 7.35 | 4.2 |
| $\alpha_{\text{LR,SaI} \rightarrow \text{FP}}$ | 15.5 | 15.5 |

To investigate the transition between the FP-DMN and FP-DAN attractor states mediated by the salience network, we introduced a transient input to the salience network areas given by

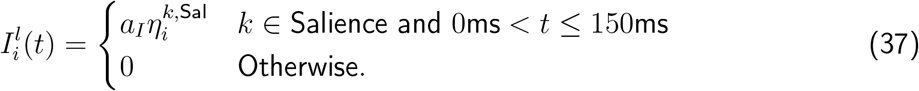

The magnitude of the external input current is *a*_*I*_ = 5 for Fig. 4 and the corresponding supplementary figures (Figs. S18-S21).

### 4.5 Mean field theory

In Fig. 2e, we present the result of a mean field analysis of the network in the limit of an infinite number of units per area, i.e., *N* → ∞. The connectivity of the network in this case is given by Eqs.(10–13). Let us first define, the recurrent input current, 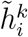, to a neuron *i* in area *k*, which is given by

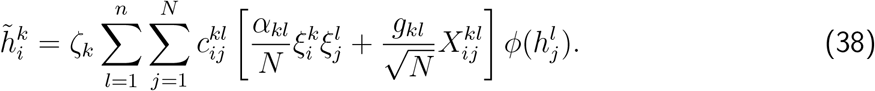

We define the overlap in area *k* as the covariance between the population activity of that area and the pattern 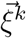, which generates the symmetric connectivity in Eqs.(10–13). Therefore, the overlap is given by

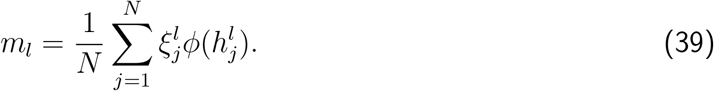

For building a mean field theory for the dynamics of our network, we first compute the average input current to a neuron *i* in area *k* conditioned on the pattern 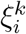. The average is taken over the randomness of the anatomical connectivity 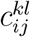 (see Eq. (4)) and the random connectivity motif 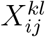, represented by the triangular brackets ⟨· · · ⟩, obtaining

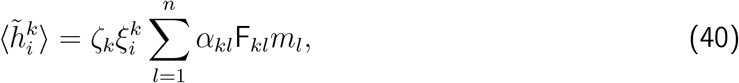

with *α*_*kk*_ = *α*_Loc_ and *α*_*kl*_ = *α*_LR_ for *l* ≠ *k*.

Here F_*kl*_ is the normalized dMRI tractography matrix (see Methods 4.1). The variance of the input current is given by

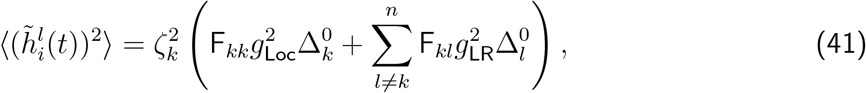

with *g*_*kk*_ = *g*_Loc_ and *g*_*kl*_ = *g*_LR_ for *l* ≠ *k*, and where

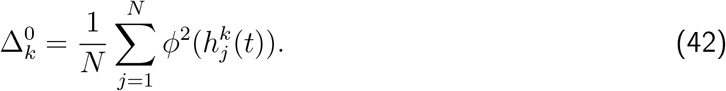

In the above calculations, we assume that the network reaches a stationary state in which the activity for each unit does not vary in time, and the network reaches a stable fixed-point attractor. This is not the case for Fig. 2, as the dynamics is chaotic. However, our computations for the overlap provide a good approximation in this chaotic regime (see Fig. 2e). Numerically solving a dynamic mean-field theory, along the lines of what is presented in [96, 113], is necessary to accurately describe the chaotic phase in our MRNNs.

In the limit where there are many units per area, we can express the stationary input current in area *k* as

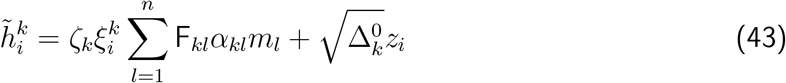

The variable *z*_*i*_ is an i.i.d. Gaussian random variable with mean zero and unit variance. This random variable represents the neuron-by-neuron variability of the input current and is assumed to be Gaussian, based on the fact that in large networks, the variability is approximately Gaussian distributed. Then, we compute self-consistent equations for the area-specific order parameters in our networks which are the overlaps *m*_*k*_ and the variance of the input current 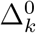, which describe the macroscopic behavior of the network in the stationary state. In the large *N* limit, Eq.(39) becomes an integral. Using the expression for the input current from Eq.(43), we obtain an equation for the overlaps

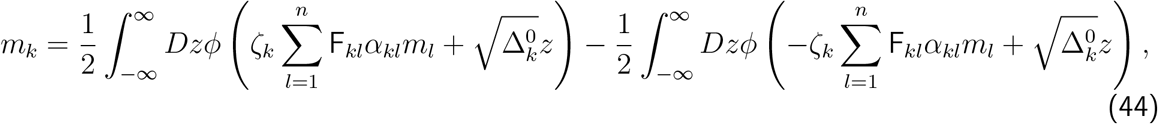

where 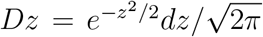. Similarly, we use Eqs. (42) and Eq.(43) for obtaining a self-consistent equation for the input currents is given by

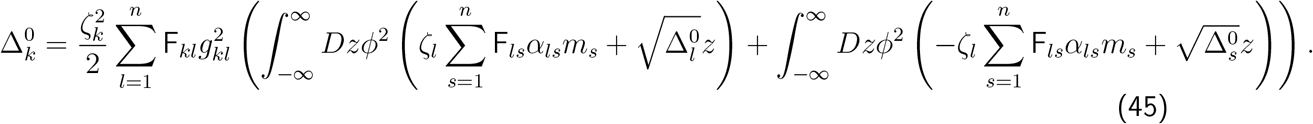

Finally, we obtain 89 self-consistent equations for the overlaps, given by Eqs.(44), and 89 for the variances of the input current, given by Eqs.(45). These equations are coupled. We solved these self-consistent equations using a custom-made code implemented with the Python package PyTorch (see Section 5).

## 5 Code availability

All code used to analyze the connectivity, simulate the networks, solve the mean field equations, and generate figures are available at https://github.com/ulisespereira/brainwide.

## 6. Acknowledgement

We would like to thank Vishwa Goudar for the valuable discussions at the early stages of this research. We thank Łukasz Kuśmierz, Shawn Olsen, Daniel Margulies, Ulysse Klatzmann, Aswathi Thrivikraman for their constructive comments on the manuscript. This work was supported by Swartz Foundation postdoctoral fellowship (to U.P.-O.); Biotechnology and Biological Sciences Research Council (UKRI) grant BB/X013243/1 to S.F.W.; the James Simons Foundation Grant 543057SPI, the NSF Neuronex grant 2015276 and National Institutes of Health grant R01MH062349 (to X.-J.W.).

## 7 Author contributions

U.P.-O., S.F.-W. and X.-J.W. designed the research; U.P.-O. and S.F.-W. performed the research with inputs from X.-J.W; U.P.-O., S.F.-W., and X.-J.W.wrote the paper.

## Supplementary Information

### Supplementary figures

**Fig. S1:**
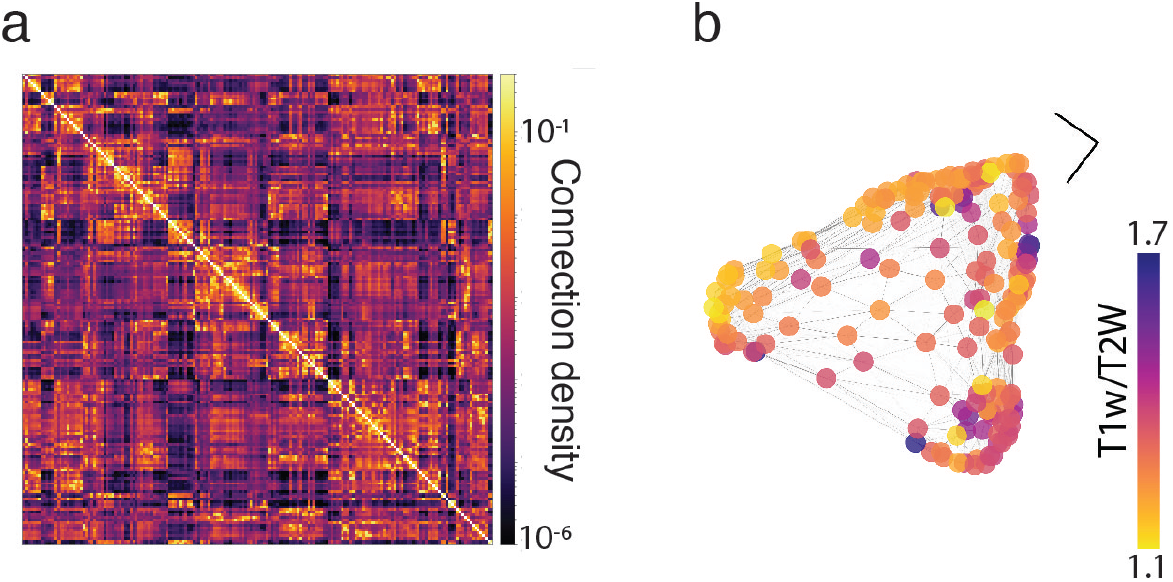
Supplementary figure corresponding to Fig. 1h and j in the main text. (a) dMRI tractography of the human cortex parcellated into 180 cortical regions [130]. (b) The T1w/T2w measure across the human cortex [36] and dMRI connectivity graph. The edges are proportional to the value of the dMRI matrix. The nodes are colored according to the T1w/T2w value. The nodes are placed using the diffusion map algorithm (Methods 4.2).

**Fig. S2:**
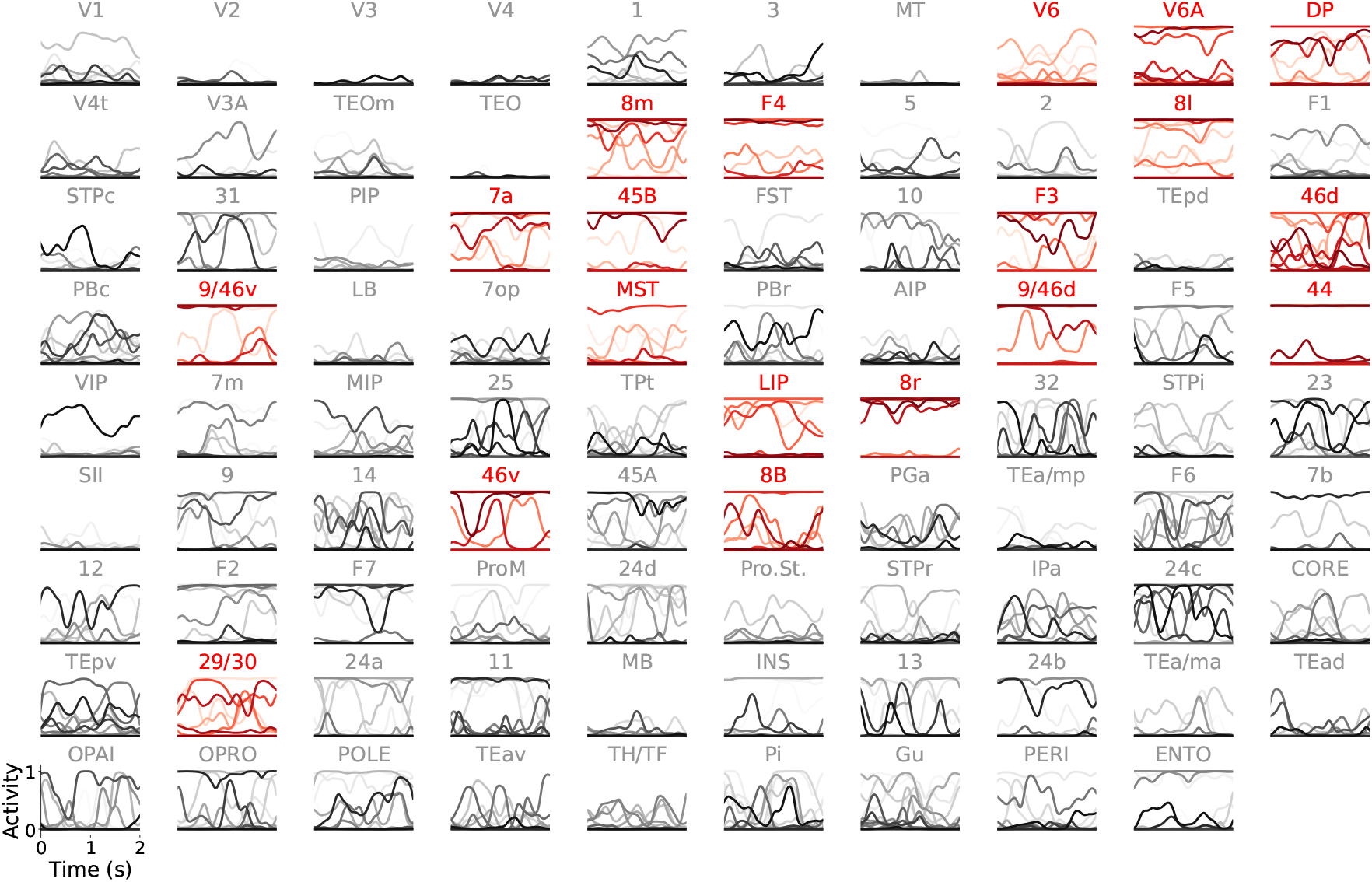
Supplementary figure corresponding to Fig. 2 in the main text. Each panel displays the dynamics of 15 representative units from each of the 89 cortical areas in the macaque cortex. Areas with red labels and traces belong to the frontoparietal network.

**Fig. S3:**
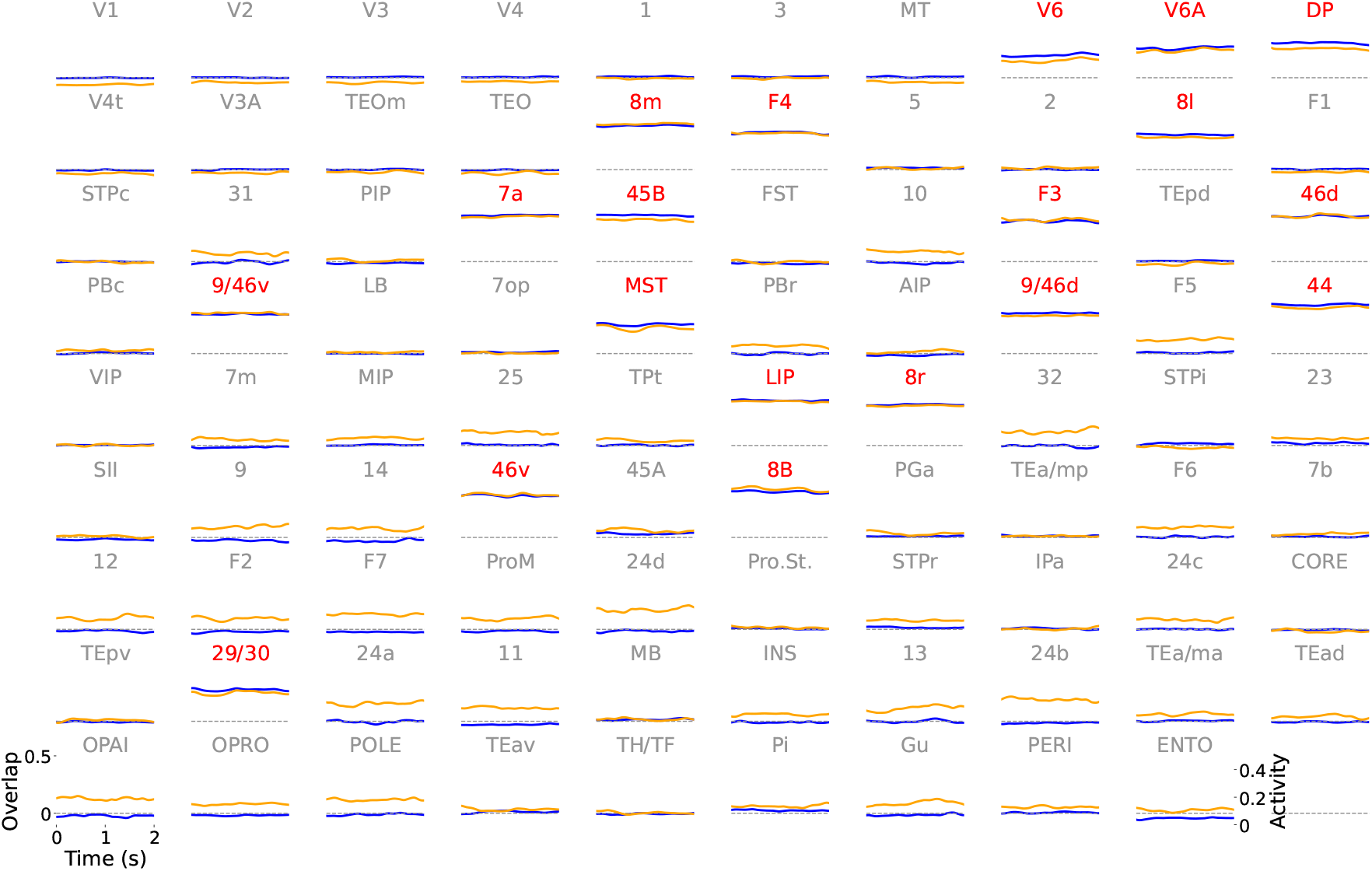
Supplementary figure corresponding to Fig. 2 in the main text. Each panel displays the overlap (in blue) and mean activity (in yellow) for each of the 89 cortical areas in the macaque cortex. Areas with red labels and traces belong to the frontoparietal network.

**Fig. S4:**
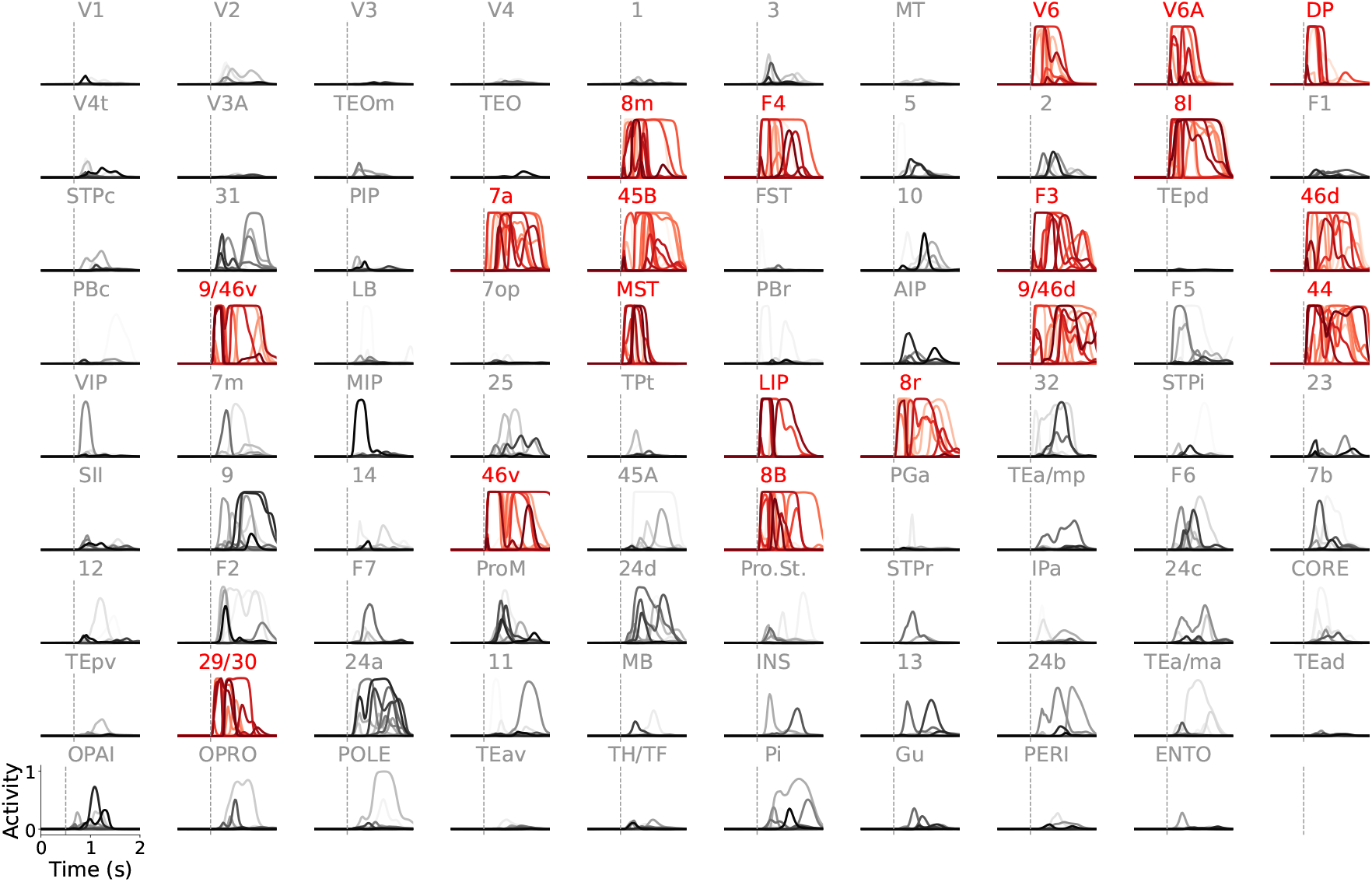
Supplementary figure corresponding to Fig. 3 in the main text. Each panel displays the dynamics of 15 representative units from each of the 89 cortical areas in the macaque cortex. Areas with red labels and traces belong to the frontoparietal network. We stimulate all areas in the frontoparietal network with a 150ms input current aligned with the first pattern in the sequence at 0.5s in our simulation. The onset of the stimulus is marked by a vertical dashed line.

**Fig. S5:**
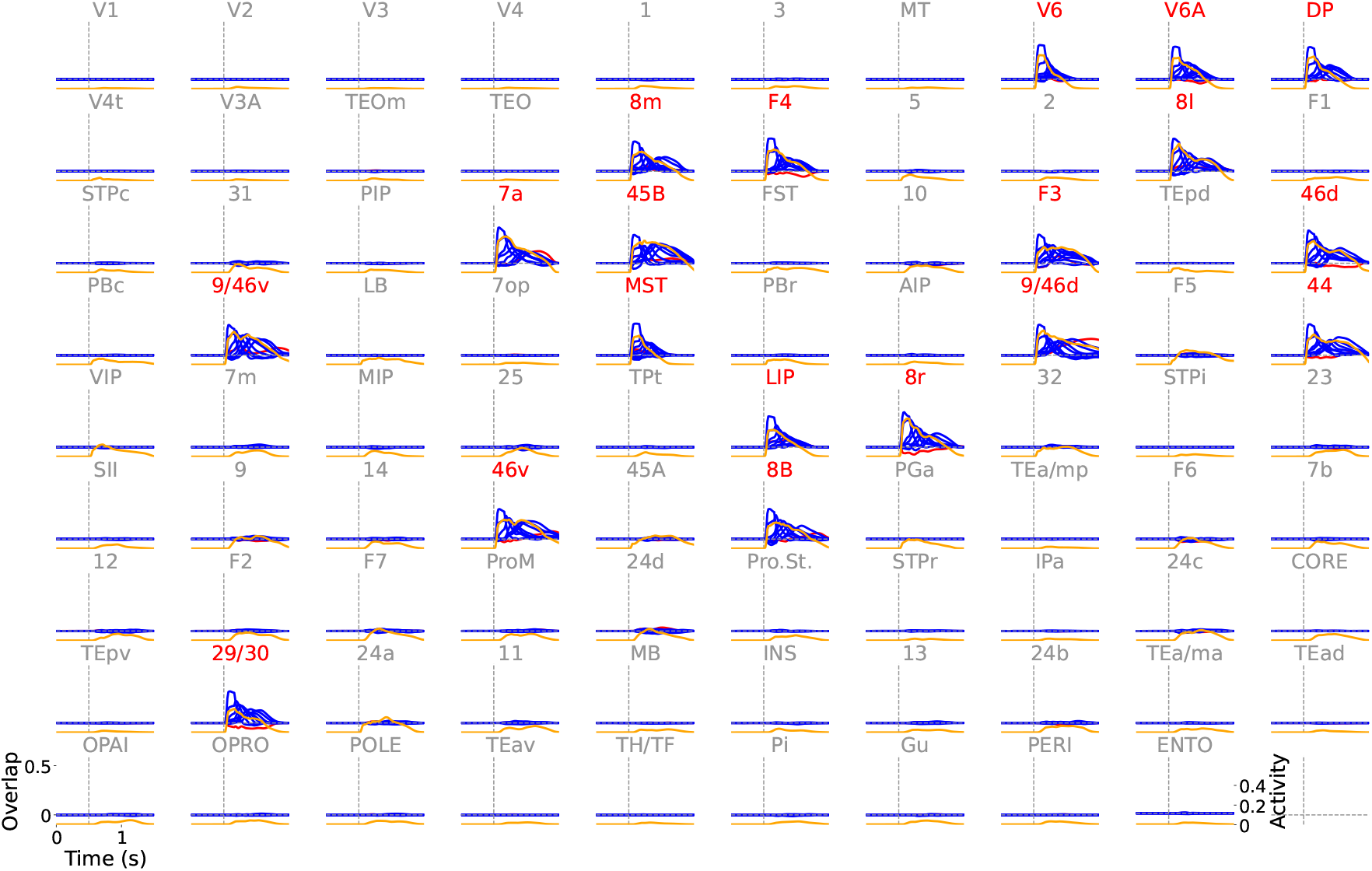
Supplementary figure corresponding to Fig. 3 in the main text. Each panel displays the overlaps (in blue) and mean activity (in yellow) for each of the 89 cortical areas in the macaque cortex. Areas with red labels and traces belong to the frontoparietal network. We stimulate all areas in the frontoparietal network with a 150ms input current aligned with the first pattern in the sequence at 0.5s in our simulation. The onset of the stimulus is marked by a vertical dashed line.

**Fig. S6:**
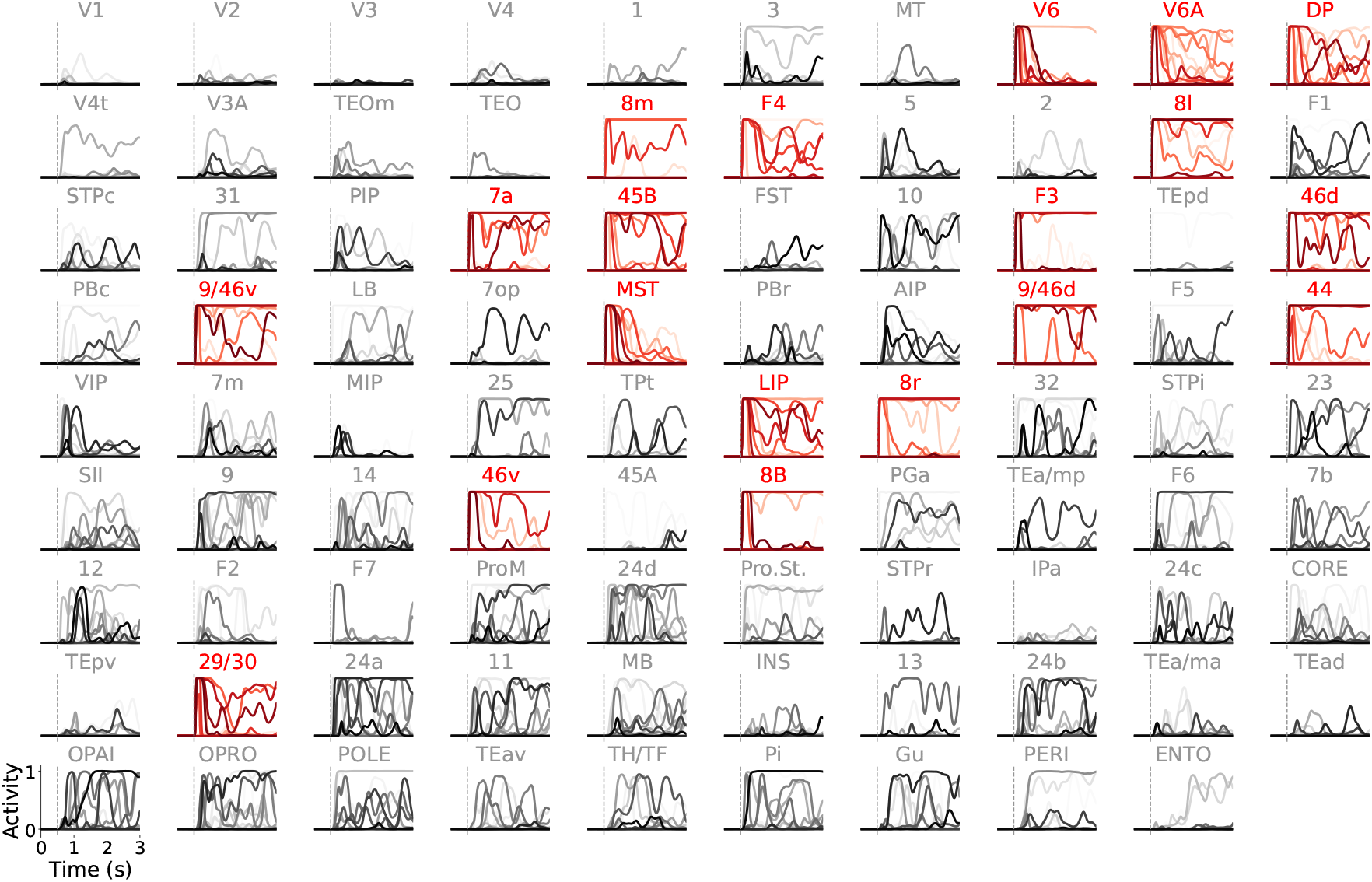
Supplementary figure corresponding to Fig. 3 in the main text. Each panel displays the dynamics of 15 representative units from each of the 89 cortical areas in the macaque cortex. Areas with red labels and traces belong to the frontoparietal network. We stimulate the frontoparietal network with a 150 ms stimulus, starting at 0.5 seconds into the simulation (the onset is marked by a vertical dashed line), aligned with the symmetric-to-symmetric communication subspace.

**Fig. S7:**
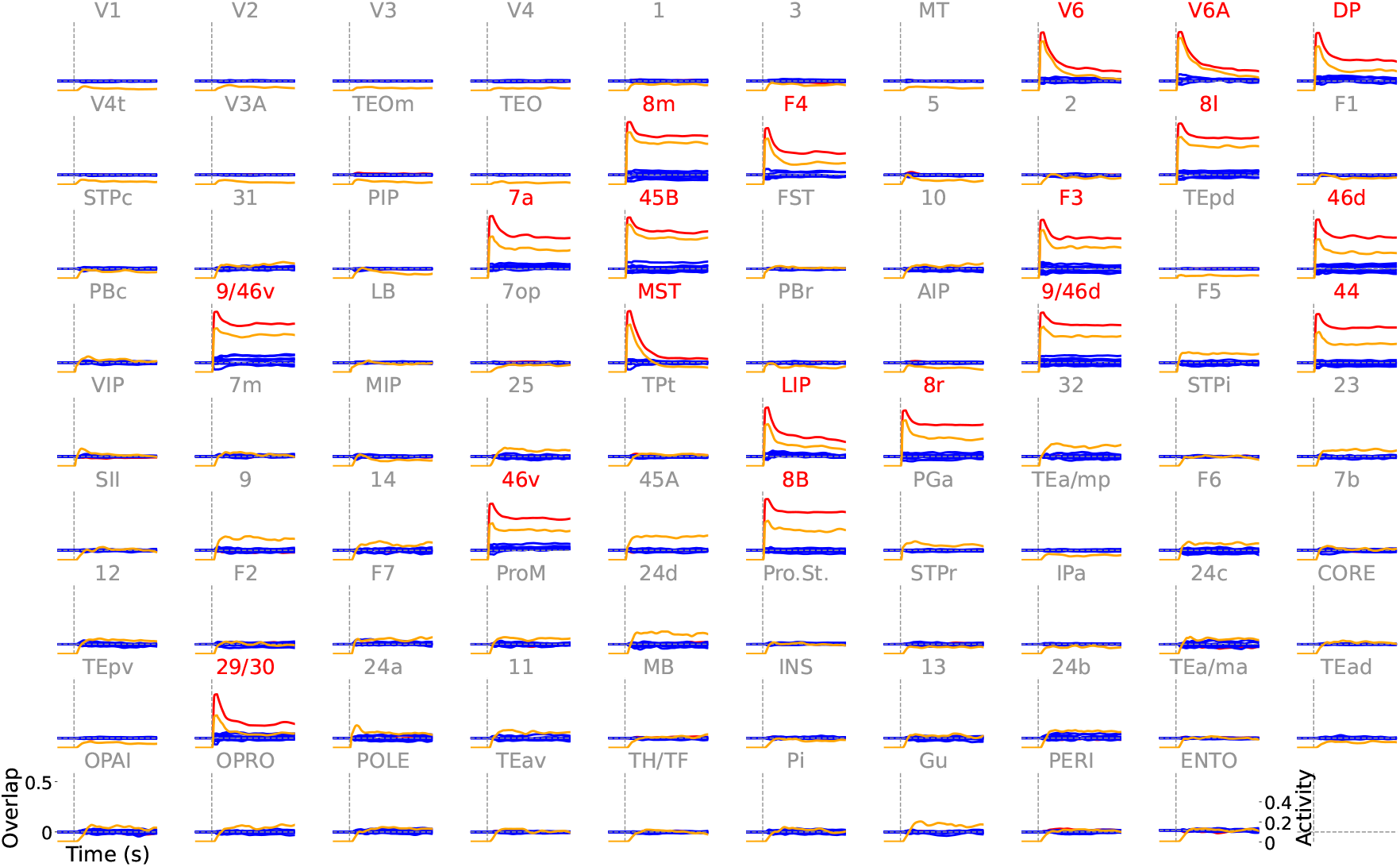
Supplementary figure corresponding to Fig. 3 in the main text. Each panel displays the overlap with the patterns comprising the symmetric connectivity (red), asymmetric connectivity (blue), and mean activity (yellow) for each of the 89 cortical areas in the macaque cortex. Areas with red labels and traces belong to the frontoparietal network. We stimulate the frontoparietal network with a 150 ms stimulus, starting at 0.5 seconds into the simulation (the onset is marked by a vertical dashed line), aligned with the symmetric-to-symmetric communication subspace.

**Fig. S8:**
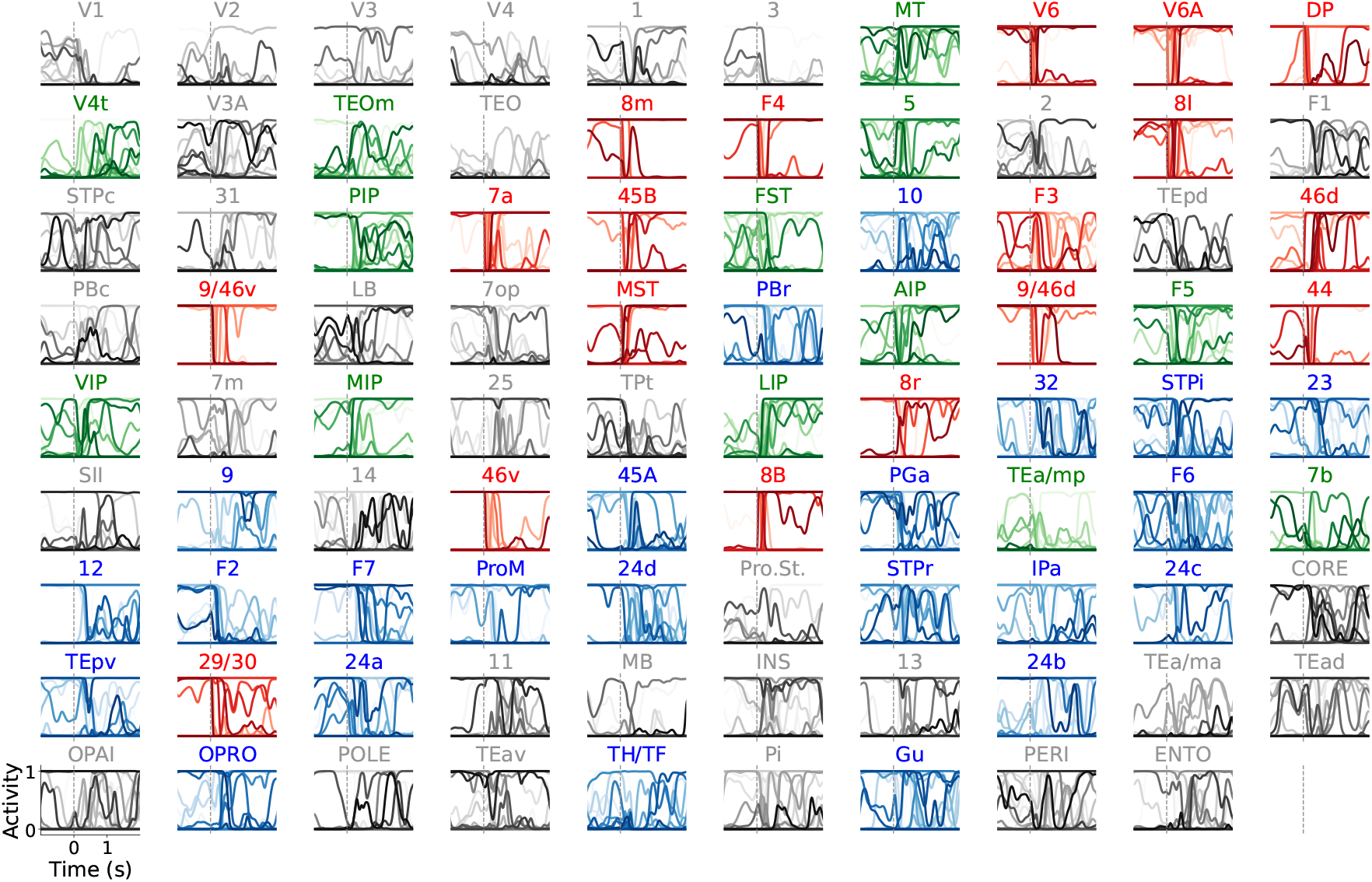
Supplementary figure corresponding to Fig. 4 in the main text. Neural dynamics for 15 representative units. Blue-labeled areas and activity traces correspond to the DMN, green-labeled areas and traces correspond to the DAN, and red-labeled areas and traces correspond to the frontoparietal network. The gray vertical dashed line represents the moment when an external constant stimulus switches its alignment from the FP-DMN to the FP-DAN attractor.

**Fig. S9:**
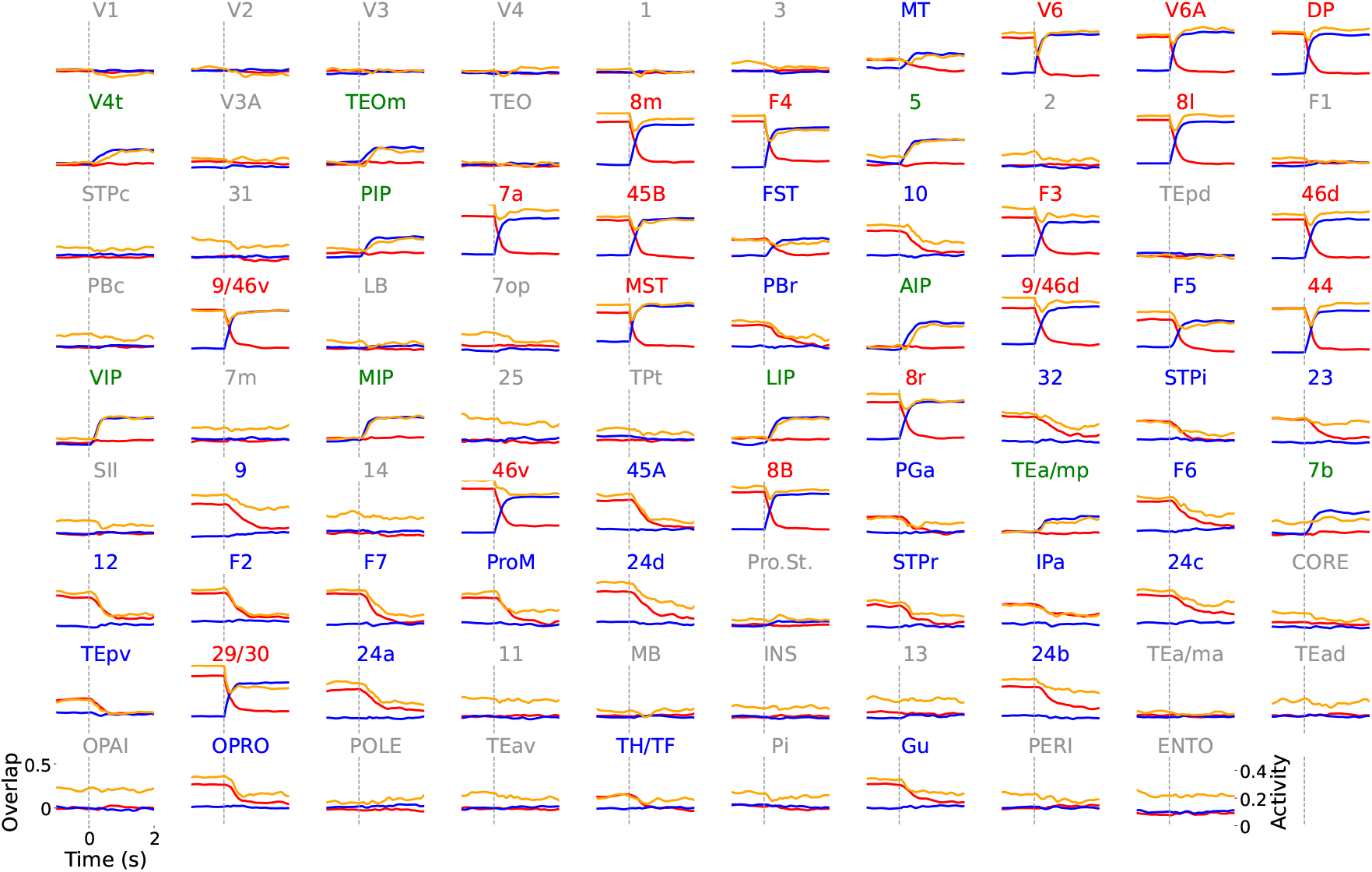
Supplementary figure corresponding to Fig. 4 in the main text. Blue represents the FP-DMN attractor, green represents the FP-DAN attractor, and yellow indicates the mean activity (bottom right axis). Areas labeled in blue, green, and red correspond to the DMN, DAN, and frontoparietal network, respectively. The gray vertical dashed line marks the moment when an external constant stimulus shifts its alignment from the FP-DMN to the FP-DAN attractor.

**Fig. S10:**
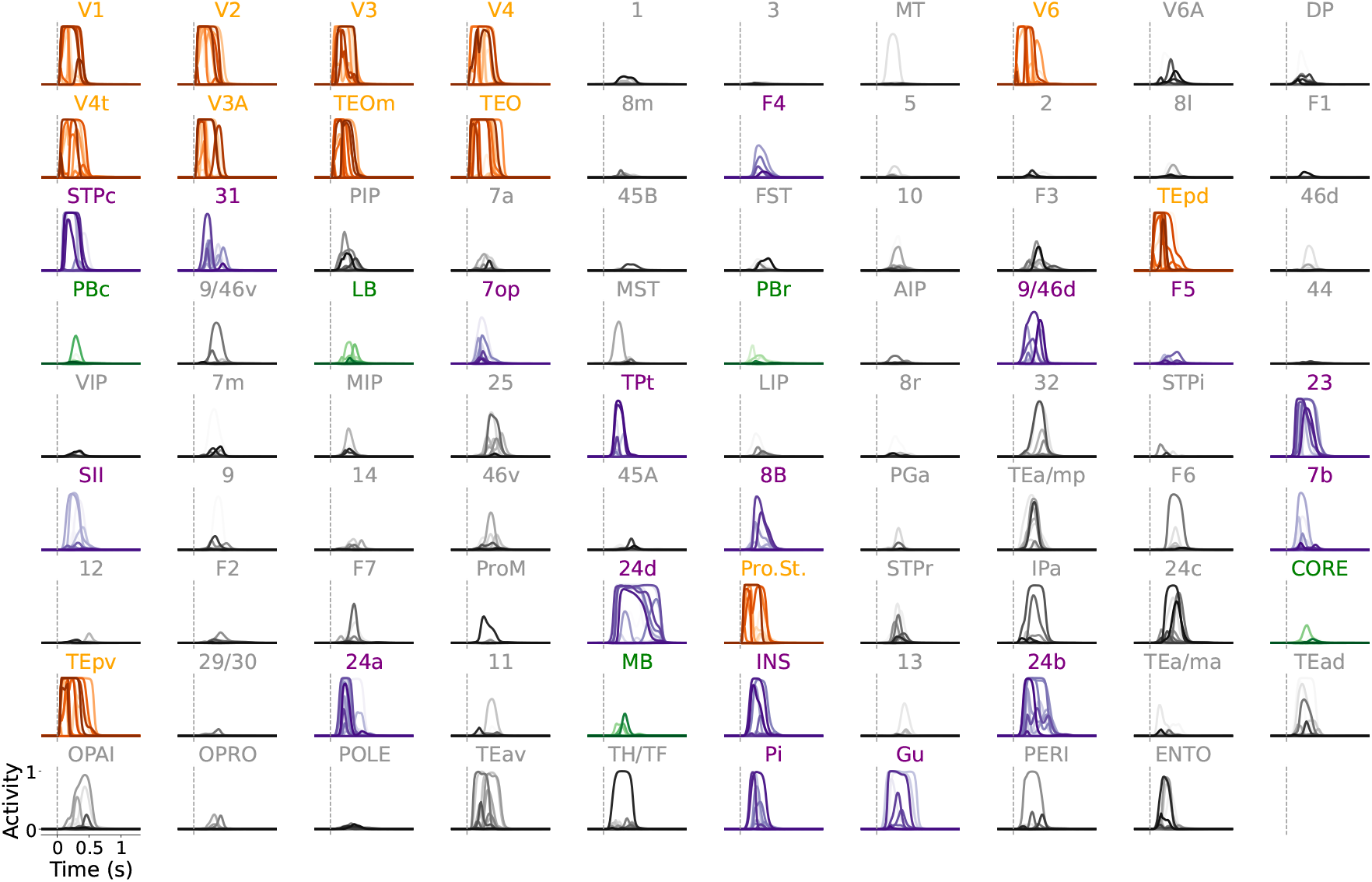
Supplementary figure corresponding to Fig. 5f in the main text. Network response to a **salient visual stimulus**. Neural dynamics for 15 representative units. Areas with yellow labels and red traces belong to the visual network. Areas with green labels and traces belong to the auditory network. Areas with purple labels and traces belong to the salience network.

**Fig. S11:**
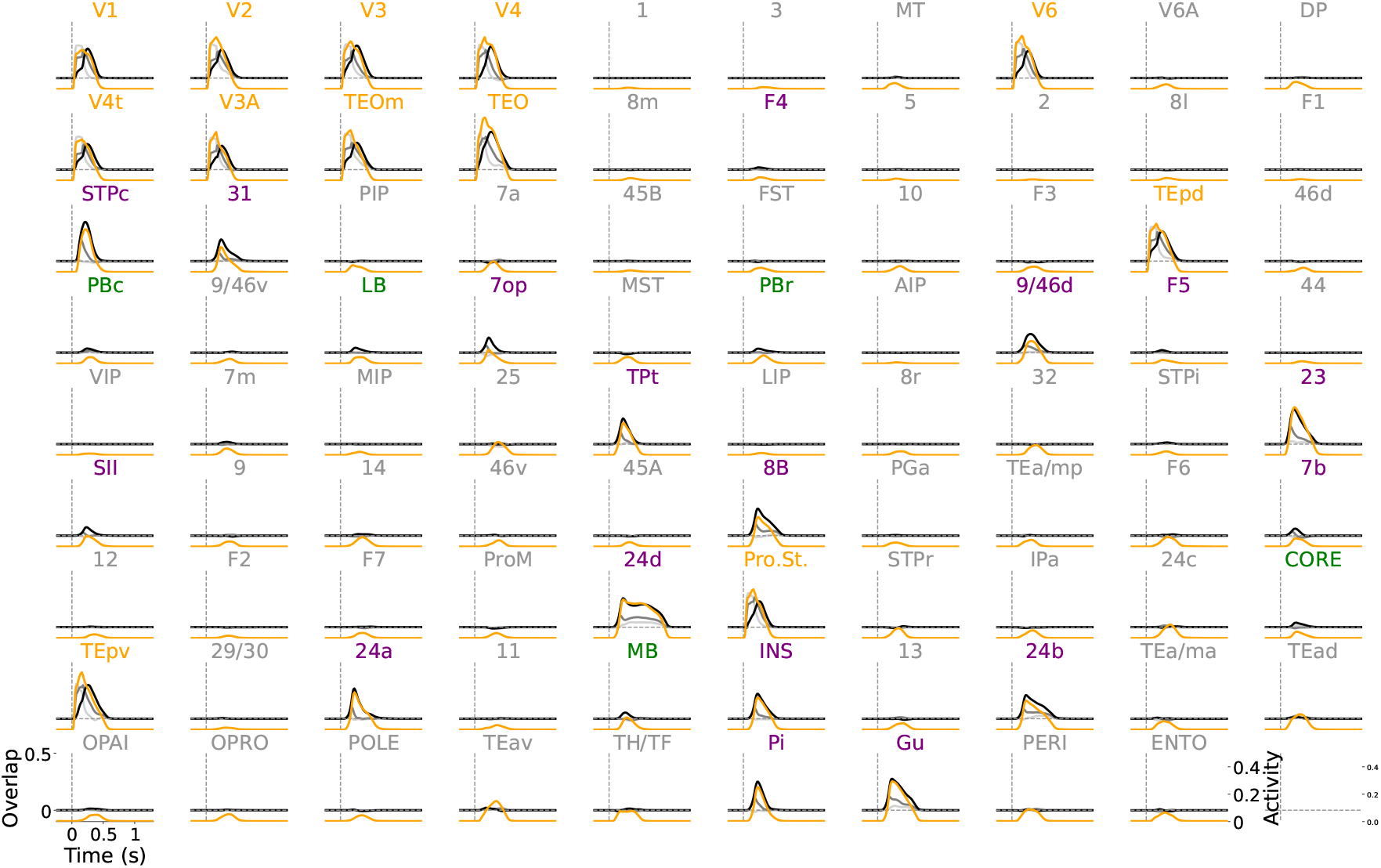
Supplementary figure corresponding to Fig. 5f in the main text. Network response to a **salient visual stimulus**. Each panel shows the overlaps (in gray) and the mean activity (in yellow) for all 89 cortical areas in the macaque cortex. Areas labeled in yellow, green, and purple correspond to the visual, auditory, and salience networks, respectively.

**Fig. S12:**
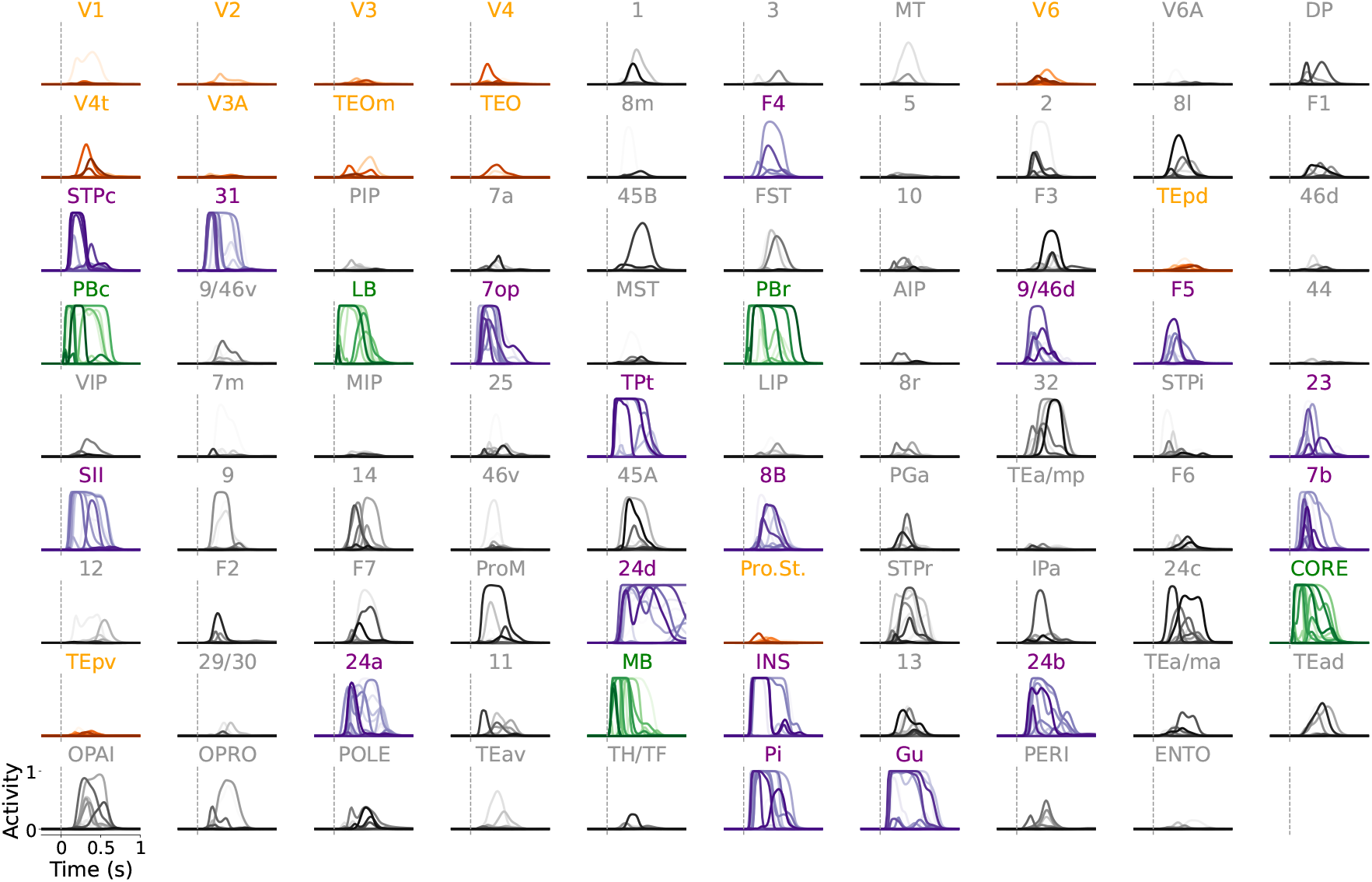
Supplementary figure corresponding to Fig. 5h in the main text. Network response to a **salient auditory stimulus**. Neural dynamics for 15 representative units. Areas with yellow labels and red traces belong to the visual network. Areas with green labels and traces belong to the auditory network. Areas with purple labels and traces belong to the salience network.

**Fig. S13:**
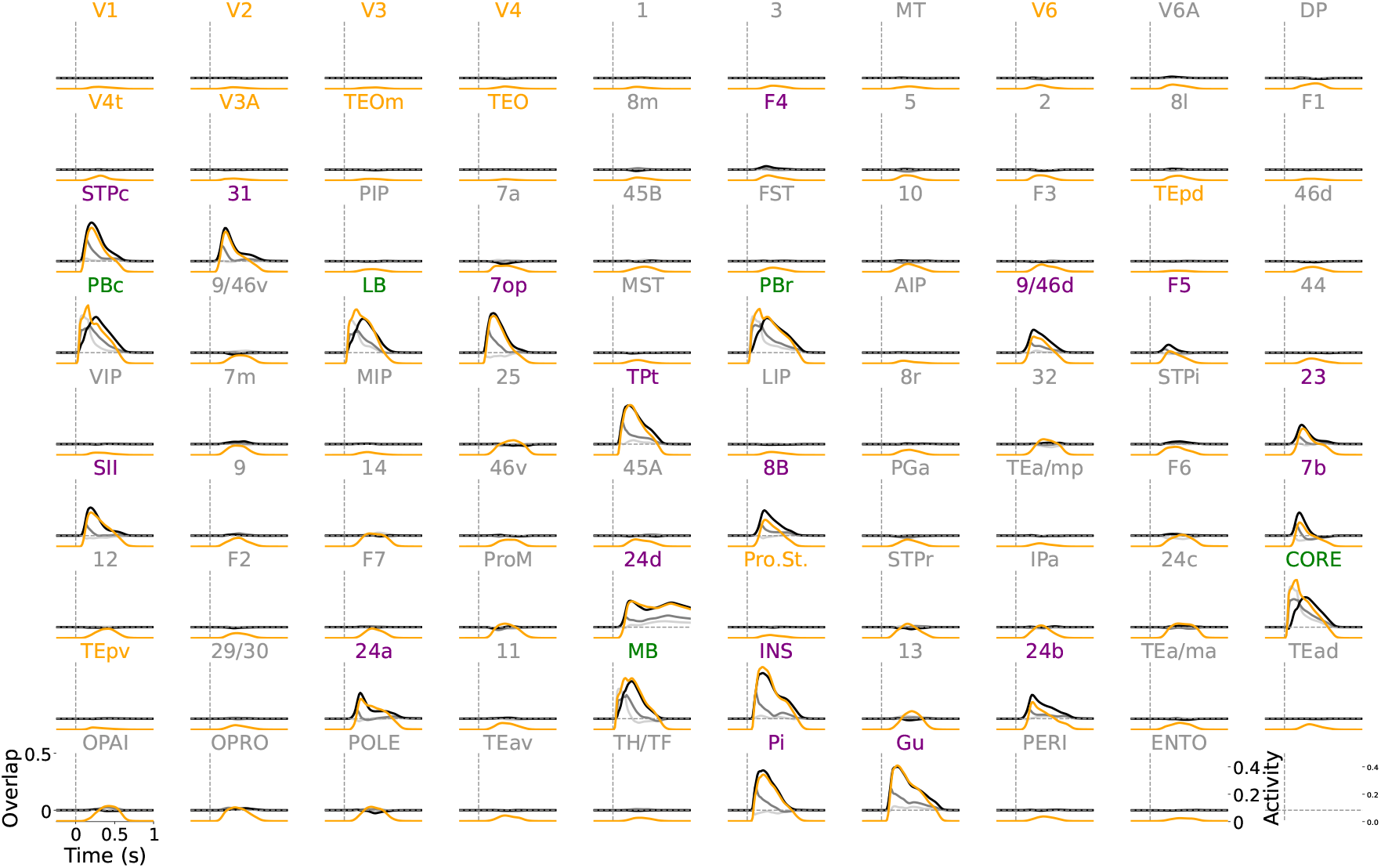
Supplementary figure corresponding to Fig. 5h in the main text. Network response to a **salient auditory stimulus**. Each panel shows the overlaps (in gray) and the mean activity (in yellow) for all 89 cortical areas in the macaque cortex. Areas labeled in yellow, green, and purple correspond to the visual, auditory, and salience networks, respectively.

**Fig. S14:**
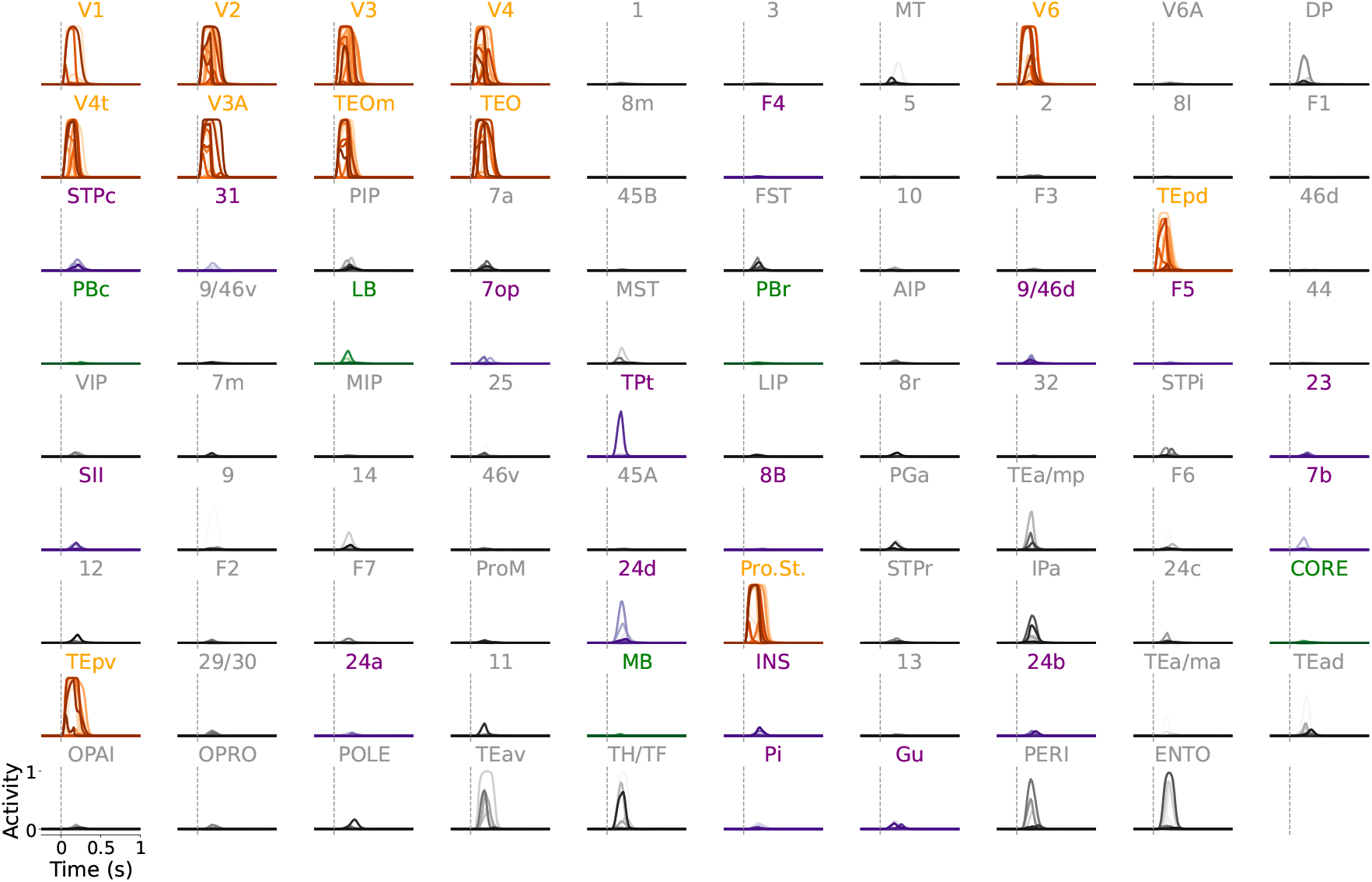
Supplementary figure corresponding to Fig. 5j in the main text. Network response to **a non-salient visual stimulus**. Neural dynamics for 15 representative units. Areas with yellow labels and red traces belong to the visual network. Areas with green labels and traces belong to the auditory network. Areas with purple labels and traces belong to the salience network.

**Fig. S15:**
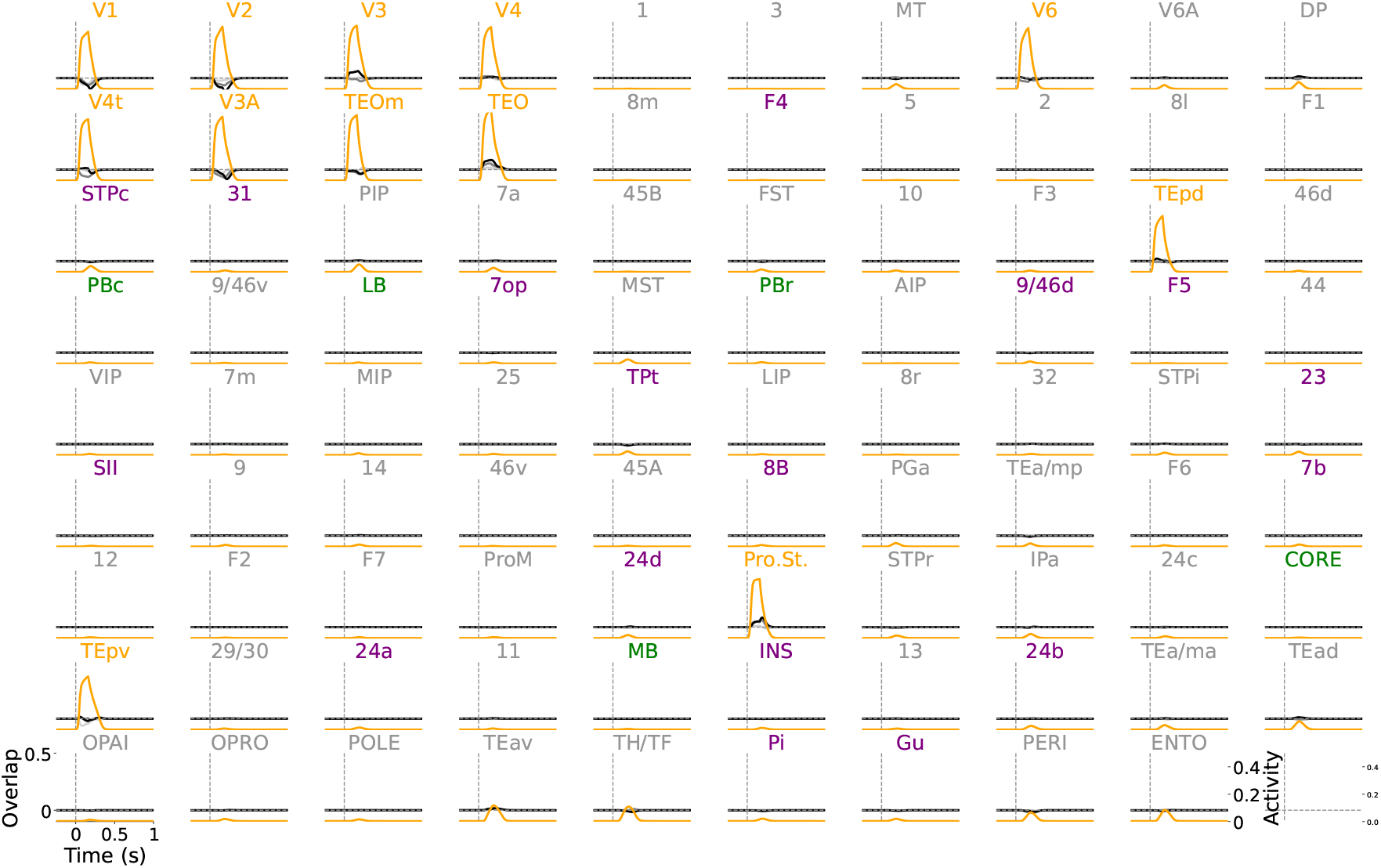
Supplementary figure corresponding to Fig. 5j in the main text. Network response to a **non-salient visual stimulus**. Each panel shows the overlaps (in gray) and the mean activity (in yellow) for all 89 cortical areas in the macaque cortex. Areas labeled in yellow, green, and purple correspond to the visual, auditory, and salience networks, respectively.

**Fig. S16:**
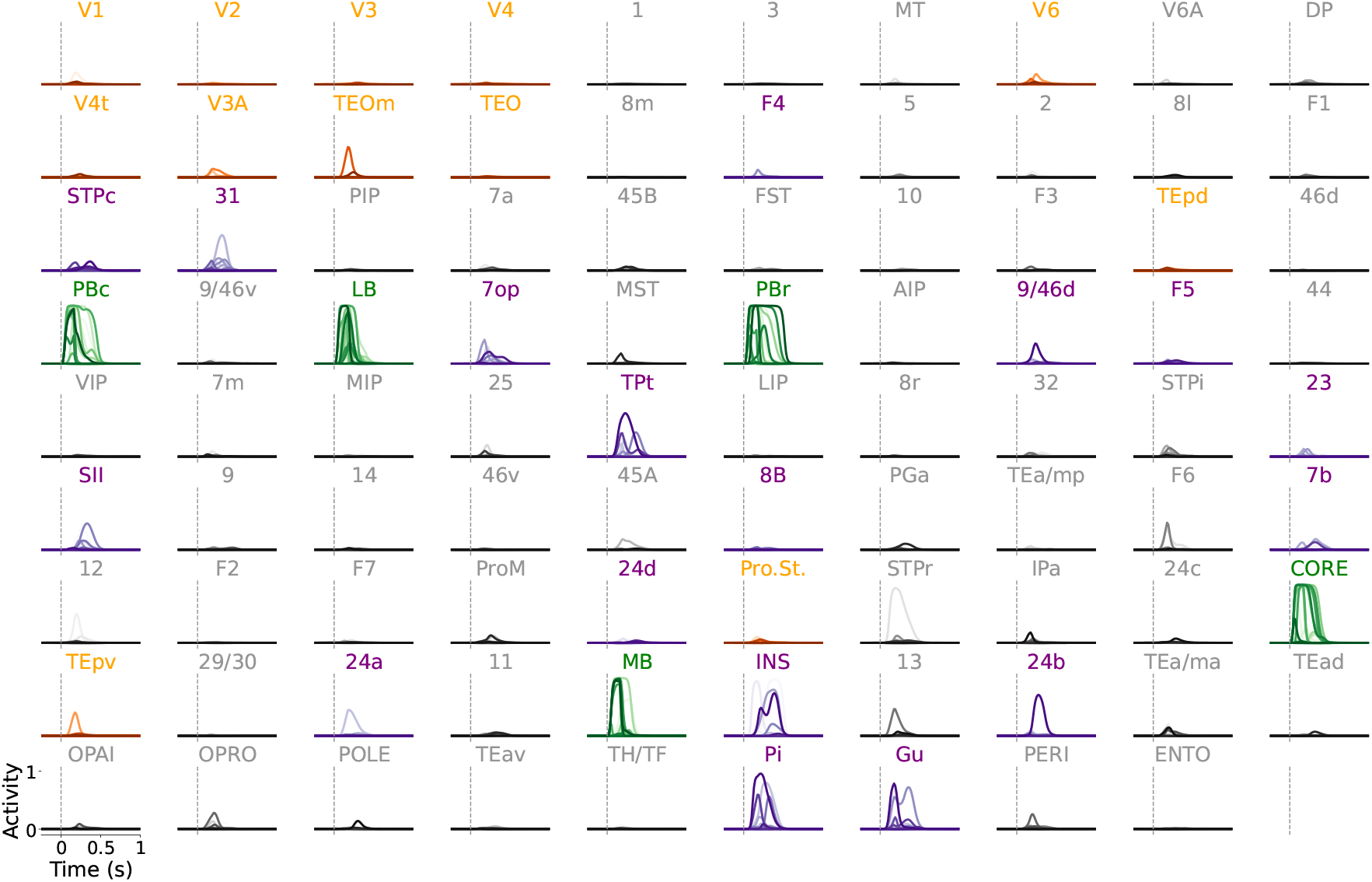
Supplementary figure corresponding to Fig. 5j in the main text. Network response to a **non-salient auditory stimulus**. Neural dynamics for 15 representative units. Areas with yellow labels and red traces belong to the visual network. Areas with green labels and traces belong to the auditory network. Areas with purple labels and traces belong to the salience network.

**Fig. S17:**
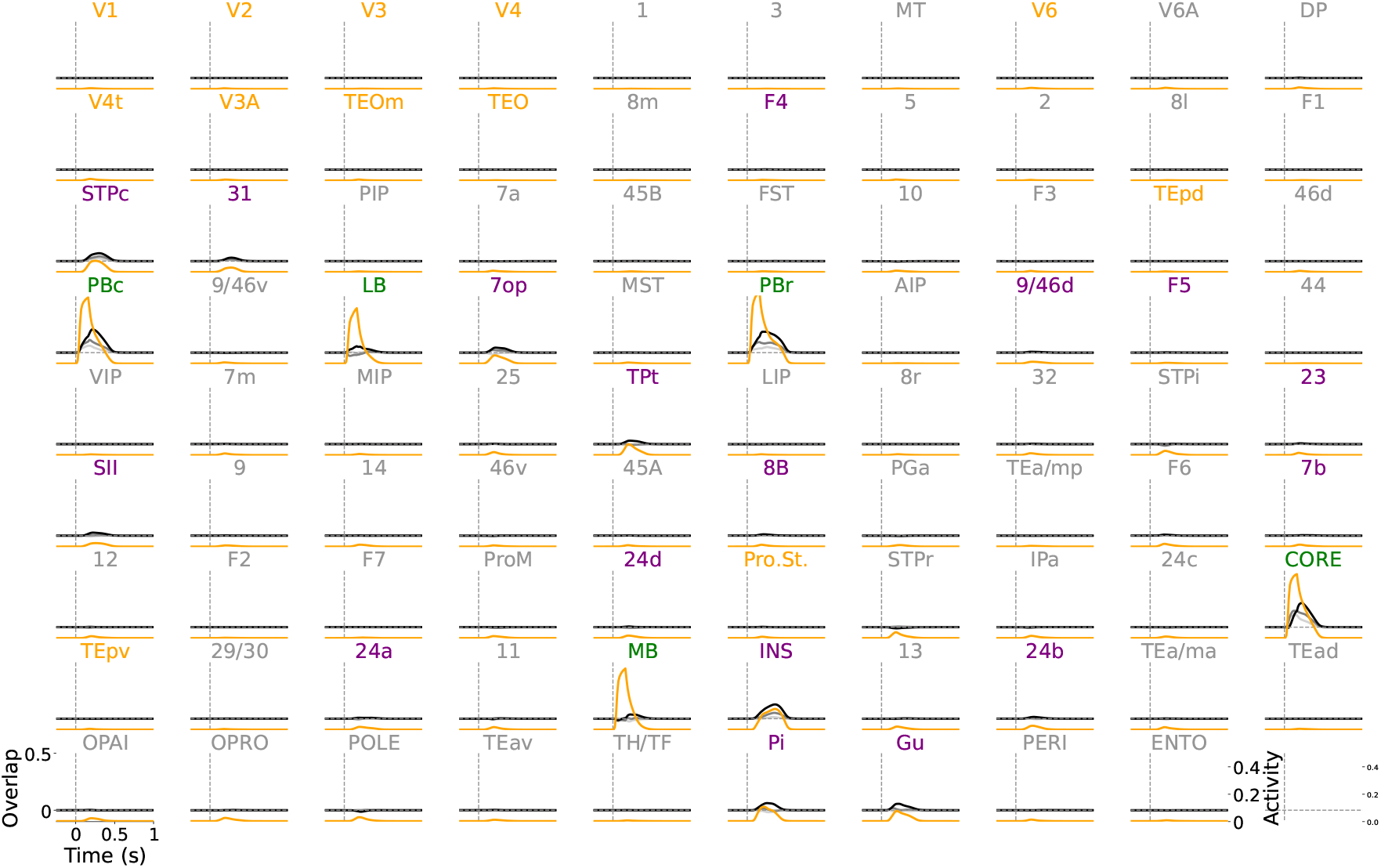
Supplementary figure corresponding to Fig. 5 in the main text. Network response to a **non-salient auditory stimulus**. Each panel shows the overlaps (in gray) and the mean activity (in yellow) for all 89 cortical areas in the macaque cortex. Areas labeled in yellow, green, and purple correspond to the visual, auditory, and salience networks, respectively.

**Fig. S18:**
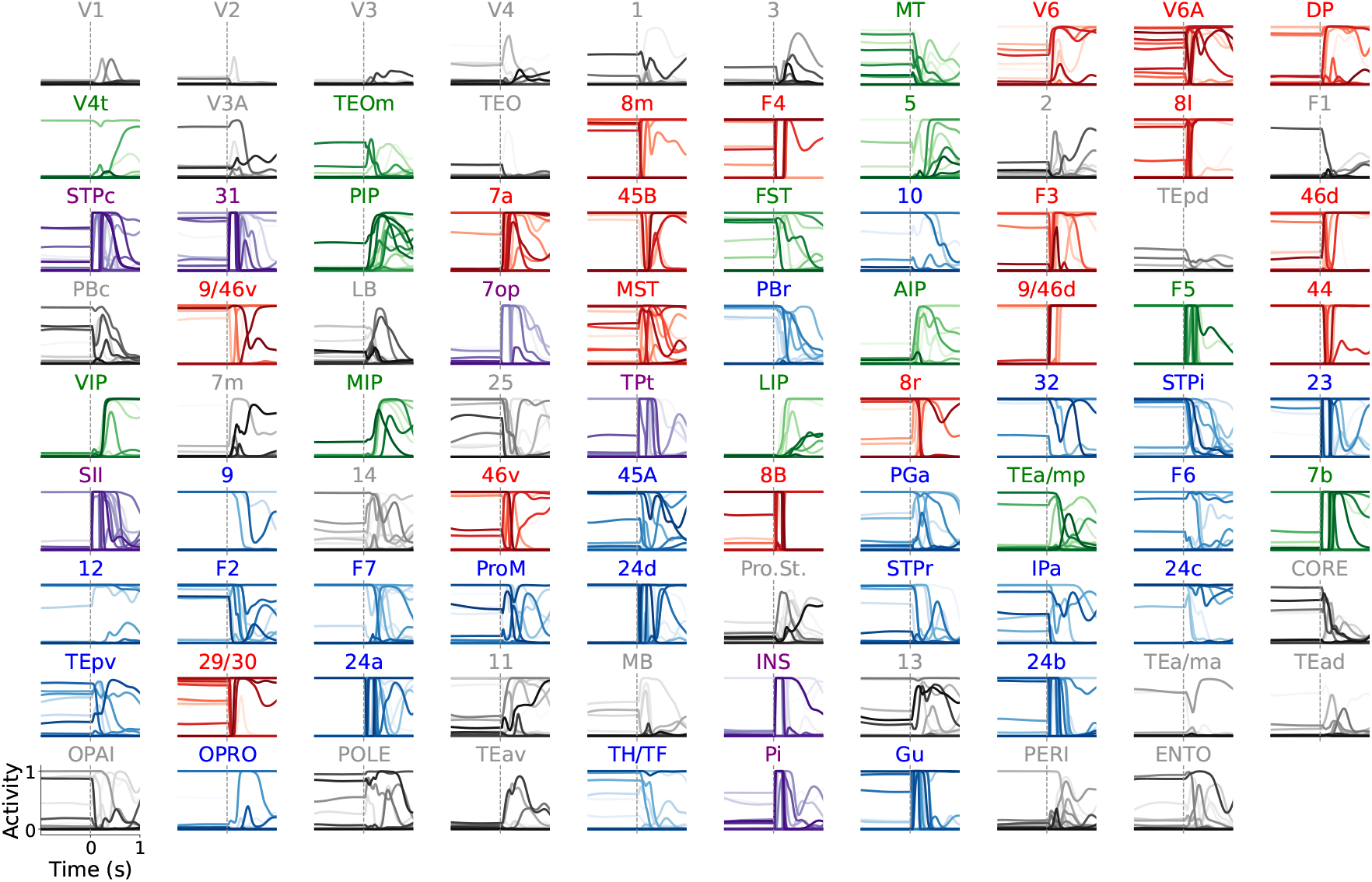
Supplementary figure corresponding to Fig. 6b in the main text. Network response to a 150ms stimulus to areas in the salience network. Neural dynamics for 15 representative units. Areas with purple labels and traces belong to the salience network. Areas with red labels and traces belong to the frontoparietal network. Areas with green labels and traces belong to the DAN. Areas with blue labels and traces belong to the DAN.

**Fig. S19:**
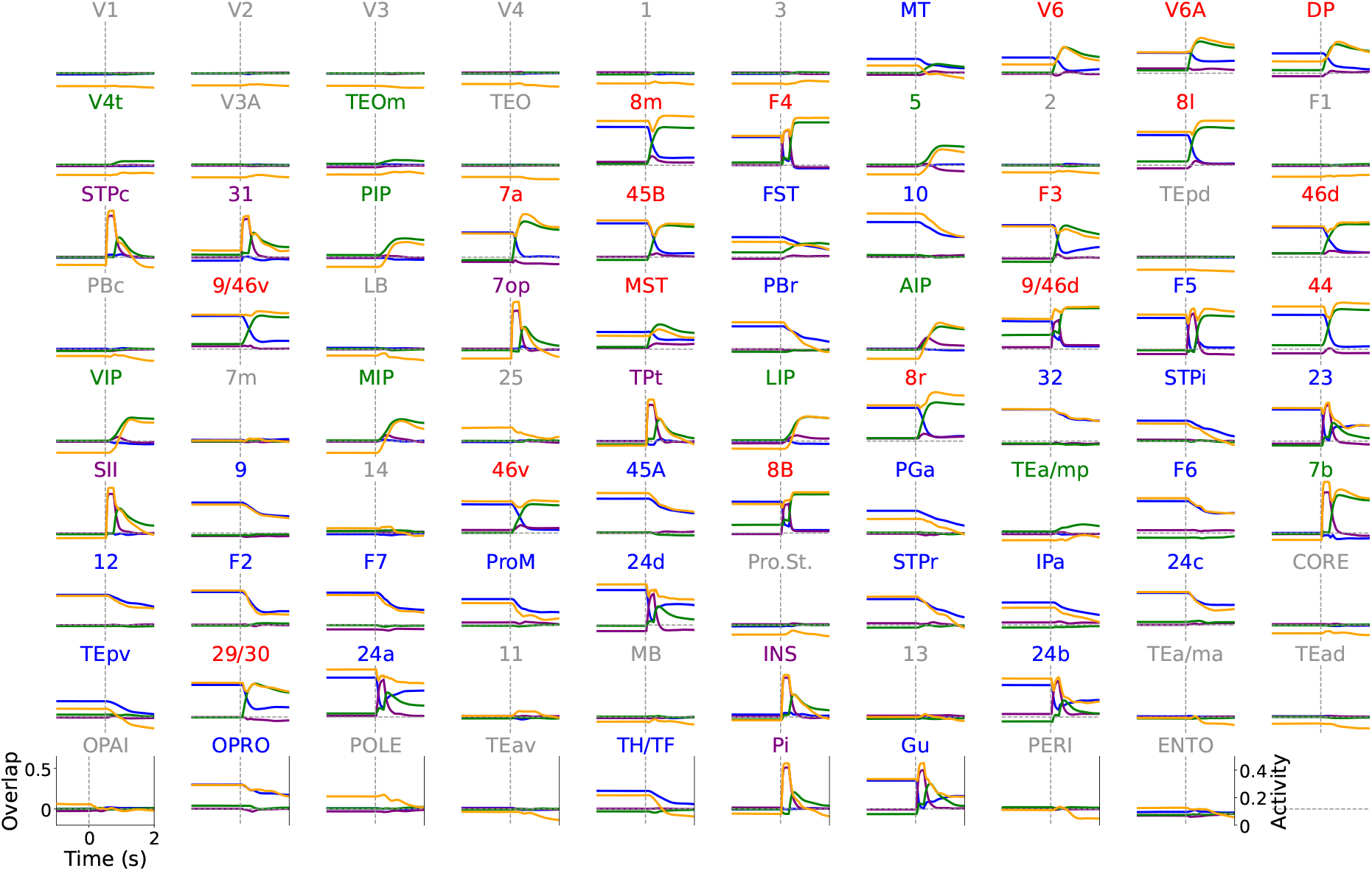
Supplementary figure corresponding to Fig. 6b, d-i in the main text. Network response to a 150ms stimulus to areas in the salience network. Blue: overlap FP-DMN attractor; Green: overlap FP-DAN attractor; Purple: overlap with the pattern stimulated in the salience network. The gray dashed line marks the transient stimulation of salience areas at 0s for 150 ms. Purple labels and traces: Salience network. Red: Frontoparietal network. Green: Dorsal Attention Network (DAN). Blue: Visual network.

**Fig. S20:**
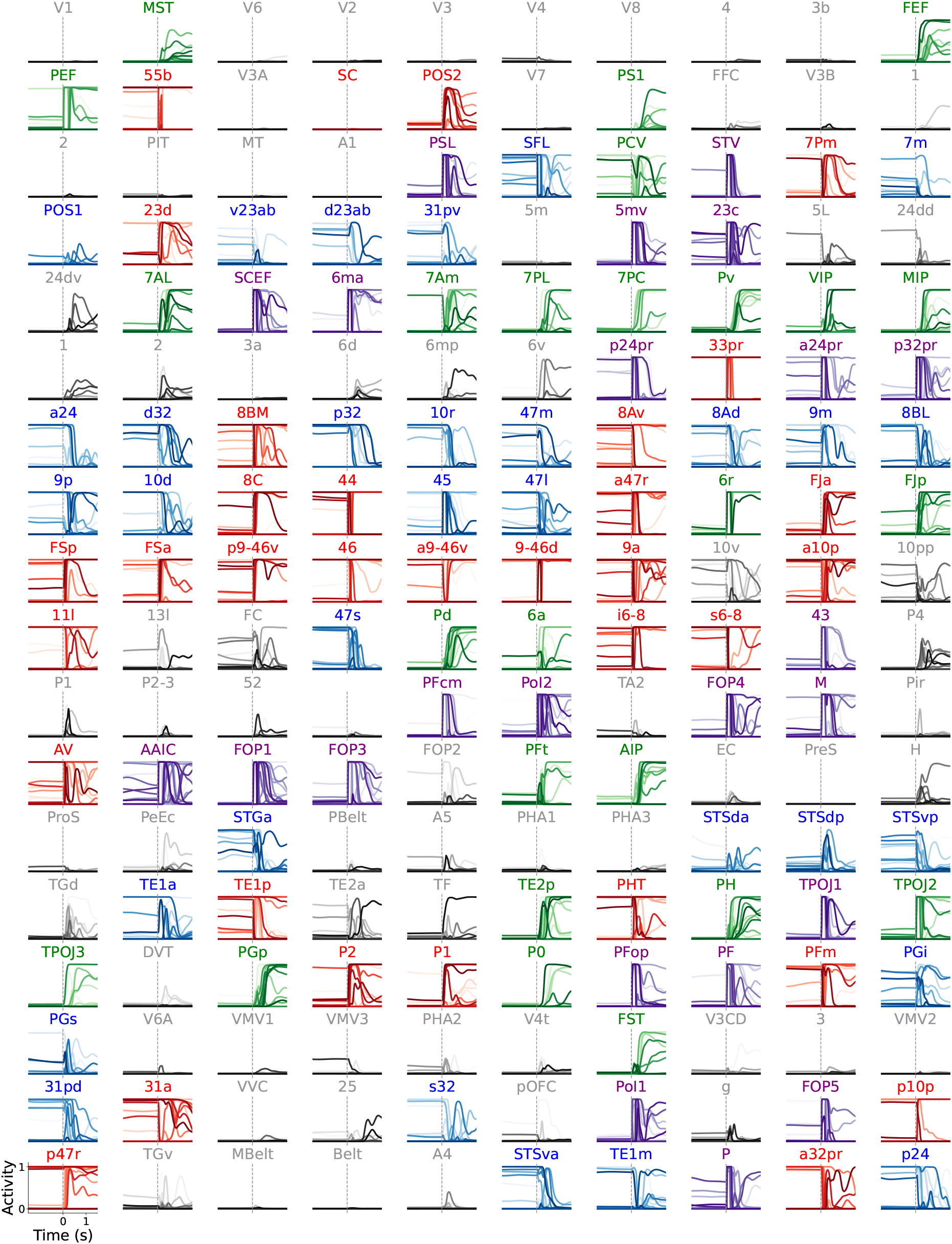
Supplementary figure corresponding to Fig. 6c in the main text. Network response to a 150ms stimulus to areas in the salience network. Neural dynamics for 15 representative units. Areas with purple labels and traces belong to the salience network. Areas with red labels and traces belong to the frontoparietal network. Areas with green labels and traces belong to the DAN. Areas with blue labels and traces belong to the DAN.

**Fig. S21:**
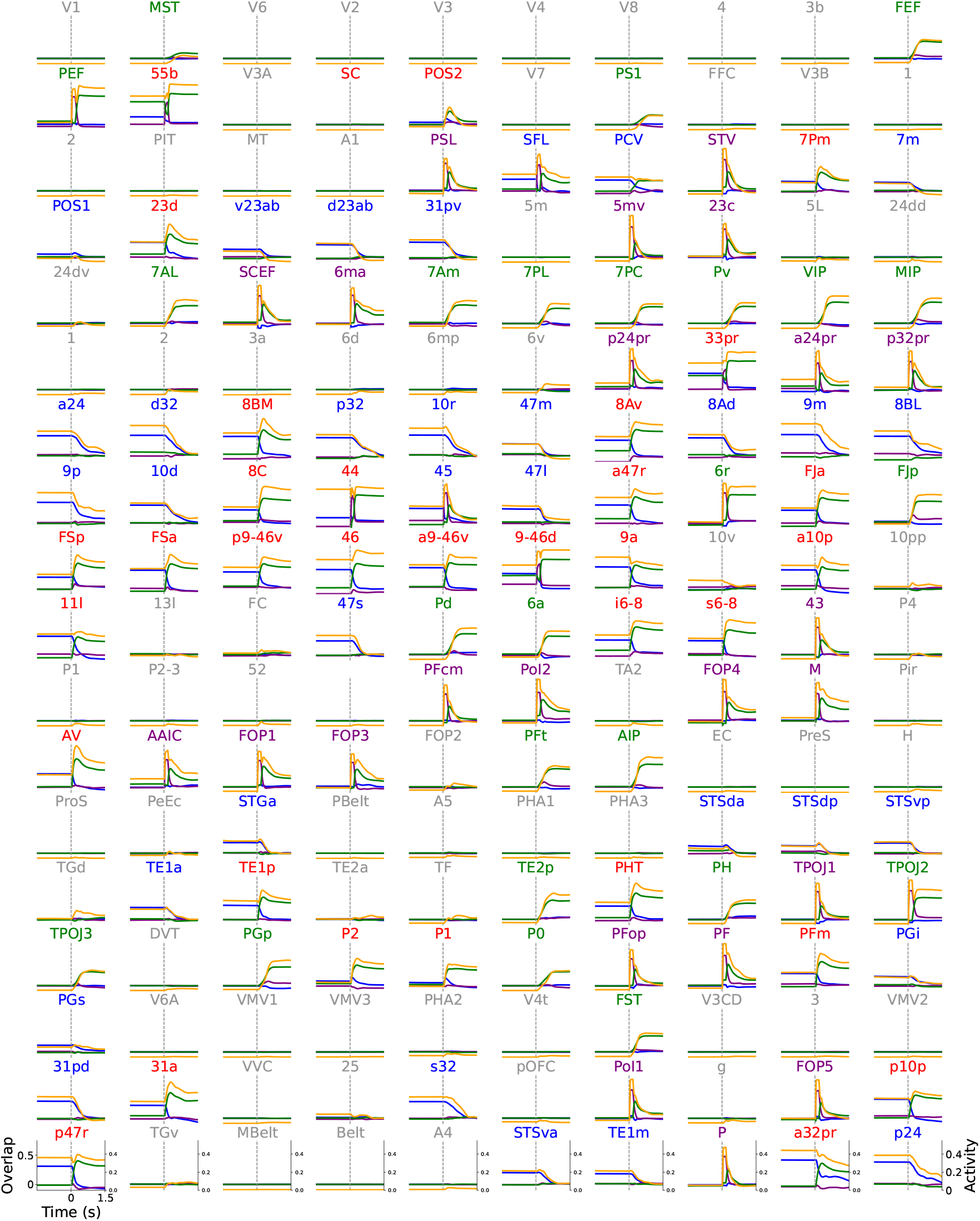
Supplementary figure corresponding to Fig. 6c, j-o in the main text. Network response to a 150ms stimulus to areas in the salience network. Blue: overlap FP-DMN attractor; Green: overlap FP-DAN attractor; Purple: overlap with the pattern stimulated in the salience network. The gray dashed line marks the transient stimulation of salience areas at 0s for 150 ms. Purple labels and traces: Salience network. Red: Frontoparietal network. Green: Dorsal Attention Network (DAN). Blue: Visual network.

## References

[1] Jonathan D Power, Alexander L Cohen, Steven M Nelson, Gagan S Wig, Kelly Anne Barnes, Jessica A Church, Alecia C Vogel, Timothy O Laumann, Fran M Miezin, Bradley L Schlaggar, et al. Functional network organization of the human brain. Neuron, 72(4):665–678, 2011.

[2] Matthew L Leavitt, Diego Mendoza-Halliday, and Julio C Martinez-Trujillo. Sustained activity encoding working memories: not fully distributed. Trends in Neurosciences, 40(6):328–346, 2017.

[3] Nicholas A Steinmetz, Peter Zatka-Haas, Matteo Carandini, and Kenneth D Harris. Distributed coding of choice, action and engagement across the mouse brain. Nature, 576(7786):266–273, 2019.

[4] Susu Chen, Yi Liu, Ziyue Aiden Wang, Jennifer Colonell, Liu D Liu, Han Hou, Nai-Wen Tien, Tim Wang, Timothy Harris, Shaul Druckmann, et al. Brain-wide neural activity underlying memory-guided movement. Cell, 187(3):676–691, 2024.

[5] Lucas Pinto, Kanaka Rajan, Brian DePasquale, Stephan Y Thiberge, David W Tank, and Carlos D Brody. Task-dependent changes in the large-scale dynamics and necessity of cortical regions. Neuron, 104(4):810–824, 2019.

[6] Steven L Bressler and Vinod Menon. Large-scale brain networks in cognition: emerging methods and principles. Trends in cognitive sciences, 14(6):277–290, 2010.

[7] Steven E Petersen and Olaf Sporns. Brain networks and cognitive architectures. Neuron, 88(1):207–219, 2015.

[8] Maurizio Corbetta and Gordon L Shulman. Control of goal-directed and stimulus-driven attention in the brain. Nature reviews neuroscience, 3(3):201–215, 2002.

[9] Michael D Fox, Maurizio Corbetta, Abraham Z Snyder, Justin L Vincent, and Marcus E Raichle. Spontaneous neuronal activity distinguishes human dorsal and ventral attention systems. Proceedings of the National Academy of Sciences, 103(26):10046–10051, 2006.

[10] Simone Vossel, Joy J Geng, and Gereon R Fink. Dorsal and ventral attention systems: distinct neural circuits but collaborative roles. The Neuroscientist, 20(2):150–159, 2014.

[11] Marcus E Raichle, Ann Mary MacLeod, Abraham Z Snyder, William J Powers, Debra A Gusnard, and Gordon L Shulman. A default mode of brain function. Proceedings of the national academy of sciences, 98(2):676–682, 2001.

[12] Justin L Vincent, Gaurav H Patel, Michael D Fox, Abraham Z Snyder, Justin T Baker, David C Van Essen, John M Zempel, Lawrence H Snyder, Maurizio Corbetta, and Marcus E Raichle. Intrinsic functional architecture in the anaesthetized monkey brain. Nature, 447(7140):83–86, 2007.

[13] Annabelle M Belcher, Cecil C Yen, Haley Stepp, Hong Gu, Hanbing Lu, Yihong Yang, Afonso C Silva, and Elliot A Stein. Large-scale brain networks in the awake, truly resting marmoset monkey. Journal of neuroscience, 33(42):16796–16804, 2013.

[14] Sarah K Barks, Lisa A Parr, and James K Rilling. The default mode network in chimpanzees (pan troglodytes) is similar to that of humans. Cerebral cortex, 25(2):538–544, 2015.

[15] Benjamin Y Hayden, David V Smith, and Michael L Platt. Electrophysiological correlates of default-mode processing in macaque posterior cingulate cortex. Proceedings of the National Academy of Sciences, 106(14):5948–5953, 2009.

[16] BT Thomas Yeo, Fenna M Krienen, Jorge Sepulcre, Mert R Sabuncu, Danial Lashkari, Marisa Hollinshead, Joshua L Roffman, Jordan W Smoller, Lilla Zöllei, Jonathan R Polimeni, et al. The organization of the human cerebral cortex estimated by intrinsic functional connectivity. Journal of neurophysiology, 2011.

[17] Devarajan Sridharan, Daniel J Levitin, and Vinod Menon. A critical role for the right fronto-insular cortex in switching between central-executive and default-mode networks. Proceedings of the National Academy of Sciences, 105(34):12569–12574, 2008.

[18] William W Seeley. The salience network: a neural system for perceiving and responding to homeostatic demands. Journal of Neuroscience, 39(50):9878–9882, 2019.

[19] John Duncan. The multiple-demand (md) system of the primate brain: mental programs for intelligent behaviour. Trends in cognitive sciences, 14(4):172–179, 2010.

[20] Theodore P Zanto and Adam Gazzaley. Fronto-parietal network: flexible hub of cognitive control. Trends in cognitive sciences, 17(12):602–603, 2013.

[21] Scott Marek and Nico UF Dosenbach. The frontoparietal network: function, electrophysiology, and importance of individual precision mapping. Dialogues in clinical neuroscience, 20(2):133–140, 2018.

[22] R Nathan Spreng, W Dale Stevens, Jon P Chamberlain, Adrian W Gilmore, and Daniel L Schacter. Default network activity, coupled with the frontoparietal control network, supports goal-directed cognition. Neuroimage, 53(1):303–317, 2010.

[23] Moataz Assem, Matthew F Glasser, David C Van Essen, and John Duncan. A domain-general cognitive core defined in multimodally parcellated human cortex. Cerebral Cortex, 30(8):4361–4380, 2020.

[24] Matthew L Dixon, Alejandro De La Vega, Caitlin Mills, Jessica Andrews-Hanna, R Nathan Spreng, Michael W Cole, and Kalina Christoff. Heterogeneity within the frontoparietal control network and its relationship to the default and dorsal attention networks. Proceedings of the National Academy of Sciences, 115(7):E1598–E1607, 2018.

[25] Rodrigo M Braga and Randy L Buckner. Parallel interdigitated distributed networks within the individual estimated by intrinsic functional connectivity. Neuron, 95(2):457–471, 2017.

[26] Jingnan Du, Lauren M DiNicola, Peter A Angeli, Noam Saadon-Grosman, Wendy Sun, Stephanie Kaiser, Joanna Ladopoulou, Aihuiping Xue, BT Thomas Yeo, Mark C Eldaief, et al. Organization of the human cerebral cortex estimated within individuals: networks, global topography, and function. Journal of Neurophysiology, 131(6):1014–1082, 2024.

[27] LF Abbott. Where are the switches on this thing. 23 Problems in systems neuroscience, pages 423–431, 2006.

[28] João D Semedo, Amin Zandvakili, Christian K Machens, M Yu Byron, and Adam Kohn. Cortical areas interact through a communication subspace. Neuron, 102(1):249–259, 2019.

[29] Adam Kohn, Anna I Jasper, João D Semedo, Evren Gokcen, Christian K Machens, and M Yu Byron. Principles of corticocortical communication: proposed schemes and design considerations. Trends in Neurosciences, 43(9):725–737, 2020.

[30] Camden J MacDowell, Alexandra Libby, Caroline I Jahn, Sina Tafazoli, and Timothy J Buschman. Multiplexed subspaces route neural activity across brain-wide networks. bioRxiv, pages 2023–02, 2023.

[31] Ramakrishnan Iyer, Joshua H Siegle, Gayathri Mahalingam, Shawn Olsen, and Stefan Mihalas. Geometry of inter-areal interactions in mouse visual cortex. bioRxiv, pages 2021–06, 2021.

[32] Xiao-Jing Wang. Theory of the multiregional neocortex: large-scale neural dynamics and distributed cognition. Annual review of neuroscience, 45(1):533–560, 2022.

[33] Gustavo Deco, Viktor Jirsa, Anthony R McIntosh, Olaf Sporns, and Rolf Kötter. Key role of coupling, delay, and noise in resting brain fluctuations. Proceedings of the National Academy of Sciences, 106(25):10302–10307, 2009.

[34] Enrique CA Hansen, Demian Battaglia, Andreas Spiegler, Gustavo Deco, and Viktor K Jirsa. Functional connectivity dynamics: modeling the switching behavior of the resting state. Neuroimage, 105:525–535, 2015.

[35] Rishidev Chaudhuri, Kenneth Knoblauch, Marie-Alice Gariel, Henry Kennedy, and Xiao-Jing Wang. A large-scale circuit mechanism for hierarchical dynamical processing in the primate cortex. Neuron, 88(2):419–431, 2015.

[36] Murat Demirtaş, Joshua B Burt, Markus Helmer, Jie Lisa Ji, Brendan D Adkinson, Matthew F Glasser, David C Van Essen, Stamatios N Sotiropoulos, Alan Anticevic, and John D Murray. Hierarchical heterogeneity across human cortex shapes large-scale neural dynamics. Neuron, 101(6):1181–1194, 2019.

[37] Joana Cabral, Etienne Hugues, Olaf Sporns, and Gustavo Deco. Role of local network oscillations in resting-state functional connectivity. Neuroimage, 57(1):130–139, 2011.

[38] Xiaolu Kong, Ru Kong, Csaba Orban, Peng Wang, Shaoshi Zhang, Kevin Anderson, Avram Holmes, John D Murray, Gustavo Deco, Martijn van den Heuvel, et al. Sensory-motor cortices shape functional connectivity dynamics in the human brain. Nature communications, 12(1):6373, 2021.

[39] Eli J Muller, Brandon Munn, Giulia Baracchini, Ben D Fulcher, Vicente Medel, Michelle J Redinbaugh, Yuri B Saalmann, Bingni Wen Brunton, Steve L Brunton, and James Shine. Thalamic control over laminar cortical dynamics across conscious states. bioRxiv, pages 2024–07, 2024.

[40] Christopher J Honey, Rolf Kötter, Michael Breakspear, and Olaf Sporns. Network structure of cerebral cortex shapes functional connectivity on multiple time scales. Proceedings of the National Academy of Sciences, 104(24):10240–10245, 2007.

[41] Jorge F Mejias and Xiao-Jing Wang. Mechanisms of distributed working memory in a large-scale network of macaque neocortex. Elife, 11:e72136, 2022.

[42] Sean Froudist-Walsh, Daniel P Bliss, Xingyu Ding, Lucija Rapan, Meiqi Niu, Kenneth Knoblauch, Karl Zilles, Henry Kennedy, Nicola Palomero-Gallagher, and Xiao-Jing Wang. A dopamine gradient controls access to distributed working memory in the large-scale monkey cortex. Neuron, 109(21):3500–3520, 2021.

[43] Xingyu Ding, Sean Froudist-Walsh, Jorge Jaramillo, Junjie Jiang, and Xiao-Jing Wang. Cell type-specific connectome predicts distributed working memory activity in the mouse brain. Elife, 13:e85442, 2024.

[44] Mengli Feng, Abhirup Bandyopadhyay, and Jorge F Mejias. Emergence of distributed working memory in a human brain network model. bioRxiv, pages 2023–01, 2023.

[45] Michael Schirner, Gustavo Deco, and Petra Ritter. Learning how network structure shapes decision-making for bio-inspired computing. Nature Communications, 14(1):2963, 2023.

[46] Licheng Zou, Nicola Palomero-Gallagher, Douglas Zhou, Songting Li, and Jorge F Mejias. Distributed evidence accumulation across macaque large-scale neocortical networks during perceptual decision making. bioRxiv, pages 2023–12, 2023.

[47] Ulysse Klatzmann, Sean Froudist-Walsh, Daniel P Bliss, Panagiota Theodoni, Jorge Mejías, Meiqi Niu, Lucija Rapan, Nicola Palomero-Gallagher, Claire Sergent, Stanislas Dehaene, et al. A connectome-based model of conscious access in monkey cortex. BioRxiv, pages 2022–02, 2022.

[48] David Sussillo and Omri Barak. Opening the black box: low-dimensional dynamics in high-dimensional recurrent neural networks. Neural computation, 25(3):626–649, 2013.

[49] John D Murray, Alberto Bernacchia, Nicholas A Roy, Christos Constantinidis, Ranulfo Romo, and Xiao-Jing Wang. Stable population coding for working memory coexists with heterogeneous neural dynamics in prefrontal cortex. Proceedings of the National Academy of Sciences, 114(2):394–399, 2017.

[50] Christos Constantinidis, Shintaro Funahashi, Daeyeol Lee, John D Murray, Xue-Lian Qi, Min Wang, and Amy FT Arnsten. Persistent spiking activity underlies working memory. Journal of neuroscience, 38(32):7020–7028, 2018.

[51] Mark G Stokes, Makoto Kusunoki, Natasha Sigala, Hamed Nili, David Gaffan, and John Duncan. Dynamic coding for cognitive control in prefrontal cortex. Neuron, 78(2):364–375, 2013.

[52] Sean E Cavanagh, John P Towers, Joni D Wallis, Laurence T Hunt, and Steven W Kennerley. Reconciling persistent and dynamic hypotheses of working memory coding in prefrontal cortex. Nature communications, 9(1):3498, 2018.

[53] Eelke Spaak, Kei Watanabe, Shintaro Funahashi, and Mark G Stokes. Stable and dynamic coding for working memory in primate prefrontal cortex. Journal of neuroscience, 37(27):6503–6516, 2017.

[54] Ethan M Meyers, David J Freedman, Gabriel Kreiman, Earl K Miller, and Tomaso Poggio. Dynamic population coding of category information in inferior temporal and prefrontal cortex. Journal of neurophysiology, 100(3):1407–1419, 2008.

[55] Jake P Stroud, John Duncan, and Máté Lengyel. The computational foundations of dynamic coding in working memory. Trends in Cognitive Sciences, 2024.

[56] Xiao-Jing Wang. 50 years of mnemonic persistent activity: quo vadis? Trends in Neurosciences, 44(11):888–902, 2021.

[57] Christopher D Harvey, Philip Coen, and David W Tank. Choice-specific sequences in parietal cortex during a virtual-navigation decision task. Nature, 484(7392):62–68, 2012.

[58] Kayvon Daie, Lorenzo Fontolan, Shaul Druckmann, and Karel Svoboda. Feedforward amplification in recurrent networks underlies paradoxical neural coding. bioRxiv, pages 2023–08, 2023.

[59] Alexandra Busch, Megan Roussy, Rogelio Luna, Matthew L Leavitt, Maryam H Mofrad, Roberto A Gulli, Benjamin Corrigan, Ján Mináč, Adam J Sachs, Lena Palaniyappan, et al. Neuronal activation sequences in lateral prefrontal cortex encode visuospatial working memory during virtual navigation. Nature Communications, 15(1):4471, 2024.

[60] S-I Amari. Learning patterns and pattern sequences by self-organizing nets of threshold elements. IEEE Transactions on computers, 100(11):1197–1206, 1972.

[61] John J Hopfield. Neural networks and physical systems with emergent collective computational abilities. Proceedings of the national academy of sciences, 79(8):2554–2558, 1982.

[62] Daniel J Amit, Hanoch Gutfreund, and Haim Sompolinsky. Storing infinite numbers of patterns in a spin-glass model of neural networks. Physical Review Letters, 55(14):1530, 1985.

[63] David Kleinfeld. Sequential state generation by model neural networks. Proceedings of the National Academy of Sciences, 83(24):9469–9473, 1986.

[64] Haim Sompolinsky and Ido Kanter. Temporal association in asymmetric neural networks. Physical review letters, 57(22):2861, 1986.

[65] Maxwell Gillett, Ulises Pereira, and Nicolas Brunel. Characteristics of sequential activity in networks with temporally asymmetric hebbian learning. Proceedings of the National Academy of Sciences, 117(47):29948–29958, 2020.

[66] Haim Sompolinsky, Andrea Crisanti, and Hans-Jurgen Sommers. Chaos in random neural networks. Physical review letters, 61(3):259, 1988.

[67] Xiao-Jing Wang. Macroscopic gradients of synaptic excitation and inhibition in the neocortex. Nature reviews neuroscience, 21(3):169–178, 2020.

[68] Elizabeth Herbert and Srdjan Ostojic. The impact of sparsity in low-rank recurrent neural networks. PLoS Computational Biology, 18(8):e1010426, 2022.

[69] Chad J Donahue, Stamatios N Sotiropoulos, Saad Jbabdi, Moises Hernandez-Fernandez, Timothy E Behrens, Tim B Dyrby, Timothy Coalson, Henry Kennedy, Kenneth Knoblauch, David C Van Essen, et al. Using diffusion tractography to predict cortical connection strength and distance: a quantitative comparison with tracers in the monkey. Journal of Neuroscience, 36(25):6758–6770, 2016.

[70] Daniel J Amit and Nicolas Brunel. Model of global spontaneous activity and local structured activity during delay periods in the cerebral cortex. Cerebral cortex (New York, NY: 1991), 7(3):237–252, 1997.

[71] A Herz, B Sulzer, R Kühn, and JL Van Hemmen. Hebbian learning reconsidered: Representation of static and dynamic objects in associative neural nets. Biological cybernetics, 60:457–467, 1989.

[72] Olivia L White, Daniel D Lee, and Haim Sompolinsky. Short-term memory in orthogonal neural networks. Physical review letters, 92(14):148102, 2004.

[73] Surya Ganguli, Dongsung Huh, and Haim Sompolinsky. Memory traces in dynamical systems. Proceedings of the national academy of sciences, 105(48):18970–18975, 2008.

[74] Mark S Goldman. Memory without feedback in a neural network. Neuron, 61(4):621–634, 2009.

[75] Maxwell Gillett and Nicolas Brunel. Dynamic control of sequential retrieval speed in networks with heterogeneous learning rules. bioRxiv, pages 2023–03, 2023.

[76] Stephen Grossberg. On learning, information, lateral inhibition, and transmitters. Mathematical Biosciences, 4(3-4):255–310, 1969.

[77] Ting Xu, Karl-Heinz Nenning, Ernst Schwartz, Seok-Jun Hong, Joshua T Vogelstein, Alexandros Goulas, Damien A Fair, Charles E Schroeder, Daniel S Margulies, Jonny Smallwood, et al. Cross-species functional alignment reveals evolutionary hierarchy within the connectome. Neuroimage, 223:117346, 2020.

[78] Sean Froudist-Walsh, Ting Xu, Meiqi Niu, Lucija Rapan, Ling Zhao, Daniel S Margulies, Karl Zilles, Xiao-Jing Wang, and Nicola Palomero-Gallagher. Gradients of neurotransmitter receptor expression in the macaque cortex. Nature neuroscience, 26(7):1281–1294, 2023.

[79] Michael N Shadlen and William T Newsome. Neural basis of a perceptual decision in the parietal cortex (area lip) of the rhesus monkey. Journal of neurophysiology, 86(4):1916–1936, 2001.

[80] Daniel J Amit. Modeling brain function: The world of attractor neural networks. Cambridge university press, 1989.

[81] B Tirozzi and M Tsodyks. Chaos in highly diluted neural networks. Europhysics Letters, 14(8):727, 1991.

[82] Ulises Pereira-Obilinovic, Johnatan Aljadeff, and Nicolas Brunel. Forgetting leads to chaos in attractor networks. Physical Review X, 13(1):011009, 2023.

[83] Francesca Mastrogiuseppe and Srdjan Ostojic. Linking connectivity, dynamics, and computations in low-rank recurrent neural networks. Neuron, 99(3):609–623, 2018.

[84] Marcus E Raichle. The brain’s default mode network. Annual review of neuroscience, 38(1):433–447, 2015.

[85] Michael D Greicius, Ben Krasnow, Allan L Reiss, and Vinod Menon. Functional connectivity in the resting brain: a network analysis of the default mode hypothesis. Proceedings of the national academy of sciences, 100(1):253–258, 2003.

[86] AM Clare Kelly, Lucina Q Uddin, Bharat B Biswal, F Xavier Castellanos, and Michael P Milham. Competition between functional brain networks mediates behavioral variability. Neuroimage, 39(1):527–537, 2008.

[87] Alan Anticevic, Michael W Cole, John D Murray, Philip R Corlett, Xiao-Jing Wang, and John H Krystal. The role of default network deactivation in cognition and disease. Trends in cognitive sciences, 16(12):584–592, 2012.

[88] Andrew C Murphy, Maxwell A Bertolero, Lia Papadopoulos, David M Lydon-Staley, and Danielle S Bassett. Multimodal network dynamics underpinning working memory. Nature communications, 11(1):3035, 2020.

[89] Aihuiping Xue, Ru Kong, Qing Yang, Mark C Eldaief, Peter A Angeli, Lauren M DiNicola, Rodrigo M Braga, Randy L Buckner, and BT Thomas Yeo. The detailed organization of the human cerebellum estimated by intrinsic functional connectivity within the individual. Journal of neurophysiology, 125(2):358–384, 2021.

[90] Daniel J Mitchell, Andrew H Bell, Mark J Buckley, Anna S Mitchell, Jerome Sallet, and John Duncan. A putative multiple-demand system in the macaque brain. Journal of Neuroscience, 36(33):8574–8585, 2016.

[91] Philippe Domenech, Jérôme Redouté, Etienne Koechlin, and Jean-Claude Dreher. The neuro-computational architecture of value-based selection in the human brain. Cerebral Cortex, 28(2):585–601, 2018.

[92] Hidehiko K Inagaki, Lorenzo Fontolan, Sandro Romani, and Karel Svoboda. Discrete attractor dynamics underlies persistent activity in the frontal cortex. Nature, 566(7743):212–217, 2019.

[93] Joaquin M Fuster and Garrett E Alexander. Neuron activity related to short-term memory. Science, 173(3997):652–654, 1971.

[94] Shintaro Funahashi, Charles J Bruce, and Patricia S Goldman-Rakic. Mnemonic coding of visual space in the monkey’s dorsolateral prefrontal cortex. Journal of neurophysiology, 61(2):331–349, 1989.

[95] Mikael Lundqvist, Pawel Herman, and Earl K Miller. Working memory: delay activity, yes! persistent activity? maybe not. Journal of neuroscience, 38(32):7013–7019, 2018.

[96] Jonathan Kadmon and Haim Sompolinsky. Transition to chaos in random neuronal networks. Physical Review X, 5(4):041030, 2015.

[97] Laurent Itti and Christof Koch. A saliency-based search mechanism for overt and covert shifts of visual attention. Vision research, 40(10-12):1489–1506, 2000.

[98] Joao Barbosa, Rémi Proville, Chris C Rodgers, Michael R DeWeese, Srdjan Ostojic, and Yves Boubenec. Early selection of task-relevant features through population gating. Nature communications, 14(1):6837, 2023.

[99] Guangyu Robert Yang, John D Murray, and Xiao-Jing Wang. A dendritic disinhibitory circuit mechanism for pathway-specific gating. Nature communications, 7(1):12815, 2016.

[100] Martin Vinck, Cem Uran, Georgios Spyropoulos, Irene Onorato, Ana Clara Broggini, Marius Schneider, and Andres Canales-Johnson. Principles of large-scale neural interactions. Neuron, 111(7):987–1002, 2023.

[101] Gyorgy Buzsaki. Rhythms of the Brain. Oxford university press, 2006.

[102] Francisco Varela, Jean-Philippe Lachaux, Eugenio Rodriguez, and Jacques Martinerie. The brainweb: phase synchronization and large-scale integration. Nature reviews neuroscience, 2(4):229–239, 2001.

[103] Steven L Bressler and JA Scott Kelso. Cortical coordination dynamics and cognition. Trends in cognitive sciences, 5(1):26–36, 2001.

[104] John E Lisman and Ole Jensen. The theta-gamma neural code. Neuron, 77(6):1002–1016, 2013.

[105] Pascal Fries. A mechanism for cognitive dynamics: neuronal communication through neuronal coherence. Trends in cognitive sciences, 9(10):474–480, 2005.

[106] Pascal Fries. Rhythms for cognition: communication through coherence. Neuron, 88(1):220–235, 2015.

[107] Wolf Singer. Neuronal synchrony: a versatile code for the definition of relations? Neuron, 24(1):49–65, 1999.

[108] Wolf Singer and Charles M Gray. Visual feature integration and the temporal correlation hypothesis. 2003.

[109] N Kopell, GB Ermentrout, Miles A Whittington, and Roger D Traub. Gamma rhythms and beta rhythms have different synchronization properties. Proceedings of the National Academy of Sciences, 97(4):1867–1872, 2000.

[110] Pieter R Roelfsema. Solving the binding problem: Assemblies form when neurons enhance their firing rate—they don’t need to oscillate or synchronize. Neuron, 111(7):1003–1019, 2023.

[111] Carsen Stringer, Marius Pachitariu, Nicholas Steinmetz, Charu Bai Reddy, Matteo Carandini, and Kenneth D Harris. Spontaneous behaviors drive multidimensional, brainwide activity. Science, 364(6437):eaav7893, 2019.

[112] Jason Manley, Sihao Lu, Kevin Barber, Jeffrey Demas, Hyewon Kim, David Meyer, Francisca Martínez Traub, and Alipasha Vaziri. Simultaneous, cortex-wide dynamics of up to 1 million neurons reveal unbounded scaling of dimensionality with neuron number. Neuron, 112(10):1694–1709, 2024.

[113] Johnatan Aljadeff, Merav Stern, and Tatyana Sharpee. Transition to chaos in random networks with cell-type-specific connectivity. Physical review letters, 114(8):088101, 2015.

[114] David G Clark and Manuel Beiran. Structure of activity in multiregion recurrent neural networks. arXiv preprint arXiv:2402.12188, 2024.

[115] Łukasz Kuśmierz, Ulises Pereira-Obilinovic, Zhixin Lu, Dana Mastrovito, and Stefan Mihalas. Hierarchy of chaotic dynamics in random modular networks. arXiv preprint arXiv:2410.06361, 2024.

[116] Alexis M Dubreuil and Nicolas Brunel. Storing structured sparse memories in a multi-modular cortical network model. Journal of computational neuroscience, 40:157–175, 2016.

[117] Matthew G Perich, Charlotte Arlt, Sofia Soares, Megan E Young, Clayton P Mosher, Juri Minxha, Eugene Carter, Ueli Rutishauser, Peter H Rudebeck, Christopher D Harvey, et al. Inferring brain-wide interactions using data-constrained recurrent neural network models. BioRxiv, pages 2020–12, 2020.

[118] Ian Goodfellow, Yoshua Bengio, and Aaron Courville. Deep learning. MIT press, 2016.

[119] David C Van Essen, John Smith, Matthew F Glasser, Jennifer Elam, Chad J Donahue, Donna L Dierker, Erin K Reid, Timothy Coalson, and John Harwell. The brain analysis library of spatial maps and atlases (balsa) database. Neuroimage, 144:270–274, 2017.

[120] Timothy EJ Behrens, H Johansen Berg, Saad Jbabdi, Matthew FS Rushworth, and Mark W Woolrich. Probabilistic diffusion tractography with multiple fibre orientations: What can we gain? neuroimage, 34(1):144–155, 2007.

[121] Matthew F Glasser, Stamatios N Sotiropoulos, J Anthony Wilson, Timothy S Coalson, Bruce Fischl, Jesper L Andersson, Junqian Xu, Saad Jbabdi, Matthew Webster, Jonathan R Polimeni, et al. The minimal preprocessing pipelines for the human connectome project. Neuroimage, 80:105–124, 2013.

[122] Nikola T Markov, MM Ercsey-Ravasz, AR Ribeiro Gomes, Camille Lamy, Loic Magrou, Julien Vezoli, P Misery, A Falchier, R Quilodran, MA Gariel, et al. A weighted and directed interareal connectivity matrix for macaque cerebral cortex. Cerebral cortex, 24(1):17–36, 2014.

[123] Matthew F Glasser, Timothy S Coalson, Emma C Robinson, Carl D Hacker, John Harwell, Essa Yacoub, Kamil Ugurbil, Jesper Andersson, Christian F Beckmann, Mark Jenkinson, et al. A multi-modal parcellation of human cerebral cortex. Nature, 536(7615):171–178, 2016.

[124] Ronald R Coifman and Stéphane Lafon. Diffusion maps. Applied and computational harmonic analysis, 21(1):5–30, 2006.

[125] Daniel S Margulies, Satrajit S Ghosh, Alexandros Goulas, Marcel Falkiewicz, Julia M Huntenburg, Georg Langs, Gleb Bezgin, Simon B Eickhoff, F Xavier Castellanos, Michael Petrides, et al. Situating the default-mode network along a principal gradient of macroscale cortical organization. Proceedings of the National Academy of Sciences, 113(44):12574–12579, 2016.

[126] Emma C Robinson, Kara Garcia, Matthew F Glasser, Zhengdao Chen, Timothy S Coalson, Antonios Makropoulos, Jelena Bozek, Robert Wright, Andreas Schuh, Matthew Webster, et al. Multimodal surface matching with higher-order smoothness constraints. Neuroimage, 167:453–465, 2018.

[127] Guy N Elston. Specialization of the neocortical pyramidal cell during primate evolution. Evolution of nervous systems, pages 191–242, 2007.

[128] Joshua B Burt, Murat Demirtaş, William J Eckner, Natasha M Navejar, Jie Lisa Ji, William J Martin, Alberto Bernacchia, Alan Anticevic, and John D Murray. Hierarchy of transcriptomic specialization across human cortex captured by structural neuroimaging topography. Nature neuroscience, 21(9):1251–1259, 2018.

[129] Matthew Farrell and Cengiz Pehlevan. Recall tempo of hebbian sequences depends on the interplay of hebbian kernel with tutor signal timing. Proceedings of the National Academy of Sciences, 121(32):e2309876121, 2024.

[130] David C Van Essen, Stephen M Smith, Deanna M Barch, Timothy EJ Behrens, Essa Yacoub, Kamil Ugurbil, Wu-Minn HCP Consortium, et al. The wu-minn human connectome project: an overview. Neuroimage, 80:62–79, 2013.

